# High-resolution transcriptional atlas of growing maize shoot organs throughout plant development under well-watered and drought conditions

**DOI:** 10.1101/2025.03.12.642568

**Authors:** Jie Zhang, Lennart Verbraeken, Heike Sprenger, Stien Mertens, Nathalie Wuyts, Bernard Cannoot, Jolien De Block, Kirin Demuynck, Annelore Natran, Katrien Maleux, Julie Merchie, Steven Crafts-Brandner, Jonathan Vogel, Wesley Bruce, Dirk Inzé, Steven Maere, Hilde Nelissen

## Abstract

Crop improvement goals for maize (*Zea mays* L.) involve the targeted optimization of various organs, making it crucial to understand the developmental characteristics and gene expression patterns during organ development and in response to environmental stresses such as drought. In this study, we investigated the development of maize leaves and internodes at both macroscopic and cellular level, and identified a shared fundamental growth design with distinct timing between the two organs. By transcriptome profiling developmental zones of leaves and internodes of different ranks, and of the ear, at different growth stages under both well-watered and drought conditions, we generated a high-resolution spatiotemporal transcriptome dataset on 272 different tissues and conditions, which we make available as a searchable database. While the gene regulatory networks governing cell division and cell elongation were highly conserved across organs, precise expression regulation of particular gene families was observed across organs and within the same organ. Additionally, we highlight the expression of key genes involved in regulating leaf angle and vascular development, showing spatiotemporal regulation of differentiation parallel to growth. This comprehensive expression atlas, combined with phenotypic data, offers a deeper understanding of the similarities and differences among shoot organs and tissues during development and drought response, and provides a valuable resource for engineering organ-specific traits in maize.

## Introduction

Improving crop productivity to produce more food and feed in an uncertain context due to climate change remains an ongoing challenge (Ortiz-Bobea et al., 2021). One tool to improve crop productivity is the idea of crop ideotypes, the conceptualization of a crop phenotype ideal for a particular environment (Donald, 1968). The environment, including management practices, plays a key role in this concept as there is not a single crop ideotype for each species, but many depending on the target circumstances (Frey, 1971).

For maize, one of the world’s most important cereal crops, proposed ideotypes advocate for upright top leaves to have the most efficient radiation interception at high planting densities (Loomis and Williams, 1969; Mock and Pearce, 1975; Donald and Hamblin, 1976; Gong et al., 2015), and more horizontal lower leaves to ensure maximal light interception at early growth stages (Gong et al., 2015). Breeding for optimized leaf angle has long been a key focus in yield improvement. A series of genes that control maize leaf angle have been identified, such as *ZmLIGULELESS1* (*ZmLG1*) and *ZmLIGULELESS2* (*ZmLG2*) which have a crucial role in ligule development at the blade-sheath boundary. Both *lg1* and *lg2* null alleles have an upright architecture with a small leaf angle (Becraft et al., 1990; Walsh et al., 1998). Brassinosteroids (BRs) have also been found to play an important role in leaf angle formation. Mutants deficient in BR biosynthesis or signaling exhibit more upright leaves due to alternations in cell division and cell elongation (Cao et al., 2022). In addition to optimizing leaf traits, breeding for a shorter, yet sturdy, stem can improve the harvest index and the resistance to lodging (Donald and Hamblin, 1976; Gong et al., 2015). Furthermore, achieving a greater yield also involves selecting certain traits of the ear, as components of ear inflorescence architecture such as kernel row number, number of kernels per row, and kernel number density are positively correlated with grain yield (Khakwani et al., 2018). Improving these traits requires affecting growth of various shoot organs in distinct ways.

Maize leaves are used as a model for organ growth, as their large size and the almost one-dimensional nature of their growth (i.e. almost exclusively growth along the proximodistal axis) greatly facilitates quantifying and analyzing their growth (Fournier et al., 2005; Nelissen et al., 2013; Avramova et al., 2015). Maize leaves also provide a convenient model at the cellular level due to the linear organization of their growth. Maize leaves contain a basal growth zone, consisting of a division zone and an elongation zone. The division zone contains dividing cells that gradually move on to the more distal elongation zone, where they stop dividing and gradually expand along the proximodistal axis until they reach their mature lengths (Nelissen et al., 2013). This interplay between cell division and elongation results in a sigmoidal growth pattern at the macroscopic level, where the elongation of an individual leaf is characterized by subsequent exponential, steady-state and asymptotic stages (Voorend et al., 2014). In contrast to maize leaves, growing internodes (the parts of the maize stem in between leaf nodes) are not used as a model or easily studied. Internodes grow while concealed by encircling leaf sheaths. Several methods have been used to study them in the past, including dissection (Robertson, 1994), slicing an opening in the surrounding leaf sheaths (Morrison et al., 1994), X-ray photography, or by proxy via the collars of the leaves connected to the internode (Fournier and Andrieu, 2000). Like leaves, internode elongation follows a sigmoidal growth pattern, with a sequential exponential, steady-state and asymptotic phase (Morrison et al., 1994; Fournier and Andrieu, 2000). Also similar to leaves, internode elongation is driven by cell production at the basal intercalary meristem, followed by cell elongation higher in the internode (Morrison et al., 1994). Although similar growth patterns have been observed in leaves and internodes, it remains unclear whether the spatiotemporal characteristics of these growth zones and the molecular mechanism governing the growth processes are alike in both organs.

The regulation of growth involves a complex interplay of internal and external factors, with transcription factors playing a central role in modulating gene expression associated with cell proliferation and expansion. Among these, the GROWTH-REGULATING FACTOR (GRF) family has been recognized for its role in promoting cell proliferation in various plant organs (Omidbakhshfard et al., 2015). In maize, the ectopic expression or mutation of specific *GRF*s has been shown to significantly affect the size of leaves and internodes (Wu et al., 2014; Nelissen et al., 2015). The expression of *GRF*s along the maize leaf gradient exhibits a spatially dynamic pattern, tightly regulating the transition between cell division and cell expansion (Nelissen et al., 2015). In addition, hormone distribution and signaling play a crucial role in regulating organ growth. In maize leaves, the basal region primarily depends on auxin and cytokinin to drive cell division, whereas gibberellin synthesis determines the size of the active cell division zone (Nelissen et al., 2012; De Vos et al., 2020). The coordinated action of these hormones ensures proper growth dynamics by regulating the balance between proliferation and expansion. Beyond these internal factors, environmental conditions such as drought can seriously affect maize organ growth. Drought restricts maize leaf growth by inhibiting cell division in the meristem and cell expansion in the elongation zone, thus resulting in a reduced growth rate (Avramova et al., 2015). Although maize plants attempt to compensate for the reduced growth rate by increasing leaf growth duration, this remains insufficient to achieve the same final leaf size as in non-droughted plants, ultimately resulting in yield losses under drought stress (Nelissen et al., 2018). Notably, the GRF transcription factors are also involved in maintaining the growth potential under mild drought (Van Hautegem et al., 2025), highlighting the importance of transcriptional networks integrating developmental and environmental cues to optimize plant growth and productivity.

Spatiotemporal analysis of gene expression plays a crucial role in understanding developmental programs and responses to environmental stresses. Over the years, several transcriptome profiling studies have been conducted to investigate the dynamics of gene expression and cellular processes in various maize organs. The highly regular and continuous manner of maize leaf growth is characterized by coordinated and localized transitions in gene expression. These transitions produce major biochemical shifts along the proximal-distal leaf gradient, from a prevalence of primary cell wall and basic cellular metabolism at the leaf base to secondary cell wall biosynthesis and C4 photosynthetic development toward the tip (Li et al., 2010). The transcriptional changes within the division and elongation zones discriminate each zone into a basal and distal part, fine-tuning the linear growth of the leaf (Nelissen et al., 2018). Similar to leaf, transcriptomic investigations on internode growth also revealed significant variations in global gene transcription patterns along the elongating internode, differing between the meristem, elongation zone, and mature zone of the internode (Zhang et al., 2014). Genes associated with cell wall synthesis exhibit notable spatial differences along the internode zones, with genes related to lignin and secondary metabolite synthesis expressing more prominently in the upper zones compared to the lower zones, thereby conferring greater mechanical strength to the upper part of the internode (Zhang et al., 2014; Wang et al., 2019b). However, these studies were conducted on a single leaf or internode and at a single time point, and hence lack comprehensive temporal resolution of the gene expression dynamics during organ development.

In this study, we combined both phenotypic and transcriptomic approaches to investigate the spatiotemporal developmental characteristics and gene expression of maize shoot organs during normal growth and under drought. We employed a new image-based method to dynamically analyze internode elongation and used epidermal imprints to investigate the internode growth zones at the cellular level, comparing the growth of internodes and leaves at the macroscopic and microscopic level. We found a much shorter elongation duration in internodes compared to leaves. Leaves have a long steady-state growth phase in which both the division zone and elongation zone are at their full size, whereas the maximum sizes of different internode zones were more temporally separated, with the division zone reaching its maximum size early on in growth and then reducing as the elongation zone expands. Through a comprehensive spatiotemporal transcriptome analysis of the development of leaves and internodes of different ranks, similar gene expression trends along the leaf growth gradient and the internode development stages were observed, suggesting that the core growth regulatory network is conserved in both organs. The high-density transcriptomes of leaves, internodes, and ear during their development under both well-watered and drought conditions reveal the global and spatiotemporally complex effect of drought on maize development and resulted in the identification of 1,444 organ-specific genes, providing a valuable resource for subsequent research of shoot organ-related traits in maize.

## Results

### Internode elongation follows the same sigmoidal pattern as leaf elongation

In maize plants, the stem and its constituent internodes are hidden from view by the encircling leaf sheaths, complicating the study of internode growth. However, internode length and growth can be estimated from the heights of the collars of the leaves attached at the nodes above and below the internode of interest (Figure 1A) (Fournier and Andrieu, 2000). Based on this approach, we developed an image-based method to estimate the internode length during development, using an automated plant phenotyping platform equipped with gravimetric watering and an RGB imaging system that images plants from multiple angles each day (Verbraeken et al., 2021). The performance of this method was evaluated on a dataset of 163 ground-truth internode length measurements, for internodes 9 (Int9), 12 (Int12), and 15 (Int15, where the primary ear is typically located) subjected to well-watered (Control), prolonged drought (starting at V5 during development, V5-Drought), and reproductive drought (V12-Drought) treatments (see Materials and methods; Supplemental Data Set 1; Supplemental Figures 1-3; Supplemental Tables 1-2). Using Leave-One-Out-Cross-Validation (LOOCV), an R^2^ value of 0.88 and a root-mean-square error (RMSE) value of 7.37 mm were obtained (Supplemental Figure 4A). Notably, the method by Fournier and Andrieu (2000) requires that the leaf collar i+1 is visible, which limits its applicability in the early stages of internode growth. Therefore, we developed an indirect internode length estimation method, using a linear model based on the Δcollar value of the highest visible pair of leaf collars, without going below leaf collar i-3 (see Materials and methods). This indirect length estimation method was evaluated for a dataset of 207 ground truth measurements for internodes 9, 12 and 15 in which collar i+1 was not visible in the image (Supplemental Data Set 1; Supplemental Tables 1-2), giving a LOOCV R^2^ value of 0.82 and a RMSE value of 2.46 mm (Supplemental Figure 4B). Combined, both methods allowed us to follow internode growth from beginning to end, with a LOOCV R^2^ value of 0.98 and a RMSE of 5.23 mm (n=370, Figure 1B).

**Figure 1.**
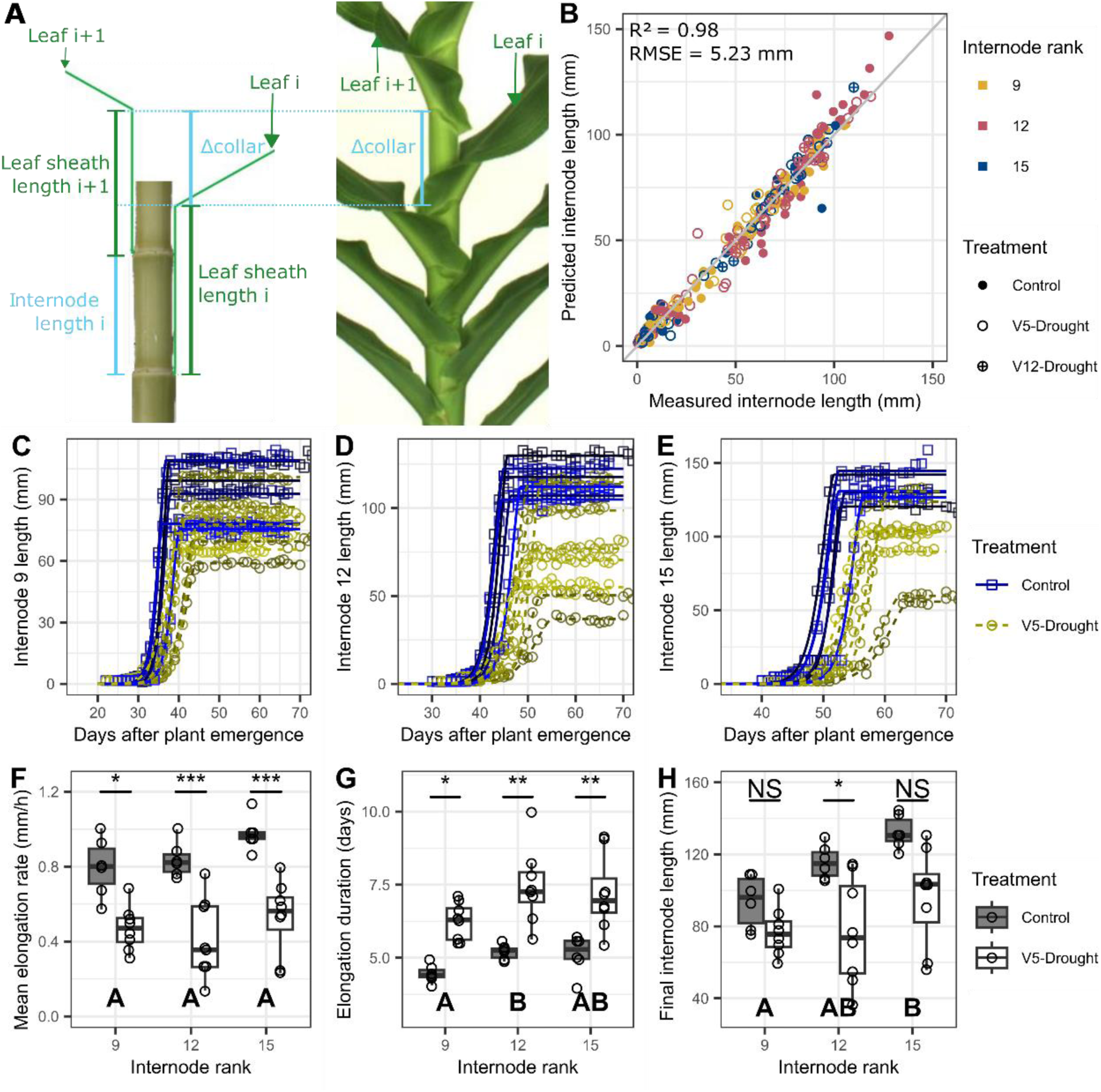
Image-based measurements of internode length and visualization of the internode growth traits. **A)** Illustration of the relation between the length of an internode i, the positions of the leaf collars of the leaves below and above the internode (i and i+1, respectively) and the lengths of the leaf sheaths of these leaves. Fournier and Andrieu (2000) estimated the length of internode i based on the difference in collar height between collars i and i+1 (Δcollar) and the difference in lengths of leaf sheaths i and i+1 (Δsheath) as: Internode length_i_ = Δcollar-Δsheath. **B)** Estimation of lengths along the entire range of internode elongation by combining both the direct and indirect internode length estimations. **C-E)** Internode length measurements for individual internodes 9 (C), 12 (D) or 15 (E) over time represented by squares (Control treatment) or circles (V5-Drought treatment) with a beta-sigmoid curve fitted to each individual internode represented as a solid line (Control treatment) or dashed line (V5-Drought treatment). Each plant is represented in a different color intensity, the same intensities refer to the same plant in C, D and E. **F-H)** Comparison of internode growth traits for different internode ranks and contrasting watering treatments. Letters A-B at the bottom indicate statistically significant differences (Holm-corrected p<0.05) between internode ranks: if two internode ranks share a letter, there is no significant difference between these two groups. Stars at the top indicate statistically significant differences between treatments within the internode ranks (***: Holm-corrected p<0.001, **: p<0.01, *: p<0.05, NS: p>0.05), F: Mean internode elongation rate (mm/h), G: Internode elongation duration (days), H: Final internode length (mm).

Using these internode length estimation methods, the lengths of internodes 9, 12 and 15 were followed over time in 14 plants, six of which were subjected to well-watered conditions and eight to the V5-Drought treatment. Beta-sigmoid growth curves were fitted individually for all 42 internodes, with R^2^ values ranging from 0.982 to 0.999 (Figure 1, C-E; Supplemental Figure 5; Supplemental Data Set 2), demonstrating that internode elongation follows a sigmoidal pattern similar to that of leaf elongation (Voorend et al., 2014). Final internode length, internode elongation duration and mean elongation rate over the growth period were then derived from the fitted growth function (Figure 1, F-H). Final internode length increased with the internode rank, with internode 15 significantly longer than internode 9 (Figure 1H; Supplemental Data Set 2). Elongation duration was highest in internode 12 when combining the treatments, and significantly higher than in internode 9 (Figure 1G; Supplemental Data Set 2). The mean elongation rate was highest in internode 15, though not significantly higher than in internodes 9 and 12 (Figure 1F; Supplemental Data Set 2). It is noteworthy that the trends over internode rank, in both well-watered and drought conditions, are different to trends previously observed over leaf rank (Verbraeken et al., 2021). In the internodes, an increasing trend was observed with rank for both elongation rate and duration, leading to an increase in final internode length with rank (Figure 1, F-H; Supplemental Table 3), while for the adult-stage leaves, leaf elongation rate (LER) displayed a decreasing trend with leaf rank while leaf elongation duration (LED) displayed an increasing trend, resulting in a net decreasing trend for final leaf length (FLL) (Verbraeken et al., 2021).

Under V5-Drought conditions, the mean elongation rates in internodes 9, 12 and 15 were reduced by 41.7%, 51.0% and 46.3%, respectively, while elongation duration increased by 40.9%, 44.1% and 41.2% (Figure 1, F-G; Supplemental Table 3). This resulted in a final internode length reduction of 19.5%, 34.4% and 28.2% in internodes 9, 12 and 15, respectively (Figure 1H; Supplemental Table 3). Similar to leaves, the final length of internodes is determined by the interplay between elongation rate and elongation duration. Both leaves and internodes had a similar response to drought, increasing the elongation duration to partially compensate for a decreased elongation rate under drought (Figure 1, F-H; Supplemental Table 3) (Nelissen et al., 2018; Verbraeken et al., 2021).

### Cellular analysis shows a spatiotemporal growth regulation in maize leaves

We investigated whether the similarities between internode and leaf growth at the macroscopic level also extend to the cellular organization of the internode and leaf growth zones. To observe the leaf growth zone over time, the basal 14 cm of growing leaves 9 (L9), 12 (L12), 15 (L15) and 18 (L18) were sampled on the day of leaf emergence (x0), 4 days after leaf emergence (x4) and 8 days after leaf emergence (x8), for both the Control and V5-Drought treatments, and the sizes of the leaf lamina division zone and elongation zone were determined (see Materials and methods; Supplemental Figures, 6-7; Supplemental Data Set 3) (Nelissen et al., 2013; Sprangers et al., 2016).

A 3-way ANOVA showed that division zone size varied significantly with leaf rank (p=2.94×10⁻⁷) and sampling time (p=1.62×10⁻⁶) but not with watering treatment (p=0.11), and there were no significant interactions (Figure 2, A-B; Supplemental Data Set 3). For both treatments combined, the division zone size was significantly larger in leaf 12 and leaf 15 compared with leaf 9 and leaf 18, and was smaller at time point x8 compared with time points x4 and x0 (Supplemental Table 4; Supplemental Data Set 3). Comparing the division zone size with the macroscopic growth traits, division zone size correlated positively with FLL (r=0.387, p=0.001434) and negatively with relative leaf growth progression (the ratio between immediate length and FLL, r=-0.451, p=1.634*10^-4^; Figure 2, E-F), but showed no correlation with estimated LER at time of sampling (r=0.107, p=0.3946) or total LED (r=-0.106, p=0.4026; Supplemental Figure 8). Overall, division zone size tended to be larger in leaf ranks with a higher FLL and decreased within each rank as leaves progressed through their growth curve.

**Figure 2.**
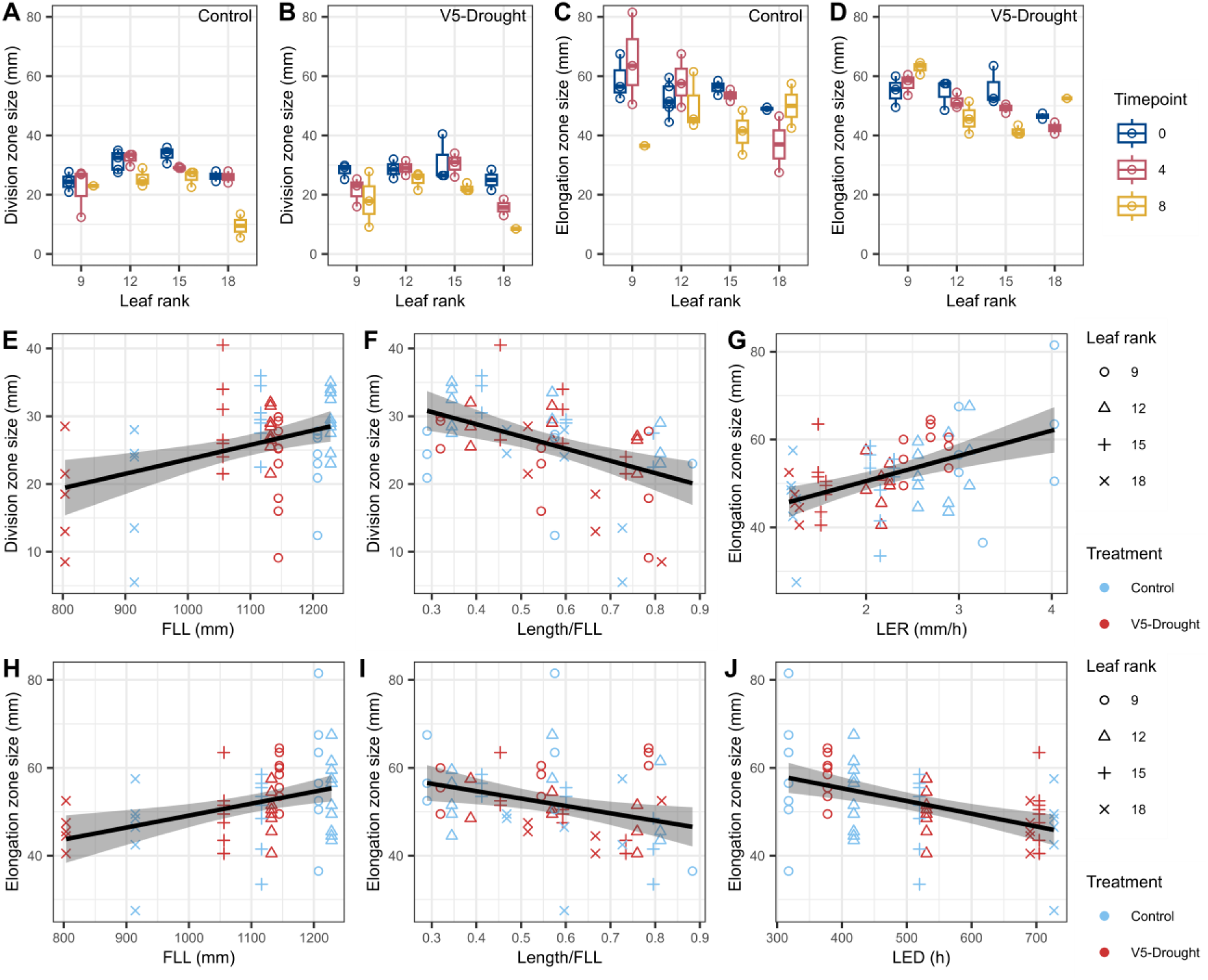
Cellular analysis of division zone and elongation zone size in relation to growth traits in maize leaves. **A-D)** Size of the division zone (A, B) and elongation zone (C, D) at the base of the leaf lamina, for the Control (A, C) and V5-Drought (B, D) treatments, for the different leaf ranks (x-axis: 9, 12, 15 and 18), at the three sampling time points (color; 0, 4 or 8 days after leaf emergence). **E-J)** Comparison between division zone or elongation zone size and macroscopic growth traits. E: Division zone size over FLL. F: Division zone size over relative leaf growth progression (length/FLL). G: Elongation zone size over instantaneous LER. H: Elongation zone size over FLL. I: Elongation zone size over relative leaf growth progression (length/FLL). J: Elongation zone size over LED. Each growth zone sample for the two treatments (color; Control and V5-Drought), four leaf ranks (shapes: 9, 12, 15 and 18), and the three sampling time points (day of leaf emergence, 4 days after, 8 days after) was matched to the FLL, leaf length, LER and LED derived for that leaf rank, treatment and sampling time point based on the fitted growth curves in Verbraeken et al. (2021). Black lines are linear regressions fitted to the data, shaded areas indicate the 95% confidence interval for the fitted lines. FLL, full leaf length; LER, leaf elongation rate; LED, leaf elongation duration.

For the lamina elongation zone size, a significant interaction effect was found between sampling time point and leaf rank (p=0.007851), and between sampling time point and treatment (p=0.002533), but not between leaf rank and treatment (p=0.8612; Figure 2, C-D; Supplemental Data Set 3). This indicates that the effect of time on elongation zone size differs between the leaf ranks and between the treatments. Considering the effect of time point over all leaf ranks and treatments combined, a significant decrease in elongation zone size occurred between the day of leaf emergence and 8 days after leaf emergence (p=0.01998; Supplemental Data Set 3). Elongation zone size correlated strongly with immediate LER (r=0.479, p=5.523*10^-5^, Figure 2G), so within an individual leaf elongation zone size increased until maximal LER was reached, followed by a decrease. Over the leaf ranks, this resulted in a positive correlation between elongation zone size and FLL (r=0.376, p=0.002032; Figure 2H) and a negative correlation with relative leaf growth progression (r=-0.320, p=0.009462; Figure 2I) as well as LED (r=-0. 465, p=9.512*10^-5^; Figure 2J).

Taken together, the division zone appears to set up the potential FLL early on in growth and diminishes in size as leaf growth progresses, while the elongation zone size increases and decreases along with the LER as the leaf progresses through its growth curve.

### The internode growth zone closely resembles the leaf growth zone but with distinct timing

To compare the internode growth zone with that of the leaf, we analyzed internode 12 growth at the cellular level using epidermal imprints at the V12, V13, V14, and V15 stages, covering its entire growth period (Supplemental Figure 9; Supplemental Data Set 4). Along the internode, we observed a zone of small, consistently sized cells at the base, resembling the leaf division zone, followed by a zone of gradually increasing cell lengths, akin to the leaf elongation zone. The sizes of the growth zones for internode 12 were then determined (see Materials and methods; Supplemental Figure 10). At V12, the internode consisted mostly of the division zone, with only a small elongation zone (Figure 3A; Supplemental Table 5). By V13, the elongation zone had increased in size and the first mature cells appeared, with a final cell length of approximately 75 µm (Figure 3A; Supplemental Table 5). By V14, the division zone had shrunk while the elongation zone had further increased in size (Figure 3A; Supplemental Table 5). By V15, both zones had decreased in size (Figure 3A; Supplemental Table 5). As internode elongation had ceased at this stage (Supplemental Figure 9), the not yet fully expanded cells at the base of the internodes indicated a possible sudden end to the growth process. The early elongating cells reached a mature cell length of approximately 75 µm, whereas those elongating after the V13-stage expanded to a larger cell length, around 150 µm (Figure 3A).

**Figure 3.**
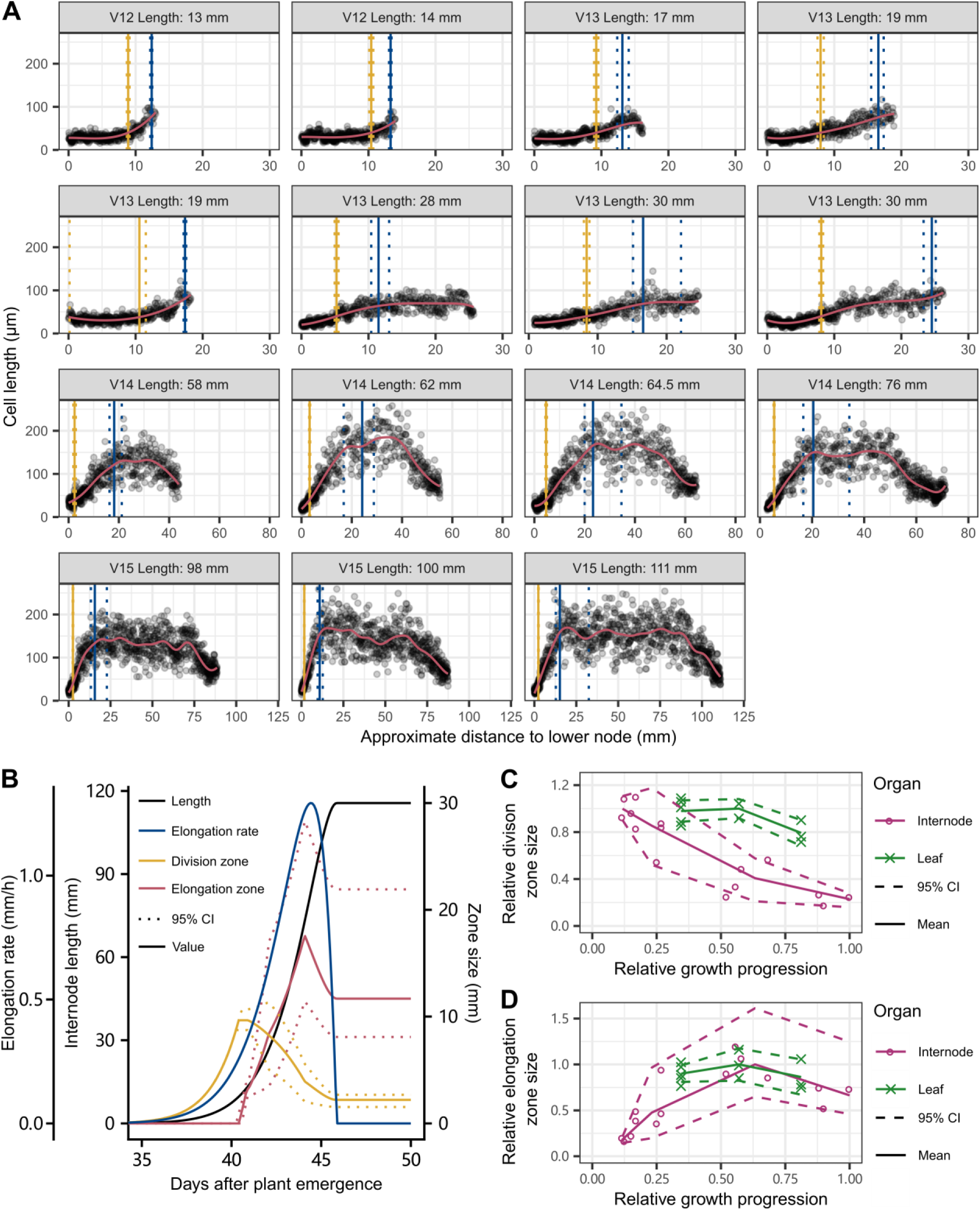
Cell length distribution and growth dynamics of internode 12 under control conditions. **A)** Cell length along the internode 12 sampled at different V-stages and different lengths, organized from smallest and youngest to largest and oldest. Graph titles indicate V-stage and internode length. X-axis sizes vary according to V-stage. A local polynomial smoothing fit to the cell length measurements, used to determine the top of the division zone and elongation zone, is indicated in red. The yellow vertical line indicates the mean end of the division zone, with the dotted yellow lines indicating the 95% confidence interval. The blue vertical line indicates the mean end of the elongation zone, with the dotted blue lines indicating the 95% confidence interval. **B)** Internode 12 length (solid black line, primary left y-axis) and elongation rate (solid blue line, secondary left y-axis) over time for well-watered plants, combined with the mean estimated size of the division zone and elongation zone (solid yellow and red lines, right y-axis) over time and their lower and upper 95% confidence interval (CI). In the early stages of internode growth, the entire internode is division zone and the CIs for division zone and elongation zone have zero width. **C-D)** In order to normalize for differences in elongation duration, time was expressed as relative growth progression (length/final length) and division zone (C) and elongation zone (D) size were normalized relative to their maximum observed value within either internode 12 or leaf 12. Observations for leaf 12 ranged from 34% to 81% of final length.

By integrating the mean internode 12 growth curve with the relationship between zone size and internode length (see Materials and methods; Supplemental Figure 11), we estimated the temporal changes in the size of the division zone and elongation zone in internode 12 (Figure 3B). The internode 12 elongation zone size exhibited similar patterns over time as the elongation zone size of individual leaves, initially increasing and later decreasing in sync with the increase and decrease in elongation rate (Figure 3B; Figure 2G). The internode 12 division zone size peaked early on in internode elongation, followed by a gradual, steady decline after formation of the elongation zone (Figure 3B). To further compare the trends in division zone and elongation zone size over time between internode and leaf, we examined their relative sizes along the growth progression (Figure 3, C-D). During the observation period for leaf 12, from leaf emergence to 8 days after leaf emergence (34% to 81% of FLL), the elongation zone size of internode 12 and leaf 12 displayed a similar trend, increasing first and then decreasing after peaking at around 60% of final length (Figure 3D). The division zone size in leaf 12 initially remained constant, followed by a slight decline after the leaf reaching around 60% of final length (Figure 3C), resulting in both a large division zone and elongation zone being present at the peak of leaf elongation. In contrast, the division zone size in internode 12 consistently decreased over time in the observed developmental time frame (Figure 3C), resulting in a substantial reduction of division zone size by the time the elongation zone size and internode 12 elongation peak. These observations indicate a more pronounced separation in time between cell division and elongation in elongating internodes than in leaves, where cell division and elongation co-occur for a longer period.

### Generation of a high-resolution transcriptomic atlas of growing maize shoot organs

The growth of both maize leaves and internodes is directed from a linearly organized growth zone at the base of the organ, consisting of a spatially separate division and elongation zone. To understand if these structural similarities between leaves and internodes are reflected in similar genetic regulation of growth processes in both organs, and to assess the spatiotemporal gene expression landscapes during the development of maize shoot organs, we generated a transcriptomic atlas of growing maize leaves and internodes, as well as of the maize ear, whose morphogenesis is one of the key target traits for maize breeding. The leaf division zone, elongation zone, differentiation zone (where cells stop elongating but are not fully photosynthetically active), and mature zone (where cells are fully developed) were sampled from leaves L9, L12, L15, and L18 at leaf emergence, 4 days after leaf emergence and 8 days after leaf emergence. Additionally, whole leaves (L9, L12, L15 and L18) were sampled before their emergence and the mature part of L12, L18 and the ear leaf (LeafEar, usually L14 or L15) was sampled at several timepoints during the reproductive stage (Figure 4A). Internodes Int9, Int12 and Int15 were sampled throughout their growth during the vegetative stage and the ear internode (IntEar) was sampled at several time points during the reproductive stage. Different internode zones were sampled by dividing the internode lengthwise into 1 zone, 2 zones (low zone and high zone) or 3 zones (low zone, mid zone, and high zone), depending on the sampling time points relative to the timing of leaf emergence, ear emergence or silking (Figure 4A). The maize ear was also sampled throughout its growth, from 4 days before ear emergence (Ear_4) until silk emergence (Silk), and divided lengthwise into sampling zones similarly to the internode (Figure 4A). To investigate the molecular mechanisms underlying drought responses throughout the development of these organs, all these tissues were sampled for the Control treatment (until silking), the V5-Drought treatment (from V5-stage to silking) and the V12-Drought treatment (from V12-stage to silking). In total, 272 distinct biological materials (varying in terms of organ, organ rank, organ zone, sampling time point or watering treatment) were sampled in each of three independent greenhouse experiments (Supplemental Table 6).

**Figure 4.**
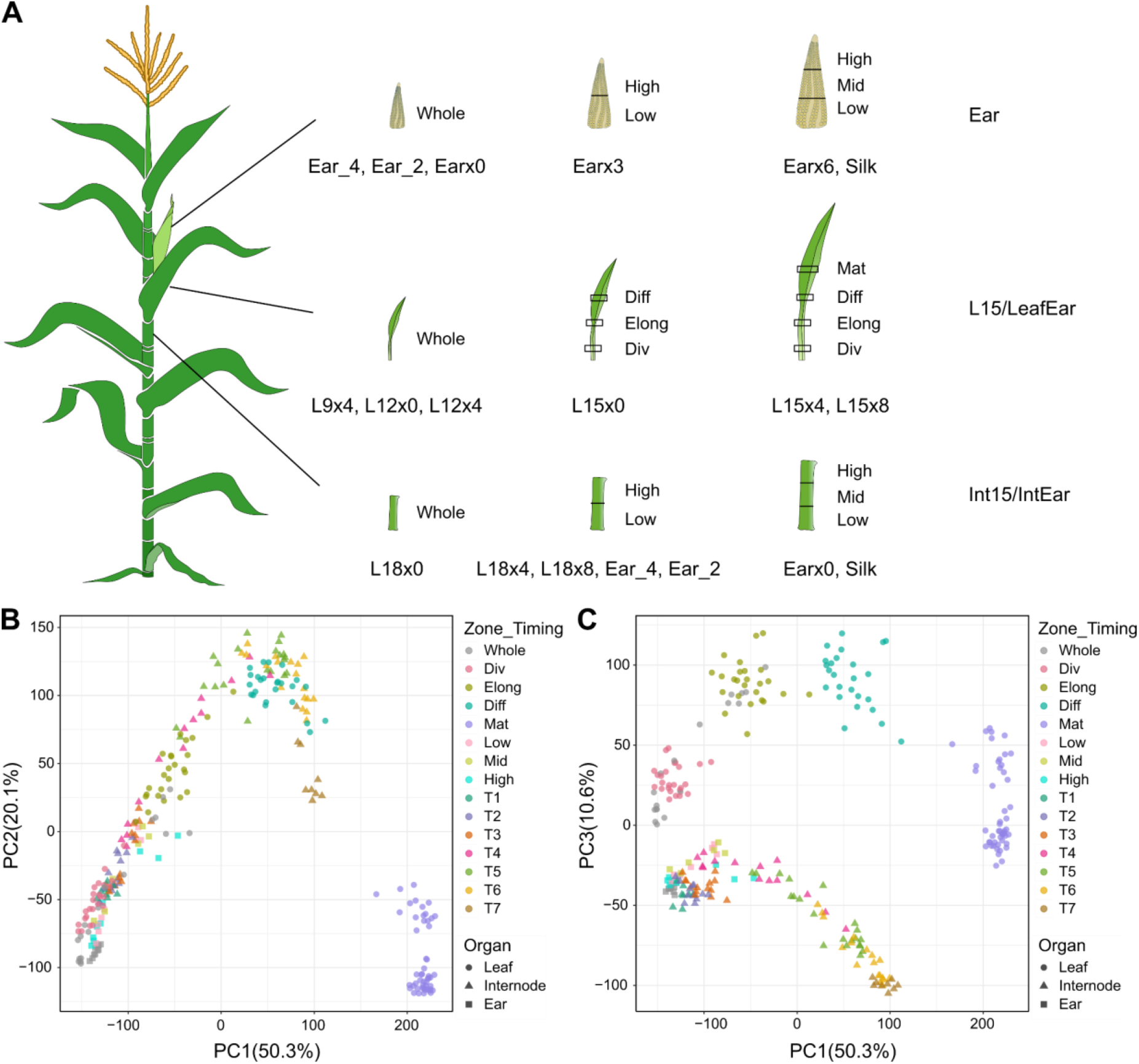
Sampling strategy and principal component analysis of transcriptomic data. **A)** Graphic representation of transcriptomic sampling for leaves, internodes, and the ear. Leaves (L9, L12, L15/LeafEar, L18, with only L15/LeafEar shown) were sampled before emergence (whole leaf), at emergence (x0), and 4 days (x4) and 8 days (x8) after emergence. Emerged leaves were sampled with different zones: division zone (Div), elongation zone (Elong), differentiation zone (Diff), and mature zone (Mat). Internodes (Int9, Int12, Int15/IntEar, with only Int15/IntEar shown) were sampled based on timing relative to leaf emergence, ear emergence, or silking and divided lengthwise into two or three zones: low zone (Low), middle zone (Mid), and high zone (High). The ear was sampled from 4 days before emergence (Ear_4) to silk emergence (Silk) and divided lengthwise into two or three zones. **B)** The percentage of variation among tissues explained by each principal component is displayed on both the PC1 and PC2 axes. **C)** The percentage of variation among tissues explained by each principal component is displayed on both the PC1 and PC3 axes. Point shape indicates the organ type. For leaf and ear samples, the colors indicate different zones, while for internode samples, the colors indicate different temporal stages. T1-T7, temporal stages from Timing 1 to Timing 7.

For each biological replicate, an average of 31 million 125 bp paired-end RNA sequencing reads were generated. Across samples, 51.4 to 75.8% (67.9% on average) of the raw reads could be uniquely mapped to the B73 reference genome (Supplemental Data Set 5). Summarizing read counts across replicates resulted in a matrix of 39,323 genes. Only genes with more than 0.5 counts per million (cpm) in at least three samples were considered as expressed, resulting in a pruning of the gene set to 28,342 genes (72.1%, Supplemental Data Set 6). We then examined the number of expressed genes in individual organ zones throughout development. The sampling time points for each tissue were grouped into temporal stage classes (T1 to T9) to make them comparable between different organs and organ ranks (Supplemental Table 6). Notably, the number of expressed genes was observed to increase over time in all three organs (Supplemental Figure 12A). Among the leaf zones, the differentiation zone expressed the largest number of genes throughout development (Supplemental Figure 12B). In internodes, the high zone featured more expressed genes than the low zone from T2 to T4, while in later stages, high and low zones expressed similar numbers of genes and higher numbers than the middle zone (Supplemental Figure 12B). In the ear however, similar numbers of genes were expressed across the different zones (Supplemental Figure 12B).

A principal component analysis (PCA) was performed for all samples using the mean expression across biological replicates. The first two principal components (PCs) explained 50.3% and 20.1% of the variation, respectively (Figure 4B). Both PCs contributed to the separation of leaf samples by developmental zone, which reflects a strong heterogeneity of gene expression along the longitudinal axis of developing maize leaves. In contrast, the expression profiles of internode samples were separated primarily according to different temporal stages in PC1 and PC2, with the internode zones in the early temporal stages grouping closer to the leaf division zone, and moving along the leaf elongation zone samples to the leaf differentiation zone for later temporal stages (Figure 4B). The third PC, which accounts for 10.6% of the total variance, distinguished the leaf samples from the internode and ear samples, with all the ear samples clustered together with the early-stage internode samples (Figure 4C), indicating transcriptome similarity during ear development and early internode development.

### Spatiotemporal gene expression during maize leaf, internode and ear development

To reveal the transcriptional features associated with developmental events during the growth of each organ, we identified genes showing differential expression (|log_2_FC|>1 and Benjamini-Hochberg FDR-adjusted *P*-value <0.05) among zones and time points in maize leaf, internode, and ear. L15 was used to identify the spatiotemporal differentially expressed genes (DEGs) during leaf development (Supplemental Data Set 7). Differential gene expression analysis among leaf zones revealed that a large number of genes varied in expression between any two leaf zones (Supplemental Table 7). Along the leaf proximodistal axis, genes related to photosynthesis were significantly upregulated while genes involved in cell cycle processes were downregulated according to GO enrichment analysis (Figure 5A). Relative to the division zone, genes with functions in water transport, peptide transport, and nitrite assimilation were upregulated in the elongation zone (Figure 5A), indicating increased transport in the leaf elongation zone. In the leaf differentiation zone, genes associated with lignin biosynthesis and cell wall thickening were significantly upregulated compared to the elongation zone (Figure 5A), coinciding with the transition from cell expansion to cell strengthening and maturation. Towards the mature zone, upregulated genes were primarily enriched for genes involved in photosynthesis, whereas genes related to cell growth, phloem or xylem histogenesis and wall biogenesis were downregulated relative to the differentiation zone (Figure 5A). To gain further insight into the transcriptomic changes in each leaf zone over time, DEGs were identified across three time points (0, 4, 8 days after leaf appearance) in each leaf zone (Supplemental Data Set 7). Compared with the DEGs among zones, fewer genes were observed to be differentially expressed between time points (Supplemental Table 8). GO enrichment was performed on DEGs between 8 days and 0 days after leaf appearance in division, elongation and differentiation zones. In the leaf division zone, genes involved in regulation of transcription, stomatal complex development, shoot system development and responses to gibberellin and auxin were significantly downregulated over time (Figure 5A). Genes downregulated over time in the leaf elongation zone were primarily associated with cell cycle processes, whereas upregulated genes were enriched for genes involved in cell wall organization, nitrate transport and photosynthesis (Figure 5A). In the leaf differentiation zone, genes involved in suberin and phenylpropanoid biosynthetic processes were upregulated over time (Figure 5A), reflecting enhanced secondary cell wall formation. For the leaf mature zone, differential expression analysis was conducted between 8 days and 4 days after leaf appearance. Notably, several genes related to photosynthesis including light and dark reaction were observed to be downregulated (Figure 5A), which might be due to a decreased sink demand from the shrinking division zone and elongation zone in L15 at 8 days after leaf appearance (Figure 2, A and C).

**Figure 5.**
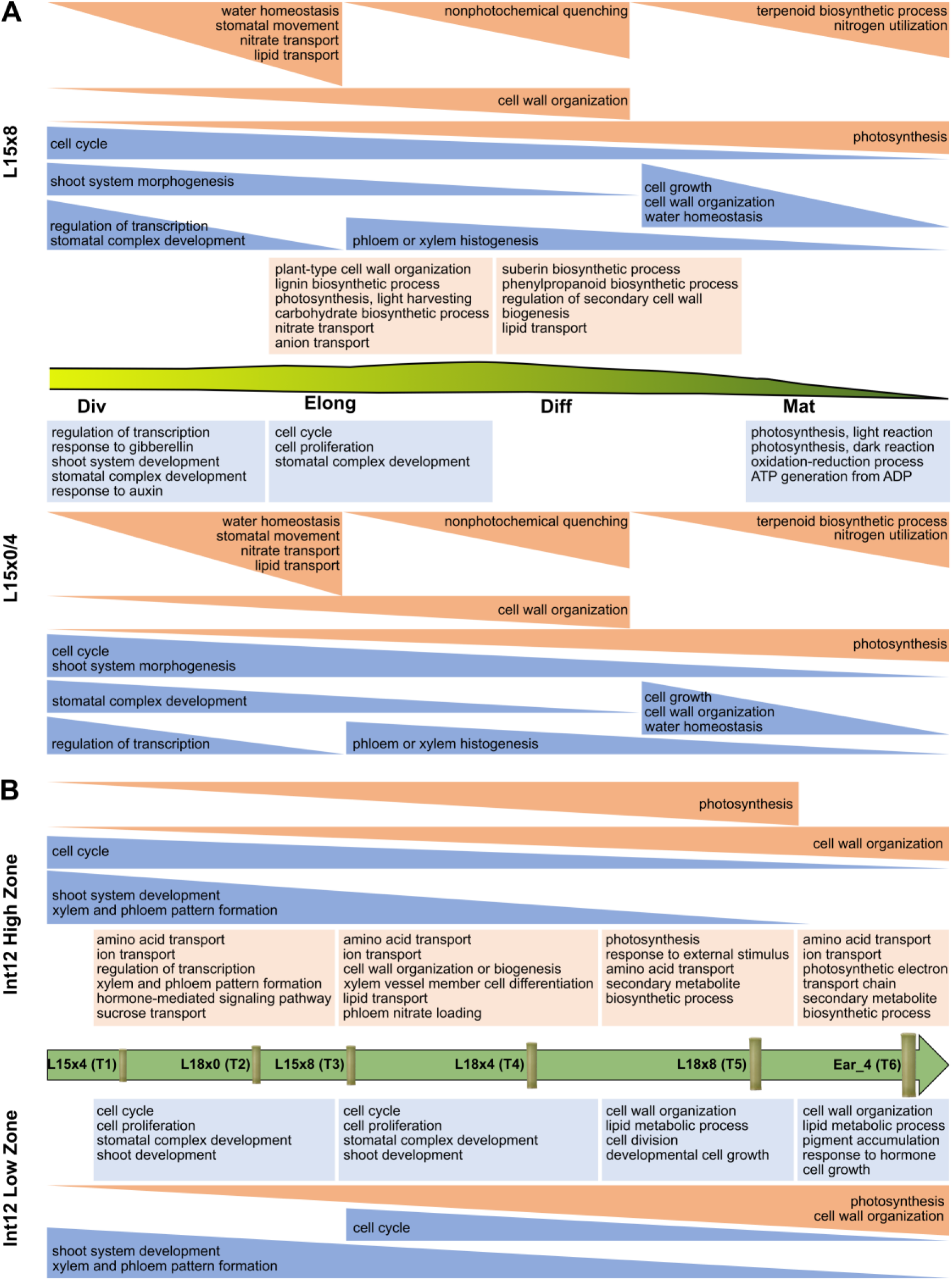
Schematic overview of the main enriched GO categories among DEGs during maize leaf and internode development. **A)** Overview of the main categories of spatiotemporal DEGs during leaf 15 (L15) development. The GO categories of genes upregulated along the leaf zones at individual time points are indicated in orange, while those downregulated are indicated in blue. The GO categories of genes upregulated over time in individual zones are indicated in light orange, while those downregulated are indicated in light blue. Div, division zone; Elong, elongation zone; Diff, differentiation zone; Mat, Mature zone. **B)** Overview of the main categories of spatiotemporal DEGs during internode 12 (Int12) development. The GO categories of genes upregulated along internode temporal stages in individual zones are indicated in orange, while those downregulated are indicated in blue. The GO categories of genes upregulated in the high zone compared with low zone are indicated in light orange, while those downregulated are indicated in light blue. T1-T6, temporal stages from Timing 1 to Timing 6.

In contrast to the leaf samples, the global transcription patterns of internode samples indicated a more pronounced effect of development stages on gene expression compared to the spatial zoning of the internode (Figure 4B). To better understand the biological processes in each internode sampling zone during development, we matched each sampling zone (low, mid and high) to the development zones (division, elongation and mature) in internode 12 based on the relative internode length. During the early stages from T1 to T3, the entire internode was made up of division zone, while at T4, the low zone included both division zone and elongation zone, and the high zone consisted of elongation zone and mature zone (Supplemental Figure 13). In the later stages, T5 and T6, the middle and high zones were entirely composed of mature zone, while the low zone included division zone, elongation zone and mature zone (Supplemental Figure 13). The number of DEGs, relative to the earliest temporal stage (T1), increased progressively during development in each internode sampling zone (Supplemental Data Set 8, Supplemental Table 9). In both the low zone and high zone of the internode, the expression of genes involved in photosynthesis and cell wall organization were upregulated over time, while genes associated with shoot system development as well as xylem and phloem pattern formation were downregulated (Figure 5B). It is noteworthy that although the entire internode was division zone during the early stages (T1 to T3), the expression of cell cycle genes in the high zone was downregulated from the earliest stage to the late stages, while in the low zone, the downregulation of cell cycle genes was only seen starting from T3 (Figure 5B), indicating a higher cell division activity in the internode low zone compared with the high zone during the early stages. The number of DEGs between the internode high zone and low zone increased from T1 to T4 (Supplemental Table 10). Genes that were more highly expressed in the high zone were enriched for genes involved in amino acid transport, ion transport, xylem and phloem pattern formation, hormone-mediated signaling and cell wall organization (Figure 5B), implying more active transport and cell differentiation took place in the internode high zone. Towards the later stages (T5 and T6), genes related to photosynthesis and secondary metabolism were upregulated in both the high zone and middle zone relative to the low zone, while genes involved in cell wall organization were upregulated in the low zone (Figure 5B), consistent with the cellular observations showing that the high zone and middle zone were fully matured during the late stages (Supplemental Figure 13).

Besides leaf and internode, we also identified spatiotemporal DEGs during ear development (Supplemental Data Set 9). In the development of a young ear, the inflorescence meristem branches several times to produce lateral meristems with determinate fates (Chuck et al., 1998). The primary and secondary branching that form the spikelet pair meristem and two spikelet meristems affect the number of spikelets per row and the number of rows per ear, both of which are fully determined during the early stages of ear elongation (Verbraeken et al., 2021). In accordance with this, genes that are known to regulate the branching architecture of the maize inflorescence, *RAMOSA1* (*RA1*) and *TASSELSEED4* (*TS4*) (Vollbrecht et al., 2005; Chuck et al., 2007), were highly expressed at the earliest time point (4 days before ear emergence, Ear_4) and decreased gradually from early to late stages (Supplemental Figure 14). Additionally, the expression of a subtilisin encoding gene *ZmSBT31* homologous to Arabidopsis *SENESCENCE-ASSOCIATED SUBTILISIN PROTEASE* (*SASP*), which is involved in determination of silique number (Martinez et al., 2015), was also found to decline significantly during ear development (Supplemental Figure 14). After the formation of the spikelet meristems, each of the spikelet meristems produces two floral meristems. Several genes from the MADS-box family which is known to regulate floral organ identity (Bortiri and Hake, 2007), such as *ZMM17* and *ZmMADS41*, were found to significantly increase in expression over time, showing highest expression at the silking stage (Supplemental Figure 14). The maize homolog of Arabidopsis *FLOR1*, which encodes a leucine-rich repeat protein that interacts with MADS-box transcription factors and is involved in promoting flowering (Torti et al., 2012), was also among the upregulated genes during the rapid ear elongation stages (Earx3 and Earx6) (Supplemental Figure 14). In the silking stage, a large number of genes were differentially expressed relative to the preceding rapid ear elongation stages (Supplemental Table 11). Genes related to the cell cycle, flower development, cell differentiation, xylem and phloem pattern formation, auxin transport and gibberellin biosynthesis showed significant downregulation in both the low zone and high zone of the ear, indicative of reduced meristem activity in the silking ear. Conversely, genes that are involved in responses to stimuli, ion transport, secondary metabolism and cell wall organization showed increased expression at silking. Many DEGs were observed between the high zone and low zone of the ear while very few DEGs were found between the middle zone and the low zone (Supplemental Table 12), implying a similar developmental progression in the low and middle zones of the ear. The downregulated genes in the ear high zone, relative to the low zone, were enriched for genes involved in secondary metabolism and wax biosynthesis, while the upregulated genes were enriched for genes involved in transcription, inflorescence development, trehalose biosynthesis and hormone-mediated signaling pathways, reflecting a higher inflorescence differentiation activity in the ear high zone than in the lower zones at the silking stage.

### Integration of gene activity and cellular function across different organs

To further compare gene expression patterns across different organs during development, a *k*-means clustering analysis was performed on all the control samples, grouping genes that were differentially expressed in at least three comparisons among zones or time points (24,880 genes) into 18 clusters (Figure 6A; Supplemental Data Set 10). In line with the PCA showing that the transcriptomic profiles of internode samples in the early temporal stages clustered closer to the leaf division zone (Figure 4B), genes in clusters 8, 9 and 12 which showed high expression in the leaf division zone also exhibited relatively high expression in internode zones at early temporal stages (Figure 6A). Genes involved in cell cycle and nucleic acid metabolic processes were significantly enriched in all these clusters (Supplemental Figure 15; Supplemental Data Set 10), which is consistent with fast cell division in leaf division zones and young internodes. Additionally, all ear tissues except for the samples at silking demonstrated high expression of genes in clusters 8, 9, and 12 (Figure 6A), indicating robust cell division activity throughout ear development prior to silking. Cluster 1 contained genes that were most highly expressed in the leaf elongation zone, which also showed elevated expression in both low and high internode zones at intermediate temporal stages when cell elongation was active in the whole internode (Figure 6A). The GO terms that were significantly enriched in cluster 1 include hydrogen transport, intracellular transport, protein transport, and ATP synthesis coupled proton transport (Supplemental Figure 15; Supplemental Data Set 10), which is in line with the acid growth hypothesis, according to which cell walls are loosened in part due to proton export, allowing cell turgor pressure to expand the cell (Cleland, 1971; Hager et al., 1971). Clusters 7 and 13 contained genes that were highly expressed in the leaf differentiation zone as well as in all late-stage internode zones (Figure 6A). These clusters were enriched for genes involved in cell wall organization or biogenesis (Supplemental Figure 15; Supplemental Data Set 10), in accordance with cell wall deposition in the corresponding tissues. Several genes that showed high expression in the late-stage internode zones were also observed highly expressed in leaf mature zone, as represented in clusters 4, 5, 6 and 14 (Figure 6A), and were enriched for genes involved in photosynthesis and response to external stimulus (clusters 4 and 5), carbohydrate and amino acid transport (cluster 5), cell wall organization (cluster 6), and ion homeostasis (cluster 14) (Supplemental Figure 15, Supplemental Data Set 10). In addition to clusters in which genes exhibited relatively high expression in a specific leaf zone or internode stage, several clusters, such as clusters 2, 3, 11, and 17, contained genes expressed across multiple leaf zones and internode stages (Figure 6A), suggesting common functional processes during transition between different leaf zones and internode stages. Furthermore, genes in cluster 10 and 18 exhibited higher expression in both leaf and ear compared with the internode, whereas the genes in cluster 15 were more expressed in internodes and ears than in leaves (Figure 6A), which may reflect specific functions in the corresponding organs.

**Figure 6.**
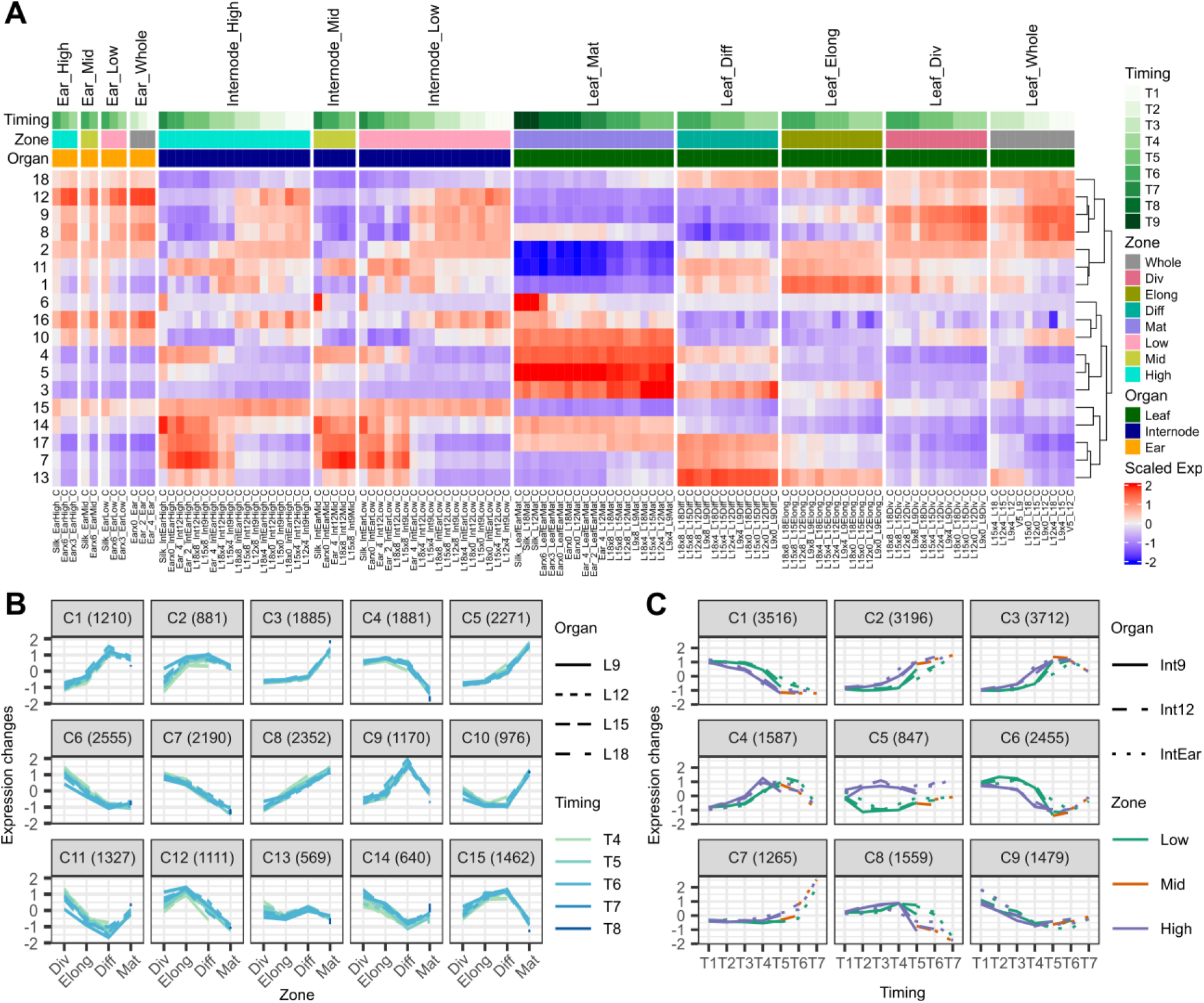
Dynamics of gene expression during maize development. **A)** *k*-means clustering grouped the gene expression profiles of all the control samples into 18 clusters. Gene expression means in each cluster are shown for individual conditions. Expression levels are color coded with red indicating high expression and blue indicating low expression. Whole, whole tissue; Div, division zone; Elong, elongation zone; Diff, differentiation zone; Mat, mature zone; Low, low zone; Mid, middle zone; High, high zone; T1-T9, temporal stages from Timing 1 to Timing 9. Sample names are formatted as Time point_Tissue zone_Treatment. **B)** *k*-means clustering grouped the leaf gene expression profiles into 15 clusters. Line types indicate the leaf rank and colors indicate the temporal stages. **C)** *k*-means clustering grouped the internode gene expression profiles into 9 clusters. Line types indicate the internode rank and colors indicate the zone.

The similarities in gene expression trends along the leaf zones and along the internode temporal stages suggest that a conserved growth regulatory network is shared between the leaves and internodes. To further compare gene expression profiles in the growth zones of these two organs, we classified genes in the leaf samples into 15 co-expression clusters and genes in the internode samples into 9 co-expression clusters (Figure 6, B-C; Supplemental Data Sets 11 and 12). The gene expression trends observed for different ranks of leaves or internodes were strikingly similar, indicating consistent developmental processes in different ranks of leaves or internodes. In leaves, the 15 co-expression clusters clearly corresponded to specific leaf zones. For instance, genes with the highest expression in the leaf division zone were found in clusters 6, 7, 11, and 14, while peak expression in the leaf elongation zone was found in cluster 4 and 12 (Figure 6B). The 9 co-expression clusters of internodes can be roughly divided into clusters peaking in expression in early (clusters 1, 6, and 9), middle (clusters 4 and 8), and late (clusters 2, 3, and 7) temporal stages, with one stage-neutral cluster (cluster 5) where genes were expressed at higher levels in the internode high zone than in the low zone across all stages (Figure 6C). We subsequently compared the genes between these internode clusters and leaf clusters. A statistically significant overlap of genes was identified between internode early-stage cluster 1 and leaf division zone clusters 6 and 7 (Supplemental Figure 16A; Supplemental Data Set 13), consistent with the observation from the overall transcriptome clustering. The overlapping genes were enriched for genes involved in cell cycle processes, including DNA replication and repair, histone modification, chromosome organization and separation, spindle checkpoint, and cytokinesis. Transcription factors that regulate cell proliferation were also found, including 11 *GRF* genes along with *ZmGRF-INTERACTING FACTOR1* (*ZmGIF1*), three homologs of Arabidopsis *AINTEGUMENTA* (*ANT*), and a PLATZ transcription factor *ZmPLATZ8* homologous to Arabidopsis *ORESARA15* (*ORE15*) which is involved in promoting the rate and duration of cell proliferation (Jun et al., 2019), indicating a conserved regulatory mechanism of cell division in both leaves and internodes. Additionally, genes related to epidermis development, such as *ZmSPCH2*, *ZmMUTE*, *ZmICE1c* and several *EPIDERMAL PATTERNING FACTOR*(*EPF*) genes (*ZmEPF1*, *ZmEPF10*, *ZmEPF12*, and *ZmEPFL2*), were also in the overlap. Strong overlaps between internode clusters and leaf clusters were also observed between internode middle-stage cluster 8 and leaf elongation zone clusters 4 and 12 (Supplemental Figure 16B; Supplemental Data Set 13). The set of overlapping genes was enriched for genes involved in unidimensional cell growth, containing homologs of Arabidopsis *LONGIFOLIA3* (*LNG3*) and *LNG4* which were reported to regulate longitudinal cell elongation (Lee et al., 2018), as well as homologs of Arabidopsis *SPIRAL1* (*SPR1*) and *WAVE-DAMPENED 2-LIKE3* (*WDL3*) which encode microtubule-associated proteins required for directional control of rapidly expanding cells (Yuen et al., 2003; Nakajima et al., 2004; Liu et al., 2013). Members of the BR signaling pathway, such as *ZmBR-SIGNALING KINASE 2* (*ZmBSK2*), *ZmBRASSINOSTEROID INSENSITIVE 1a* (*ZmBRI1a*), and homologs of Arabidopsis *TETRATRICOPEPTIDE THIOREDOXIN-LIKE1* (*TTL1*) genes that scaffold brassinosteroid signaling components at the plasma membrane (Amorim-Silva et al., 2019), were also found to be overlapping, which reflects the role of BR in regulating cell elongation in both leaves and internodes. Taken together, our analysis reveals a conserved growth regulatory network steering the growth processes of cell division and cell elongation in both leaves and internodes.

### Leveraging gene expression data to understand spatiotemporal regulation of organ development processes

To enable easy access to the transcriptomic atlas and visualize gene expression patterns, we developed a user-friendly shiny app in R which is available at https://www.psb.ugent.be/shiny/grand-transcriptome. This tool allows users to search the database using gene identifiers to obtain the corresponding expression profiles across tissue types and developmental stages under different treatments, or to create an expression heatmap for genes of interest in selected organs, zones and for selected time points, with options for clustering genes and samples.

As an application example for this database, we examined the spatiotemporal expression of the maize *GRF* gene family during development, as this gene family plays an important role in determining organ size (Omidbakhshfard et al., 2015). Consistent with previous studies, most *GRF* genes exhibited high expression in the leaf division zone and showed a gradual decrease in expression along the developmental gradient to the mature zone, with exception of *ZmGRF4*, which displayed relatively stable expression among leaf zones and was reported to be insensitive to miR396a that negatively regulates *GRF* expression (Figure 7A) (Candaele et al., 2014). In addition, a downregulation in expression over time was also observed in the leaf division zone for the *GRF* genes differentially expressed across zones (Figure 7A), reflecting reduced cell division activity as time progressed. Interestingly, when comparing different leaf ranks, the average expression of *GRF* genes was higher in L15 and L12 compared with L18 and L9 after leaf appearance (Supplemental Figure 17A), which is in line with larger division zone size observed in L15 and L12 (Figure 2A), and suggests subtle regulation of *GRF* genes among different leaves. The expression patterns of *GRF* genes in the internode and ear were notably similar to those observed in the leaf (Figure 7B; Supplemental Figure 17B). Except for *ZmGRF4* and *ZmGRF10*, all *GRF* genes were downregulated over time during internode development. Additionally, differential expression of these *GRF* genes was also observed between the internode low zone and high zone, with higher expression in the low zone (Figure 7B). Despite a similar trend, the expression levels of several *GRF* genes vary across different organs. For instance, *ZmGRF9* exhibited higher expression in the internode compared to the leaf and ear, while *ZmGRF7* and *ZmGRF16* showed more pronounced expression in the ear (Supplemental Figure 18). This divergence in *GRF* gene expression suggests that different members may have specialized roles tailored to the unique growth needs of specific organs.

**Figure 7.**
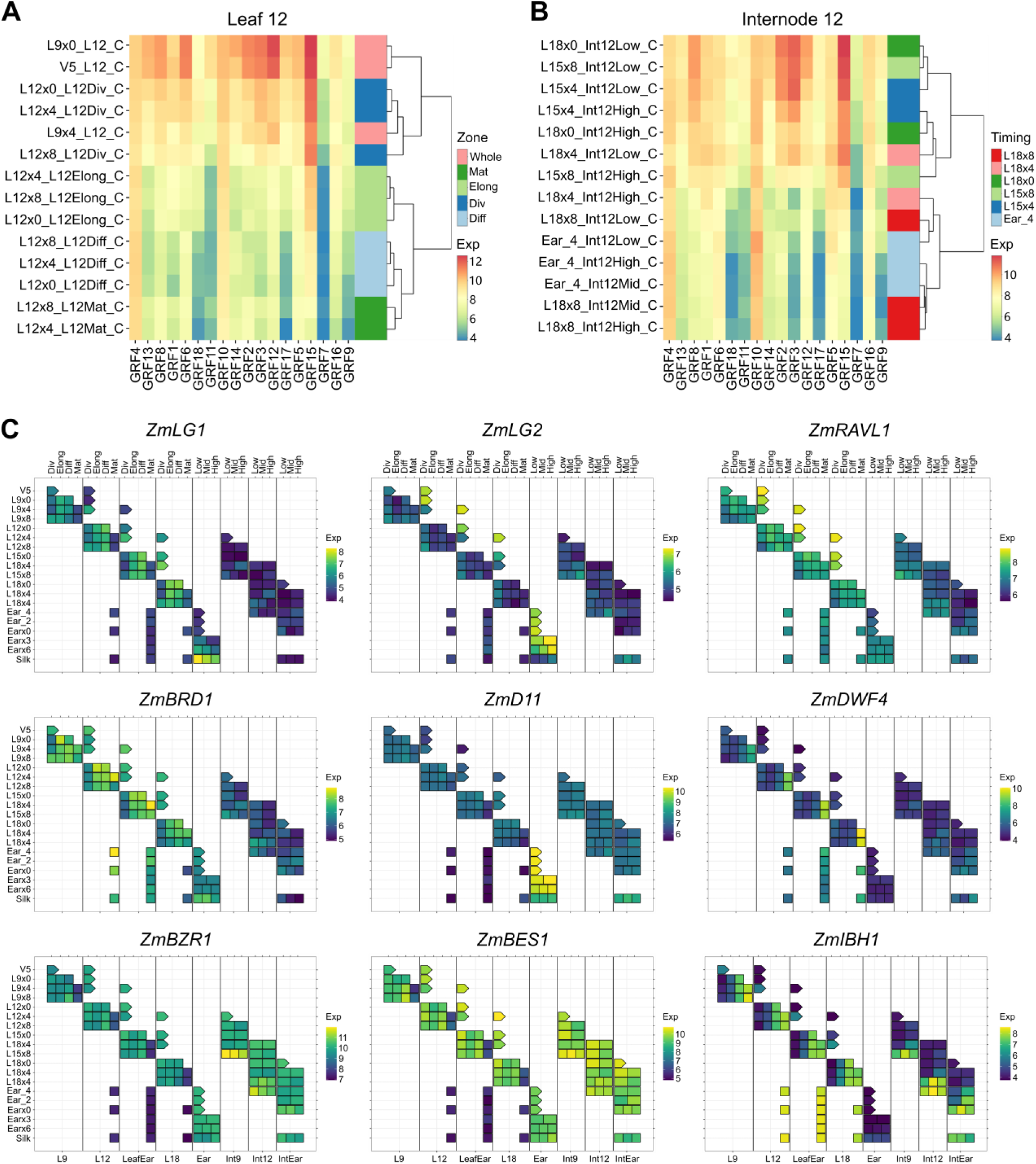
Spatiotemporal expression analysis of maize *GRF* family genes and leaf angle genes. **A-B)** Hierarchical clustering heatmaps of *GRF* genes across different zones of leaf 12 (A) and internode 12 (B) over time. **C)** Expression heatmaps of leaf angle genes during development. Gene expression values (Exp) are variance-stabilized transformed counts.

In addition to examining the *GRF* genes that regulate leaf size, we also investigated the spatiotemporal expression of genes involved in regulating leaf angle, an important trait of plant architecture affecting planting density and final yield. The key genes regulating ligule development, *ZmLG1* and *ZmLG2*, were primarily expressed in leaves and ear (Figure 7C). Interestingly, the expression of *ZmLG1* increased over time before leaf appearance and was higher in the elongation and differentiation zones during subsequent development (Figure 7C). Conversely, the expression of *ZmLG2* decreased over time and was more prominent in the leaf division zone, occurring earlier than *ZmLG1* both spatially and temporally (Figure 7C). This expression pattern, which was also observed in the ear (Figure 7C), aligns with previous studies showing that *ZmLG2* expression precedes that of *ZmLG1* in the ligule region, supporting that *ZmLG2* functions upstream of *ZmLG1* (Harper and Freeling, 1996; Walsh et al., 1998). Notably, *ZmRELATED TO ABI3/VP1RAV-LIKE1* (*ZmRAVL1*), a transcription factor downstream of *ZmLG1* that regulates leaf angle (Tian et al., 2019), exhibited early expression in the leaf similar to *ZmLG2* (Figure 7C), suggesting the presence of other upstream factors activating *ZmRAVL1* expression during the early stages of leaf development. ZmRAVL1 activates the expression of BR biosynthesis genes, such as *ZmBRASSINOSTEROID-DEFICIENT DWARF 1* (*ZmBRD1*), *ZmDWARF4* (*ZmDWF4*), and *ZmD11*, which when mutated all displayed altered leaf angle (Tian et al., 2019; Tian et al., 2024). The expression of *ZmBRD1* in leaf was observed to increase from the leaf division zone to the elongation zone, similar to that of *ZmRAVL1* (Figure 7C). Interestingly, *ZmDWF4* and its homolog *ZmD11* showed distinct expression patterns. *ZmD11* was expressed in leaf division, elongation and differentiation zones as well as in internodes and highly in the ear, while *ZmDWF4* was primarily expressed in the leaf mature zone (Figure 7C). This is consistent with the recent finding that *ZmD11* functions in maintaining BR levels for overall plant growth, while *ZmDWF4* specifically supports BR levels for leaf inclination (Tian et al., 2024). The key transcription factors *ZmBRASSINAZOLE-RESISTANT 1* (*ZmBZR1*) and *ZmBRASSINOSTEROID INSENSITIVE1-EMS-SUPPRESSOR1* (*ZmBES1*), two BR signaling genes, were expressed broadly from the leaf division zone to the differentiation zone (Figure 7C), whereas the negative regulator of BR signaling, *ZmILI1-BINDING BHLH1* (*ZmIBH1*), showed prominent expression in the differentiation and mature zone (Figure 7C), suggesting active BR responses mainly in the leaf growth zones, where they regulate leaf growth and leaf angle.

As cells grow to multiply and expand in size, they also need to acquire the specific functions and characteristics essential for their roles, a process known as cell differentiation. We examined the expression of vascular-related genes along the leaf growth gradient to investigate the processes of xylem and phloem differentiation. The maize ortholog of the Arabidopsis gene *LONESOME HIGHWAY* (*LHW*), a key transcription factor that controls initial processes in vascular development and xylem fate (Ohashi-Ito and Bergmann, 2007), exhibited the highest expression in the leaf division zone and gradually decreased towards the mature zone, indicating early vascular cells differentiation at the leaf base (Supplemental Figure 19). LHW forms heterodimeric complexes with TARGET OF MONOPTEROS 5 (TMO5) to regulate xylem cell differentiation (De Rybel et al., 2013). Interestingly, maize *ZmTMO5* and its paralogs showed divergent patterns along leaf zones. *ZmTMO5* exhibited relatively high expression in the leaf differentiation zone, while *ZmTMO5-LIKE2* (*ZmTMO5L2*) was predominantly expressed in the leaf elongation zone, and *ZmTMO5L3* in the leaf division zone (Supplemental Figure 19), implying distinct functions for the *TMO5* family members and a coordinated regulation of xylem differentiation along the leaf growth gradient. Notably, *ZmCYSTEINE PROTEASE5* (*ZmCCP5*), which is homologous to Arabidopsis *XYLEM CYSTEINE PEPTIDASE 1* (*XCP1*) involved in xylem tracheary element autolysis (Funk et al., 2002), was highly expressed in the differentiation zone (Supplemental Figure 19), suggesting fast formation of tracheids after cell expansion to facilitate water and minerals transport. Consistent with this, the expression of *ZmCINNAMOYL-CoA REDUCTASE1* (*ZmCCR1*) which is required for lignifying tracheary elements (Tamasloukht et al., 2011), was more pronounced in the differentiation zone and mature zone (Supplemental Figure 19). Similar to xylem development, the maize orthologs of early phloem development regulators in Arabidopsis such as *SUPPRESSOR OF MAX2 1-LIKE3* (*SMXL3*) and *SMXL4* (Wallner et al., 2017), were highly expressed in the leaf division zone and decreased as time progressed (Supplemental Figure 19), indicating early phloem differentiation at the leaf base during the early stages of leaf development. The expression of maize *ZmCALLOSE SYNTHASE2* (*ZmCALS2*), which is homologous to Arabidopsis *CALS7* involved in callose deposition in developing sieve elements (Xie et al., 2011), exhibited higher levels in the leaf growth zones compared with the leaf differentiation zone and mature zone (Supplemental Figure 19). Furthermore, the maize ortholog of Arabidopsis *NAC45* transcription factor associated with phloem enucleation (Furuta et al., 2014), *ZmNAC78*, showed higher expression in both leaf elongation zone and differentiation zone (Supplemental Figure 19). Additionally, an exonuclease encoding gene *ZmUTP-GLUCOSE-P-URIDYLTRANSFERASE* (*ZmUGU1*) which is homologous to Arabidopsis *NAC45/86-DEPENDENT EXONUCLEASE-DOMAIN PROTEIN 1* (*NEN1*) (Furuta et al., 2014), was predominantly expressed in the leaf differentiation zone (Supplemental Figure 19). In the mature zone, high expression of the sugar transport gene *ZmSWEET13a* was observed (Supplemental Figure 19), reflecting the maturation of phloem cells and the enhanced transport of photosynthetic products.

Taken together, this transcriptomic database enables the use of spatiotemporal gene expression data to better understand the regulation of gene families, signaling pathways, and developmental events during the maize shoot organ growth.

### Identification of organ-specific genes

Elucidating and characterizing organ-specific genes is crucial for comprehending the unique functions and regulatory networks that govern each organ. Using the gene expression atlas data of all the control samples, we identified 1,444 organ-specific genes based on the tau score as a specificity index (Kryuchkova-Mostacci and Robinson-Rechavi, 2017). Among these, 930 genes were identified as leaf-specific, while 238 and 276 genes displayed internode-and ear-specific expression, respectively (Supplemental Data Set 14). This aligns with previous studies that reported the largest number of organ-specific genes was observed for leaves in maize (Sekhon et al., 2011; Hoopes et al., 2019). GO enrichment analysis showed that the leaf-specific genes were predominantly involved in photosynthesis such as photosystem I and photosystem II assembly, chlorophyll biosynthesis, light reaction, C4 carbon fixation and shuttling, the Calvin cycle and redox regulation, highlighting the specialized role of leaves in photosynthesis. Several genes associated with lipid metabolism, including genes necessary for phospholipid biosynthesis, fatty acid synthesis and elongation, lipid desaturation and degradation were also found to be leaf-specific. Furthermore, some enrichment was observed among leaf-specific genes in genes related to secondary metabolic processes, including phenylpropanoid metabolism, terpene biosynthesis, coumarin metabolism and indole glucosinolate catabolism. In particular, a set of genes involved in responses to abiotic stimuli were identified as leaf-specific, indicating the important roles of leaves in environmental interactions.

The internode-specific genes were primarily enriched for genes involved in cell wall organization, phenylpropanoid metabolism and lignin metabolism, which contribute to stem strength. A number of genes encoding enzymes for primary and secondary cell wall biosynthesis, such as cellulose synthase, pectinesterase, polygalacturonase, xyloglucan endotransglucosylase, lignin-forming anionic peroxidase and laccase, were identified to exhibit internode-specific expression. Several expansin genes, which play critical roles in regulating cell wall enlargement in growing cells (Cosgrove, 2005), including *ZmEXPA10*, *ZmEXPA23*, *ZmEXPB7*, *ZmEXPB17, ZmEXPB22* and *ZmEXPB25*, were also among the internode-specific genes. Moreover, there was an internode-specific expression of genes related to hormone homeostasis, including *ZmETHYLENE INSENSITIVE3-LIKE6* (*ZmEIL6*) and *ZmEIL7*, two *EIL* transcription factors that function in ethylene signaling, *ZmISOPENTENYLTRANSFERASE 6* (*ZmIPT6*), a cytokinin biosynthesis gene, and a homolog of Arabidopsis *HISTIDINE-CONTAINING PHOSPHOTRANSMITTER4* (*AHP4*) involved in cytokinin responses, suggesting the role of ethylene and cytokinin signaling in internode development. Additionally, we also found several genes responsible for malate and aspartate transport among the internode-specific genes, indicating a role of the internode in C4-dicarboxylate transport.

The most prominent GO category enriched for the ear-specific genes was “regulation of transcription, DNA-templated”. Sixty-eight transcription factors were found to be ear-specific, of which many have been reported to be involved in inflorescence and flower development. For example, the TCP transcription factor gene *TEOSINTE BRANCHED 1* (*TB1*) and the HD-ZIP transcription factor gene *GRASSY TILLERS1* (*GT1*) have been demonstrated to affect axillary outgrowth, whereas the ERF transcription factor gene *BRANCHED SILKLESS*1 (*BD1*) controls the spikelet-pair meristem identity (Doebley et al., 1995; Chuck et al., 2002; Whipple et al., 2011). A total of 22 MADS-box genes were identified to be ear-specific, including *ZAG1*, *ZAG2*, *BEARDED-EAR1* (*BDE1*), *SILKY1* (*SI1*), *ZMM2*, *ZMM8*, *ZMM14*, and *ZMM16,* which are key regulators in the control of flowering time, determination of floral meristem identity and floral organ identity (Callens et al., 2018). Moreover, other potentially development-related transcription factor genes were ear-specific as well, including nine *HB*, six *YABBY*, three *GATA*, three *WRKY* and three *NAC* transcription factor genes. Besides transcriptional regulation, some enrichment was also observed for genes involved in several metabolic processes, including chitin catabolism, triterpenoid biosynthesis, trehalose biosynthesis and xyloglucan biosynthesis. Interestingly, among the ear-specific genes, we found two *ARGONAUTE* genes involved in RNA interference and *TS4*, which encodes a *mir172* microRNA (Chuck et al., 2007), highlighting the important roles of small RNA in ear development. Together, these identified organ-specific genes reflect the primary functions of each organ and provide a valuable resource for further research of shoot organ-related traits in maize.

### Transcriptomic analysis of different organs in response to drought stress

Drought is the most important abiotic stress inhibiting growth of maize plants, leading to a reduction of growth rate of individual organs (Verbraeken et al., 2021). To investigate the molecular mechanisms underlying the drought effects on maize organ development, transcriptomic comparisons were conducted between plants subjected to well-watered control and drought treatments starting at the V5 or V12 stage. Remarkably, the PCA showed that it was mainly PC11 (capturing 0.6% of variation) that discriminated the drought samples from control samples (Figure 8, A-B), indicating the differences along the development gradient were much more prominent than the differences between watering treatments. Both V5 and V12 drought treatments gave rise to higher values for PC11 (Figure 8B). The genes with positive loadings for PC11 were mainly involved in response to heat, response to water deprivation and response to abiotic stimulus, whereas the genes with negative loadings showed enrichment in terms linked to anion transport, catabolism and secondary metabolism, reflecting decreased metabolic and cellular activities under drought.

**Figure 8.**
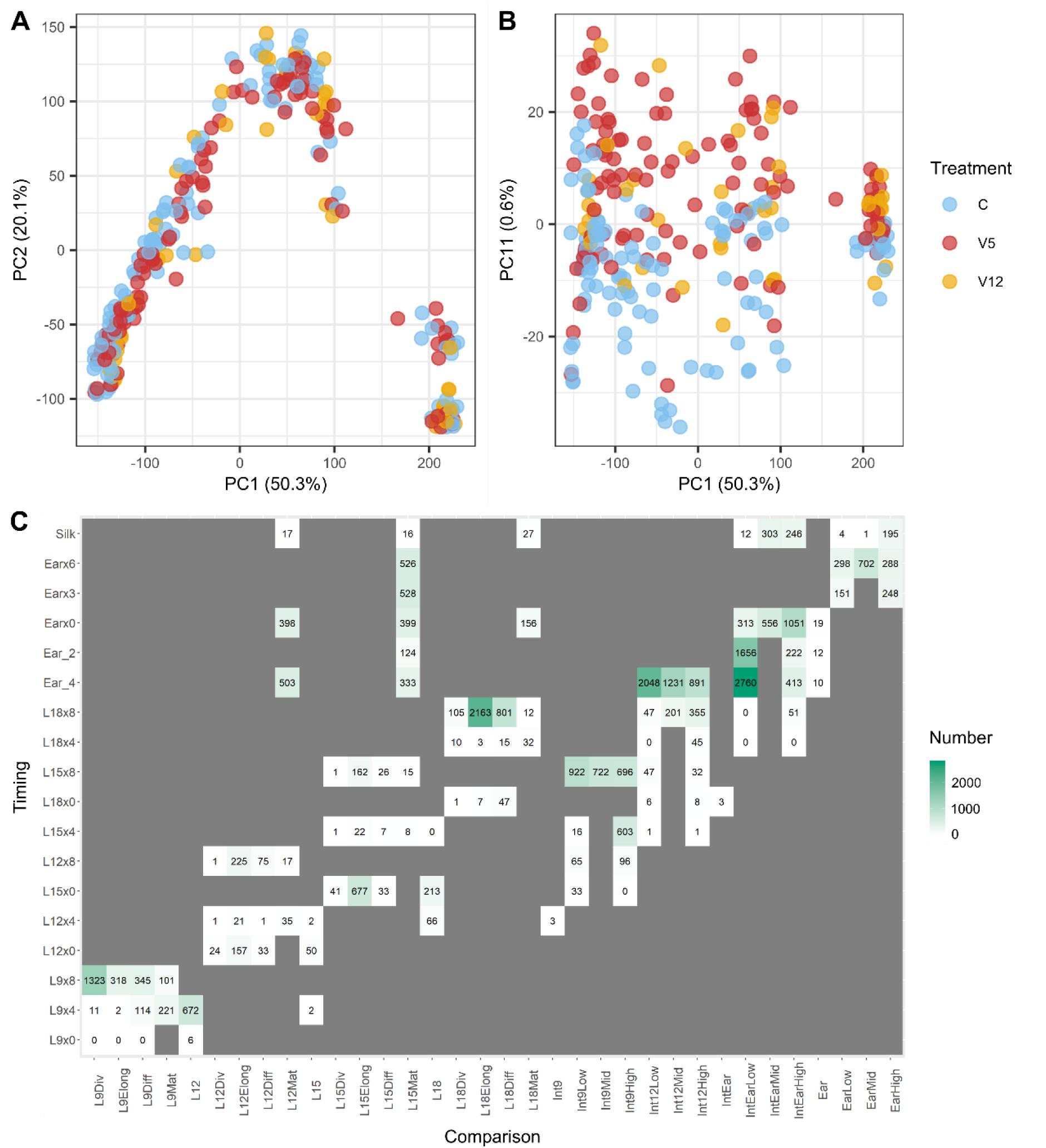
Transcriptomic analysis of gene expression in response to drought stress in leaf, internode, and ear. **A-B)** PCA of transcriptome profiles of all the control and drought samples. The percentage of variation among tissues explained by each principal component is displayed on the corresponding axes. The colors indicate different treatments. **C)** Number of DEGs in each comparison between V5-Drought and its corresponding control sample. The x-axis presents the organ together with zone; the y-axis presents the time points.

We identified DEGs between V5 drought and control conditions throughout leaf, internode and ear development (Figure 8C; Supplemental Data Sets 15-17). Since the V5 drought treatment started just before L9 emergence, we first investigated the DEGs in different zones of L9. At the time when L9 appeared, no significant DEGs were observed. As the soil was drying down, the amount of DEGs gradually increased with the duration and increasing intensity of the drought treatment. At four days after L9 emergence (L9×4), after five days of V5 drought treatment on average, very few genes were observed to be differentially expressed in the division zone and elongation zone, while a moderate number of DEGs were identified in the differentiation zone and mature zone (Figure 8C). In the leaf differentiation zone, several genes encoding chloroplast-localized proteins related to photorespiration, starch metabolism and detoxification of reactive carbonyl species were significantly upregulated. Conversely, the downregulated genes were enriched for genes involved in carbohydrate metabolism, including genes encoding hexokinase, inositol oxygenase, as well as soluble and cell wall invertases. In the mature zone, the most prominent enrichment among the upregulated genes was for genes involved in proline biosynthesis, highlighting the pivotal roles of proline in protecting the plants from drought (Hayat et al., 2012). Additionally, genes associated with starch and sucrose metabolism, including two β-amylase and three sucrose synthase-encoding genes, were found to be significantly upregulated in the leaf mature zone. At the later time point of L9 development (L9×8), the numbers of DEGs in the division zone and elongation zone were greatly increased (Figure 8C). The genes that were upregulated by drought in the elongation zone showed enrichment for genes involved in oxidation-reduction processes, water homeostasis and ABA metabolism, while genes related to adaxial/abaxial axis specification, wax biosynthesis and lipid metabolism were downregulated by drought. Remarkably, a large number of genes associated with the cell cycle were significantly upregulated in the division zone, while many genes involved in photosynthesis and cell wall organization were downregulated, which may result from delayed leaf development under drought stress. However, this was not observed in subsequently developing leaves (L12, L15 and L18) at 8 days after emergence, which might be due to the longer LED of these leaves compared with L9 (Verbraeken et al., 2021). By contrast, in these subsequent leaves, a large number of cell cycle genes were downregulated in the elongation zone, which hints at a spatially accelerated transition from cell division to differentiation by drought stress. During the reproductive stage, numerous genes were observed to be upregulated in the mature zone of these leaves (Figure 8C). Genes with functions in cell wall organization, cell tip growth and cell maturation were found among the upregulated genes, revealing delayed cell maturation in leaves under drought conditions, which may affect the energy supply for ear growth. When the plants reached silking stage, the number of DEGs was greatly decreased (Figure 8C), suggesting the drought-stressed leaves had fully matured.

The effect of V5 drought on the internode was evident from the increase in DEGs over time during the internode developmental process (Figure 8C; Supplemental Data Set 16). Genes involved in ABA biosynthesis and signaling, such as *ZmNINE-CIS-EPOXYCAROTENOID DIOXYGENASE3* (*ZmNCED3*), *ZmNINE COMPLEX PROTEIN2* (*ZmNCP2*) homologous to Arabidopsis *ABI FIVE BINDING PROTEIN3* (*AFP3*), and several *PROTEIN PHOSPHATASE 2C* (*ZmPP2C*) genes (*ZmPRH3*, *ZmPRH12*, and *ZmPRH13*), were significantly upregulated from the early stages (Supplemental Figure 20), indicating the crucial role of ABA in drought response and acclimation. Furthermore, the heat shock transcription factors *ZmHSF18* and *ZmHSF28* exhibited enhanced expression under drought (Supplemental Figure 20), aligning with previous studies demonstrating the importance of *HSF* genes in regulating drought response (Guo et al., 2016). Additionally, genes related to sugar metabolism and translocation were among the upregulated genes, including *ZmSWEET4a*, *ZmSWEET15b* and a sucrose synthase encoding gene (Supplemental Figure 20), highlighting the significance of soluble sugar content during drought response. Among the genes downregulated by drought, two genes encoding proline dehydrogenase that catalyze proline degradation were identified to be consistently downregulated in both the low zone and high zone of the internode (Supplemental Figure 20), pointing to the important role of proline in response to drought, not only in leaves but also in internodes. Compared to the early stages of internode development, a greater number of DEGs were observed in individual internode zones during the later stages (Figure 8C). In Int9, upregulated genes in both the low zone and high zone were significantly enriched for genes involved in starch metabolism, response to ABA, and response to water. Among these were the dehydrin encoding genes *ZmDHN1*, *ZmDHN2* and *ZmRD29B*, whose expression showed more significant upregulation during the late stage (Supplemental Figure 20), suggestive of a long-term protective role of dehydrin in response to drought. The downregulated genes in the high zone of Int9 exhibited enrichment for genes involved in cell wall organization and phenylpropanoid biosynthesis, suggesting reduced cell wall deposition upon drought stress in internodes. Notably, at the last time point of Int9 (T5), a number of cell cycle genes were found to be upregulated in the low zone of Int9, implying a prolonged duration of cell division in the internode under drought, which aligned with the upregulation of cell cycle genes in L9 of V5-Drought plants during the late stage of development (L9×8). Towards the reproductive stage (Ear_4), a large number of genes were differentially expressed in both Int12 and IntEar (Figure 8C). In both the low zone and high zone of Int12, genes associated with cell wall organization were significantly downregulated. In IntEar, genes involved in cell wall organization and photosynthesis were upregulated in the low zone, whereas the expression of genes governing cell cycle were repressed, suggestive of drought-induced inhibition of cell division and acceleration of cell maturation, which was also observed in L12, L15 and L18.

Besides leaf and internode, V5 drought also inhibited ear development, leading to reduced numbers of ear rows and spikelets per row (Verbraeken et al., 2021). The transcriptomic analysis showed that, upon the drought treatment, there was an increasing amount of DEGs over time before silking in ear tissues (Figure 8C; Supplemental Data Set 17). Before ear emergence, few DEGs were observed in the whole ear. Among them, *ZmBURP4*, which is homologous to Arabidopsis ABA-induced dehydration-responsive gene *RESPONSIVE TO DESICCATION22* (*RD22*) (Harshavardhan et al., 2014), was most strongly upregulated by drought (Supplemental Figure 21), implying an early accumulation of drought*-*induced ABA in ear. Expression of MADS-box genes *ZMM1*, *ZMM8*, *ZMM14* and *ZMM27* that are involved in flower development displayed a significant increase in expression under drought, while *TUNICATE1* (*TU1*), *ZmMADS68* and *ZmMADS73*, which are homologous to Arabidopsis *SHORT VEGETATIVE PHASE* (*SVP*) that acts as a floral repressor (Andres et al., 2014), were downregulated (Supplemental Figure 21), suggesting that drought stress may promote flowering (Takeno, 2016). However, the expression of *ZmMCT1*, a paralog of *TERMINAL EAR1* (*TE1*) that plays a role in spikelet pair meristem initiation and differentiation (Sun et al., 2024), showed significant decrease under drought conditions, implying reduced meristem activity and potentially impaired inflorescence development (Supplemental Figure 21). After ear emergence (Earx3 and Earx6), the transcriptomic changes between V5 drought and control were much more pronounced (Figure 8C). Genes associated with cellular responses to external stimuli, protein polymerization, anion transport, cell wall organization, and hormone-mediated signaling pathways were significantly upregulated. The hormone signaling category included genes responsible for ABA synthesis and degradation, jasmonic acid (JA) biosynthesis and cytokinin (CK) oxidation, suggesting the collaborative regulation of hormone activities in order to cope with the drought stress and to adjust growth and development. The downregulated genes in ear at both time points were primarily enriched for genes involved in the regulation of gene expression. Notably, 11 *GRF*s were observed to be significantly downregulated in at least one ear zone between control and drought samples, which indicated that drought stress restricts ear development by reducing cell proliferation. In line with this, an enrichment of DNA replication initiation and cell cycle genes was also observed among the downregulated genes. However, by the time of silk emergence, these cell cycle genes showed no obvious expression difference between drought and control conditions in the ear low zone and middle zone, consistent with a diminished cell division activity in the lower part of the ear at the silking stage. Overall, our investigations suggest that transcriptional regulation of gene expression underlies major reprogramming events in response to drought.

## Discussion

The growth of organs depends on the coordinated regulation of cell division and cell elongation, two fundamental processes that determine the cell number and cell size, respectively. These processes are generally considered to occur in a relatively sequential manner, as exemplified by the maize leaf where cell division takes place at the leaf base, forming a division zone, followed by a zone of cell elongation. In addition to this spatial separation, development is also highly dynamic over time, with growth zones changing in size as time progresses (Fournier et al., 2005). Although the growth zones in maize leaves have been extensively studied, the dynamic regulation of genes over time remains poorly understood. Moreover, different leaf ranks exhibit distinct developmental characteristics, which require precise gene regulation. The maize internodes exhibit similar developmental processes as leaves, but internode growth is more rapid compared to leaf growth (Verbraeken et al., 2021), implying distinct developmental regulation. In this study, we systematically analyzed the developmental characteristics of growth zones in maize leaves and internodes over time. We examined how gene expression changes over time in different developmental zones of leaves and internodes of different rank, and of the ear, under Control conditions and in response to drought.

### Cell division and elongation processes are highly conserved with different timing between leaf and internode

Using a new non-destructive method to estimate maize internode length, we observed a sigmoidal growth pattern in internodes similar to that in leaves (Voorend et al., 2014). The growth zones of internodes are closely reminiscent to those of growing maize leaves, characterized by a basal division zone followed by the elongation zone during development. These similarities indicate a shared fundamental growth regulation network in both organs, with growth processes building on the same foundation of cell division and pushing newly produced cells along the organ’s length axis while they elongate. In agreement with this, our transcriptome analysis revealed similar global gene expression patterns during leaf and internode development. In both organs, genes controlling the cell cycle and epidermis development exhibit highest expression in the meristematic region, with expression levels gradually decreasing towards mature sections, and progressively over time as the division zone becomes smaller. Supporting this, disruptions in genes regulating cell division often lead to abnormal development in both leaves and internodes. For instance, the maize *TANGLED1* (*TAN1*) gene is required for spatial control of cytoskeletal arrays associated with cell division, and the *tan1* mutants display shortened leaves and stunted growth (Martinez et al., 2020). Mutants of *ZmGIF1*, which encodes a protein that interacts with GRFs to form a functional transcriptional complex, develop small and narrow leaves as well as short internodes (Zhang et al., 2018). Compared with the dividing tissues, cells undergoing elongation in both leaves and internodes exhibit upregulation of genes involved in cell wall organization, photosynthesis, and active transport. Several genes related to BR signaling pathways were identified to be highly expressed in elongating cells, aligning with the crucial role of BR in cell elongation (Planas-Riverola et al., 2019). Mutants impaired in BR biosynthesis or signaling, such as *brd1*, *na2*, *bri1* and *bon1*, exhibit both shorter leaves and reduced internode lengths (Makarevitch et al., 2012; Kir et al., 2015; Best et al., 2016; Jing et al., 2024), demonstrating that conserved mechanisms regulate cell elongation across organs. These conserved pathways of cell division and cell elongation ensure that different plant organs develop within a similar framework. By adopting similar mechanisms for cell growth in various organs, plants can more efficiently utilize internal resources such as hormones and nutrients, reducing the need for different signaling pathways and regulatory factors.

Despite the highly conserved molecular mechanisms governing cell division and elongation, our kinematic analysis of division zone and elongation zone size in leaf and internode revealed significant differences in the timing of these processes between the two organs. Leaf elongation exhibits a steady state / linear growth phase during which the LER, division zone size and elongation zone size remain relatively stable and close to their maximum values (Schnyder et al., 1990; Ben-Haj-Salah and Tardieu, 1995). In internodes on the other hand, the elongation duration and matching steady state phase in individual internodes is much shorter than for comparable leaves. The internode division zone size is maximal very early on and then consistently diminishes throughout growth, remaining at about half its maximal size at the time of maximal elongation, in contrast to the leaf division zone, which remains close to its maximal size until the time point of maximal elongation, and only substantially decreases afterwards. The elongation zone size behaves similarly in both organs, waxing and waning together with the elongation rate. In leaves, fundamental growth processes are mainly characterized by a spatial separation, with cell division occurring at the base of the leaf and cell elongation occurring simultaneously in the area just beyond the division zone (Tardieu et al., 2000). In internodes however, the growth processes are more characterized by a temporal separation (although there is still spatial separation as well), with cell division mainly occurring early in internode growth and most of the cell elongation occurring afterwards. These distinct growth characteristics result in the leaf exhibiting a steadfast, balanced sort of growth by maintaining a long-term balance between cell division and cell elongation, allowing the leaf to steadily expand its surface area over time to maximize sun energy absorption. In contrast, internode growth has a more rushed nature, quickly producing cells followed by quickly expanding them, enabling the stem to quickly increase in height and strength to support newly formed leaves and reproductive structures. This growth pattern of the internodes may also be related to their stacked nature. The presence of an extensive zone of dividing, undifferentiated cells with weak cell walls in the lower part of an internode, while the upper part is already elongating and hence increasing the leverage effect that higher internodes and the associated leaves can exercise on the division zone below, might create a significant structural vulnerability. Therefore, the rapid transition from cell division to expansion in internodes likely serves to mitigate this weakness, ensuring mechanical stability and facilitating efficient vertical growth. Investigating the molecular mechanisms underlying this divergent timing program between leaves and internodes presents an intriguing area for further research.

### Spatiotemporal dynamics of gene expression coordinate cell growth and differentiation

The development of an organ requires an intricate balance between cell growth and differentiation. In maize leaves, the differentiation occurs progressively along the developmental gradient, resulting in the formation of distinct cell types. Through differential gene expression analysis among leaf zones and temporal stages, our transcriptomics data revealed the precise spatiotemporal regulation of cell differentiation parallel to cell growth. The expression of key transcription factors controlling stomatal differentiation, *ZmSPCH2* and *ZmMUTE*, was highest in the leaf division zone and significantly downregulated in the elongation zone, consistent with the observation that the earliest temporal stages of stomata exist only basally (McKown and Bergmann, 2020). Furthermore, the differentiation activity of stomata diminishes over time, as indicated by the temporal downregulation of these transcription factors. A similar expression pattern was observed for genes related to trichome development, such as *ZmOUTER CELL LAYER4* (*ZmOCL4*) and *ZmHOMEODOMAIN GLABROUS11* (*ZmHDG11*) (Nakamura et al., 2006; Vernoud et al., 2009), implying early trichome differentiation alongside rapid cell progressive differentiation, as reflected by the spatiotemporal expression of key genes involved in xylem and phloem development. Both early differentiation genes for xylem and phloem, such as *ZmLHW* and *ZmSMXL3*/*4*, exhibited their highest expression levels in the leaf division zone, while genes related to further vascular cell specialization, e.g. *ZmCCP5* and *ZmUGU1*, were most highly expressed in the leaf differentiation zone where cells cease growth. This coordination between cell growth and cell differentiation ensures that each cell type achieves the appropriate quantity and size while acquiring the characteristics and functions necessary for their specific roles.

The regulation of cell growth and differentiation is governed by various signaling pathways. Through mutant identification and analysis of genetic and molecular interactions, many components of these signaling pathways have been uncovered and their regulatory relationships established. However, our current understanding of these signaling pathways is often built on a static network representation and dynamic information is comparably lacking. Through spatiotemporal transcriptome analysis, deeper insights can be gained into how these pathways function in a developing multicellular structure. Our high-resolution transcriptomic map shows specific transcriptional programs controlling when and where key players are being produced, as exemplified by the analysis of the spatiotemporal expression of leaf angle genes. When examining the expression of signaling pathway members from a broad spatiotemporal perspective, more complex regulation might emerge. For example, we observed that the expression of *ZmLG1* and its downstream target *ZmRAVL1* was not temporally synchronized. Additionally, the BR biosynthesis genes *ZmDWF4*, *ZmD11*, and *ZmBRD1* were all found to be targeted by ZmRAVL1, but they showed distinctly different expression patterns within the leaf, which implying a complex spatiotemporal regulation of BR synthesis in maize leaves.

### Differential gene expression regulates development across different organs and within the same organ

Different plant organs exhibit unique morphological features and perform distinct functions, requiring specialized developmental processes. In our transcriptome atlas, organ-specific genes for leaves and internodes were predominately enriched for genes involved in photosynthesis and cell wall organization, respectively, aligning with their primary functions of these organs: photosynthesis in leaves and structural support in internodes. Although photosynthesis-related genes were also observed to be upregulated in the internode as cells matured, the expression of these genes was much lower in the internode than in the leaf. It has been reported that stem photosynthesis can contribute significantly to carbon assimilation and stress tolerance in a wide range of woody plants (Simkin et al., 2020). In maize however, the role of stem photosynthesis for overall plant development remains unclear. As internodes of maize are completely enclosed by leaf sheaths, this reduces their ability for light capture and gas exchange. In contrast, genes involved in biosynthesis of lignin, which imparts significant strength and rigidity to the cell wall (Liu et al., 2018), were expressed at relatively higher levels in the internode compared to the leaf. This is consistent with a recent study showing that the expression of most lignin biosynthetic genes is higher in root and stem compared to leaf and tassel (Yang et al., 2024). In the ear, a notable enrichment of transcription factors was identified among ear-specific genes, aiding in the discovery of new regulators of inflorescence development. These organ-specific genes provide a valuable resource for engineering targeted traits in maize. For example, internode-specific genes hold promise for developing short-stature maize varieties with enhanced tolerance to lodging under high wind conditions. A previous study demonstrated that targeted suppression of gibberellin biosynthetic genes *ZmGA20ox3* and *ZmGA20ox5*, both of which exhibit higher relative expression levels in vegetative tissues and lower levels in reproductive tissues, can produce a short-stature maize ideotype by inhibiting elongation of internodal cells (Paciorek et al., 2022). In our analysis, we identified six internode-specific expansin genes, which are likely involved in cell wall loosening and expansion in internode. Additionally, two *SMALL AUXIN-UP RNA* (*SAUR*) genes, *ZmSAUR3* and *ZmSAUR56*, were found to be specifically expressed in the internode. *SAUR* genes play a central role in auxin-induced growth (Stortenbeker and Bemer, 2019). Notably, the short-stature *brachytic2* (*br2*) maize mutants exhibit defects in auxin transport within the internode (Multani et al., 2003), highlighting the potential to engineer the expression of these *SAUR* genes to develop short-stature maize varieties.

The understanding of the development of each type of organ, such as maize leaves, often comes from studying a single leaf. However, different ranks of maize leaves exhibit distinct morphological characteristics. The seedling leaves and flag leaves are typically shorter than the middle leaves, and their leaf angle differs. Additionally, distinct leaves also vary in developmental dynamics, as shown by a decrease in LER and an increase in LED from L9 to L18 (Verbraeken et al., 2021). Although different ranks of leaves exhibited a highly similar global expression pattern, subtle differences in gene expression regulation were observed. The expression levels of *GRF* genes were higher in the middle leaves compared with lower and upper leaves. Moreover, the BR biosynthesis gene *ZmDWF4* showed an increase expression from L9 to L18, which is in line with the observation that *ZmDWF4* expression has the highest correlation with upper leaf angle (Tian et al., 2024). Understanding how these genes are differentially expressed in different leaves remains an interesting question. In addition to leaves, different internodes of a maize stem also exhibit distinct characteristics, differing in length, thickness, anatomy, chemical composition, and digestibility (Boon et al., 2008). The precise regulation of traits across different organ ranks is crucial for optimizing maize architecture. For example, the ideal maize architecture involves compact and small upper leaves with a smaller leaf angle, while the middle and lower leaves are ideally longer and more spread out, to facilitate maximum light energy absorption. Identifying genes associated with variations among different organ ranks will improve our understanding of plant development and enable more precise genetic engineering, contributing to developing maize varieties with optimized overall characteristics.

### The responses to drought stress involve tissue- and developmental stage-specific reprogramming of gene expression

Chronic drought, here in the form of the V5-Drought treatment, affects internode growth in a similar way to leaf growth, reducing elongation rate, compensated partially by an increased elongation duration, resulting in a net reduction of final length which is less severe than expected from the reduction in elongation rate alone. This effect is known in leaves as ‘compensation’ (Avramova et al., 2015; Hisanaga et al., 2015; Avramova et al., 2017; Nelissen et al., 2018; Verbraeken et al., 2021; Tabeta et al., 2022; Van Hautegem et al., 2025). The relative effect of V5-Drought on growth traits is much stronger in internodes than it is in leaves. In terms of final length, internodes display a reduction between 19.51% and 34.35%, while leaves only display a reduction between 5.2% and 12.1% (Verbraeken et al., 2021). Previously, we described a decreasing trend in LER and an accompanying increasing trend in LED over leaf rank under well-watered conditions, resembling the effects of V5-Drought (Verbraeken et al., 2021). We hypothesized that this effect might be due to a reduction in carbohydrate and water supply to the growing leaf due to increased competition with the growing stem (Verbraeken et al., 2021). Looking now in detail at this competing organ, the stem, we see an opposite trend over rank: elongation rate and duration both display an increasing trend with internode rank, resulting in a significant increase in internode length with rank. This implies the internode sink strength increases with rank, supporting the aforementioned resource reduction hypothesis for leaves of higher ranks. Further evidence that growing internodes receive the bulk of the plant’s resources during their growth period is the strong effect of V5-Drought on the growth of internodes 9, 12 and 15. Growing organs do not easily shut down their growth completely (Hisanaga et al., 2015; Tabeta et al., 2022), and when growing organs are limited in the resources they receive, any reduction in resource supply reduces the growth of the strongest sink organs first, before shutting down the growth of the weakest sink organs (Verbraeken et al., 2021).

When grown under drought conditions, plants attempt to adjust their adaptive mechanisms by reprogramming their gene expression in various organs and tissues. In our study, the dynamic changes of gene expression in response to drought treatment were investigated during leaf, internode, and ear growth, revealing a tissue-specific response to drought in terms of organ and developmental change. More differentiated tissues tend to exhibit stronger responses to drought compared to the younger tissues. For instance, in L9, which was the earliest leaf subjected to drought stress, a higher number of DEGs were observed in its differentiation and mature zones compared with the fast-growing division and elongation zones after five days of drought treatment. Additionally, very few DEGs were identified in the early temporal stages of both internode and ear tissues, followed by a progressive increase in DEGs over time, indicating that rapidly dividing internode and ear cells were minimally affected by drought. Our results are consistent with transcriptomic profiling of soybean seed development, which demonstrated that the influence of drought is little at the early stage of seed development and increases during the later stages (Tang et al., 2023). This relatively smaller drought response in rapidly growing cells enables them to recover quickly upon rewatering, as was seen for the leaf (Van Hautegem et al., 2025).

With the progression of organ development, more drought-induced DEGs were observed, including stress response genes and developmental genes. Genes related to ABA metabolism were found to be upregulated in all three organs, including the transcription factor *ZmNCP2* and the *ZmPP2C* genes. This indicates that ABA-mediated signaling is a common drought stress response mechanism across different organs, which is supported by a previous study showing that the ABA responsive element (ABRE) binding site was significantly enriched in the promoters of common drought-responsive DEGs in maize ear, kernel and ear leaf (Wang et al., 2019a). Numerous development-related genes were also found differentially expressed in response to drought. In internodes, a dramatic downregulation of cell wall-related genes was observed in the late stages of internode development, which is in agreement with previous studies showing that drought stress affects cell wall rigidity and can influence the mechanical strength of the stalk, leading to lodging and subsequent grain yield losses (Flint-Garcia et al., 2003; Kumar et al., 2021; Wang et al., 2023). In the ear, transcription factors that regulate cell division and cell cycle genes were significantly downregulated upon drought, consistent with the field-based transcriptomic studies on the maize ear in response to drought and explaining observations of retarded ear growth under drought (Danilevskaya et al., 2019; Wang et al., 2019a). It is noteworthy that in the latter study, cell cycle and DNA replication genes were enriched in the set of genes upregulated under drought during the late stages of ear development, associated with delayed ear growth under drought. In our transcriptomic data, the upregulation of cell cycle genes under drought in the late stages of development was observed in leaf L9 (L9×8) and internode Int9 (T5), but not in other leaves, internodes, or the ear, which might be due to developmental stage differences at the corresponding sampling time points for each organ. These findings suggest that the transcriptional response to drought is strongly influenced by the developmental stage of the tissue at the time of sampling.

In summary, our dataset provides a high-resolution spatiotemporal atlas of gene expression in developing maize shoot organs throughout plant development under both well-watered and drought conditions. The identification of organ-specific, stage-enriched, and drought-responsive genes offers a valuable foundation for future efforts to uncover key factors contributing climate-resilient maize varieties.

## Materials and methods

### Experimental setup

Five experiments were conducted in the Phenovision plant phenotyping platform (Verbraeken et al., 2021). As the first four experiments (experiments 1-4) in the periods between March-May and September-November in the years 2015 and 2016 are the same experiments detailed by Verbraeken et al. (2021), we refer to the materials and methods of Verbraeken et al. (2021) for phenotyping platform details and experimental conditions. The fifth experiment (experiment 5) was run in the period January-March 2017, occupying three lines of the Phenovision platform (84 positions in total, 78 experimental and 6 border), and otherwise copying the conditions of the first four experiments. All plants were of the stiff-stalk maize (*Zea mays* L.) inbred line B104. Plants were automatically watered each day, receiving water up to a specified gravimetric soil water content. The target soil water contents were 2.4 g water*g^-1^ dry soil for the Control treatment, at a soil water potential of around −10 kPa, which is close to field capacity, and 1.4 g water*g^-1^ dry soil for V5-Drought and V12-Drought treatments, corresponding to a soil water potential of around −100 kPa. All drought-treated pots started at the well-watered target soil water content and were switched to drought conditions at either the V5-stage (vegetative stage drought) or the V12-stage (reproductive stage drought). It took several days for the soil to dry down below the target soil water content and for the pots to receive daily watering again. Experiments 1-4 featured Control, V5-Drought and V12-Drought treatments, experiment 5 featured only Control and V5-Drought treatments.

A sixth experiment was performed for the internode cell length measurements. These plants were not grown on the Phenovision phenotyping platform but were grown in pots, under greenhouse conditions (daytime 06:00-22:00, daytime temperatures of 26°C and nighttime temperatures of 22°C, supplemental lighting to maintain light intensity above 150 μmol photons*m^-2^*s^-1^ during the daytime) and were manually kept well-watered. These plants were grown in two batches, the first batch grew in September-October 2019, the second batch in January-February 2020.

### Destructive internode sampling

The internode sampling scheme was identical in the experiments 2-4 and is detailed in Supplemental Table 1. Separate plants were sampled for transcriptome analysis on the one hand and destructive phenotyping on the other hand. Three replicates were taken for each sample per experiment, resulting in 9 replicates in total. 129 internodes at positions 9, 12 and 15 or at the ear position were sampled (54 Control, 54 V5-Drought, 21 V12-Drought) from 87 plants (36 Control, 36 V5-Drought, 15 V12-Drought) per experimental repeat for destructive phenotyping, and 129 internodes from 87 plants per experimental repeat were sampled for transcriptomics. The internodes were sampled by cutting through the middle of the nodes above and below the internode. For experiments 2-4, the internodes intended for destructive phenotyping were stored as a whole in an ethanol:acetic acid (3:1) mixture and the lengths of the sampled internodes 9, 12, 15 and the ear internode were later determined by measuring end-to-end.

In experiment 5, 52 plants in total (28 Control, 25 V5-Drought) were dissected between the V5-stage and the V18-stage. Upon dissection, the lengths of all internodes, starting from the 6th, were immediately measured from the middle of the lower flanking node to the middle of the upper flanking node. By measuring plants up to the V18-stage, the growth of the internodes up to internode 15 was recorded.

### RGB imaging setup and internode length estimation method

Details of the Phenovision imaging setup can be found in Verbraeken et al. (2021). Only the frontal RGB camera, with an image plane parallel to the background screen and the plane formed by the maize leaves is relevant for this study. This camera was an 11 megapixel Allied Vision Technologies Prosilica GE4000C (Allied Vision Technologies GmbH, Germany) camera equipped with a Canon EF 24 mm f/1.4L II USM lens (Canon Inc., Japan). Images were stored in JPEG format with a resolution of 2673*4009 pixels. The camera setup was calibrated with a conversion factor between image measurements in pixels and real-world measurements at the average location of the plant stem of 0.674 mm*pixel^-1^ (details in Verbraeken et al. (2021)).

Fournier and Andrieu (2000) show how internode lengths can be determined from the difference in leaf collar height between the leaves attached below and above the internode. The relationship between internode length and collar heights depends on the lengths of the leaf sheaths as well, as visualized in Figure 1A.

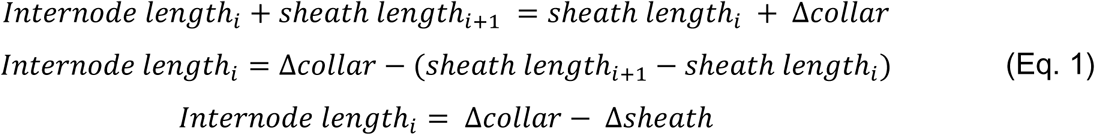

A ground truth internode length dataset was constructed for internodes 9, 12 and 15 under Control, V5-Drought and V12-Drought conditions using the internode length measurements from the destructive internode sampling for phenotyping performed in experiments 2-4 and experiment 5, described above. Leaf collar heights were determined from the images of the destructively sampled maize plants on their day of sampling, by determining the position of each visible leaf collar relative to the soil level, using an ImageJ macro (Supplemental Figure 1) (Schneider et al., 2012). These values were translated from pixel values to real world (mm) measurements using the aforementioned conversion factor. The resulting dataset contained 370 internode lengths (120 for internode 9, 145 for internode 12, 105 for internode 15) from 221 different plants with image-based collar height values. Collar i+1 was visible for 163 of the ground truth internodes and not visible for the remaining 207 internodes. The ground truth internode length data, Δcollar data, and the number of replicates for each of the combinations of factors can be found in Supplemental Data Set 1.

The leaf sheath lengths and resulting Δsheath could not be derived from the images and were not measured destructively for the ground truth plants from experiments 2-4, only in experiment 5. Instead, Δsheath was estimated as the difference between the ground truth internode lengths and their matching Δcollar values (Δ*sheath* = Δ*collar* − *Internode length*_*i*_). The distributions of the Δsheath values per internode rank and watering treatment are shown in Supplemental Figure 2A. The relation between internode length and Δcollar shown in Supplemental Figure 2B indicates that changes in Δcollar can be attributed entirely to changes in internode length when collar i+1 is visible, or in other words, that Δsheath remains constant after emergence of collar i+1.

As a confirmation, in experiment 5, sheath lengths and Δcollar were measured manually for 52 destructively sampled plants to assess the relationship between the length of sheath i+1 and Δcollar, for internodes i=9, i=12 and i=15 in the Control and V5-Drought watering treatments, at several time points both before and after the emergence of collar i+1 (Supplemental Figure 2). Linear models for sheath i+1 length as a function of Δcollar for visible collars i+1 (Supplemental Figure 2) indicate that there is indeed no more sheath elongation occurring once collar i+1 is visible, indicating Δsheath remains constant after emergence of collar i+1. The mean Δsheath values per internode rank and watering treatment from the ground truth dataset (Supplemental Data Set 1) were therefore used as constant Δsheath values in the estimation of internode lengths whenever collar i+1 was visible, in the form of 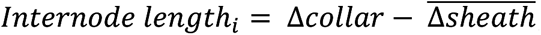, allowing the estimation of internode growth rates over time in setups where the leaf sheath lengths could not be measured.

For the cases in which collar i+1 was not visible on the image, an indirect internode length estimation method was developed for internodes 9, 12 and 15 in the situation that leaf collar 10, 13 or 16, respectively, was not visible on the image. The length of the internode of interest was predicted using linear models based on the Δcollar value of the highest visible pair of leaf collars, without going below leaf collar i-3 (i.e., not using a leaf collar below leaf 6, leaf 9 or leaf 12 for the length of internode 9, 12 and 15 respectively). So, for internode 12 this meant the length was estimated based on the difference in collar height between leaves 12 and 11, 11 and 10 or 10 and 9, depending on which was the highest visible collar. The relationship between these alternative Δcollar values and the internode length was determined using linear modelling (Supplemental Data Set 1).

### Internode growth trait analysis

A total of 14 plants that were not destructively sampled were selected for analyzing the growth of internodes 9, 12 and 15 using image-based internode length estimation. Six of these plants were subjected to the Control treatment (2 plants each in experiments 2, 3 and 4) and eight plants were subjected to the V5-Drought treatment (two plants in experiment 2, 3 plants each in experiments 3 and 4). The plants were imaged daily, allowing daily estimation of the lengths of internodes 9, 12 and 15 as detailed above, resulting in internode length time series. A beta-sigmoid growth function (Eq. 2; Supplemental Figure 5) (Yin et al., 2003; Voorend et al., 2014) was fitted to the length over time for each individual internode, using a nonlinear least squares approach (nls function (Bates and Watts, 1988) in R 3.6.3 (R Core Team, 2013)).

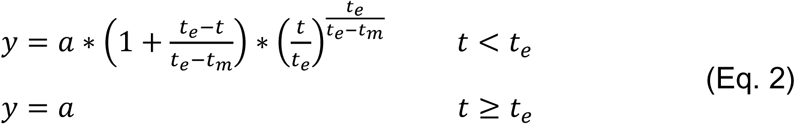

With *y* the internode length, *t* the plant age, *a* the final internode length, *t_e_* the plant age at the end of internode elongation and *t_m_* the plant age at the moment of maximal internode elongation.

The growth traits of interest were final internode length, elongation duration and mean elongation rate. The final internode length was determined directly during curve fitting, as a parameter in the growth function. Elongation duration was determined as the time between the internode reaching 10% of its final length (*t*_10_) and the end of internode growth (*t*_e_). The first derivative of the beta-sigmoid growth function gives an estimate of the internode elongation rate at any given time (Supplemental Figure 5). The mean elongation rate was determined between *t*_10_ and *t*_e_.

For the statistical analysis of the growth traits between the different internode ranks and treatments, a linear mixed-effect model (lmer function, lme4 package (Bates et al., 2015) in R 3.6.3 (R Core Team, 2013)) was used, featuring internode rank, treatment and the interaction as fixed effects and experimental repeat as a random effect. Significance of effects was tested using the Wald chi-squared test (car package) (Fox and Weisberg, 2018) and post-hoc general linear hypothesis tests (glht function, multcomp package (Hothorn et al., 2008) in R 3.6.3 (R Core Team, 2013)) were used to test for significant differences between pairs of internode ranks and for treatment effects within internode ranks. The p-values resulting from the post-hoc tests were corrected for multiple testing using Holm p-value correction (Holm, 1979).

### Destructive leaf sampling and cellular analysis

To analyze the leaf growth zone, plants were sampled at the day of leaf appearance, 4 days and 8 days after leaf appearance for leaves 9, 12, 15 and 18 under Control and V5-Drought conditions. The basal 14 cm of these leaves, containing the growth zone, was divided into pieces of 1 cm long and these pieces were stored in an ethanol:acetic acid (3:1) mixture in multiwell plates. These leaves were later microscopically analyzed to determine the cell length pattern of the epidermal cells throughout the growth zone and determine the delineation of the division zone and elongation zone in the leaves at these time points. The samples for leaves 12, 15 and 18 were taken during experiment 1, the samples for leaf 9 were taken during experiment 4.

For each 1 cm long sample, a series of images along the length of the sample was taken using the automated moving stage of a Zeiss Axiovert microscope (Carl Zeiss Microscopy, Jena, Germany), with each image covering a length of 671.84 µm. The length of 20 random cells was measured on each image using ImageJ (Schneider et al., 2012) and the distance to the leaf base, along the leaf length axis, was calculated. The position of the ligule along the leaf length axis was also determined on these images in order to differentiate between leaf sheath and leaf lamina (blade). Maize leaf growth is characterized by two distinct growth zones, one for the leaf sheath and one for the leaf blade, separated by the ligule (Sylvester et al., 1990). The growth zone is made up of a basal division zone and a subsequent elongation zone (Nelissen et al., 2013). The separate growth zones for the leaf sheath and leaf blade were especially noticeable in the cell length profiles for leaf 9 and leaf 18 at 8 days after leaf emergence, by which stages the leaf sheath had started to elongate (Supplemental Figure 6). The leaf blade was the focus of this study.

The cell length measurements were binned per 5 mm distance to the leaf base (Supplemental Figure 7). To determine the end of the division zone, the bin with the lowest mean cell length was determined. Then t-tests were performed to check for significant differences in cell length between the bin with the lowest mean cell length and the subsequent bins. If five bins in a row displayed a significantly different cell length from the bin with the lowest mean cell length, the most basal of these five bins was taken as the end of the division zone and the start of the elongation zone.

To determine the end of the elongation zone, a similar method was used but in the other direction. The bin with the highest mean cell length was determined and t-tests were performed to check for significant differences in cell length between the bin with the highest mean cell length and the preceding bins. If five bins in a row displayed a significantly different cell length from the bin with the highest mean cell length, the most distal of these five bins was taken as the end of the elongation zone.

This resulted in a collection of 65 replicates in total, for the various leaf ranks, time points and treatments. These data were matched with the mean macroscopic leaf growth traits for each leaf rank and treatment (LED and FLL) and with each sampling time point within rank and treatment to extract the immediate LER and estimated length at time of sampling from growth curves determined earlier by Verbraeken et al. (2021). The resulting dataset is included in Supplemental Data Set 3.

The effects of leaf rank, watering treatment and time after leaf emergence on the size of the lamina division zone and lamina elongation zone were analyzed using three-way ANOVA and effect significance was tested using an F-test, using the car package (Fox and Weisberg, 2018) in R version 3.6.3 (R Core Team, 2013). This was followed by a post-hoc Tukey’s Honestly Significant Difference test to test for significant pairwise differences between factor levels.

### Internode cell length sampling, cell length analysis and linking zone sizes to internode length

Two batches of plants were grown for determining cell length profiles, collectively referred to as experiment 6. Eighteen plants were sampled in the first batch, three plants per V-stage for V10 to V15. Seven plants were sampled in the second batch, of which two plants were sampled at the V12-stage, four plants were sampled at the V13-stage and one plant was sampled at the V14-stage. Leaves were stripped from these plants to reveal internode 12. The length of internode 12 was measured and an imprint of the internode epidermal cells over the entire internode length was taken by applying translucent nail varnish to the internode epidermis, waiting for the varnish to dry and then applying translucent tape to the varnish. The varnish stuck to the tape when it was lifted. The tape was then applied to a microscopy slide and the imprints of the epidermal cells in the varnish were visible under an optical microscope (Axio Imager; Carl Zeiss Microscopy, Jena, Germany). One cell file along the internode length axis was selected and the length of all cells on this file were measured using the AxioVision software (Carl Zeiss Microscopy, Jena, Germany), leading to a cell length profile along the internode length axis. In practice, this method failed to produce useful imprints for small internodes (< 10mm), and as a result the cell length profile for the samples at V-stages 10 and 11 and three of the samples at V-stage 12 could not be determined. Another sample at V-stage 13 was damaged and the cell length profile could not be determined. In total 15 cell length profiles were determined (2 at V12, 6 at V13, 4 at V14 and 3 at V15). These data covered most of the internode 12 growth timespan (Supplemental Figure 9).

The division zone and the elongation zone were determined from these cell length profiles using methods derived from those used for determining growth zones in juvenile maize leaves (Nelissen et al., 2013). A local polynomial smoothing was performed on the cell length profiles, to determine the average cell length along the internode length (locpoly function, 2^nd^ degree local polynomial, kernel smoothing bandwidth of 4 mm, KernSmooth package (Wand and Jones, 1994), for R (R Core Team, 2013)). The end of the elongation zone was defined as the position at which the smoothed cell length reached 90% of its maximal value observed within the basal 40 mm of the internode. The extra condition to use the maximal value within the basal 40 mm of the internode was added as there were some cases where cell length would plateau and then have an extra peak close to the top of the internode, resulting in the end of the elongation zone being placed very close to the top of the internode, including a long plateau with constant cell length (Supplemental Figure 10). The end of the division zone could not be determined by visualizing the nuclei as in Nelissen et al. (2013) as cell imprints were used, and instead had to be estimated from the cell length data. First, the start of the elongation zone was determined as the location where the first derivative of the local polynomial smoothing curve fit to the cell length profile exceeded 1 µm*mm^-1^, indicating cell elongation. In leaves, a gradual transition zone has been observed, where both elongating cells and dividing cells coexist (Nelissen et al., 2018), indicating the end of the division zone is located further from the base than the start of the elongation zone. 95% of cells located more basally than the start of the elongation zone had cell lengths below 40 µm, thus the location where the local polynomial smoothing indicated that cell length reached 40 µm was taken as the end of the division zone.

For each individual internode, the cell lengths and their associated positions along the internode length were bootstrapped 10,000 times. This resulted in distributions of estimates for the position of the end of the division zone, the end of the elongation zone and the length of the elongation zone, allowing the calculation of 95% confidence intervals for these traits. By combining these distributions per individual internode at the V-stage level, mean division zone and elongation zone sizes and their 95% confidence interval intervals were determined for each V-stage.

To investigate the behavior of the division zone and elongation zone over time, first the relationship between zone sizes and internode length was determined, by linear interpolation of the size of the zones in function of internode length between the V-stages, (Supplemental Figure 11). The resulting relationship is a piecewise linear function (Eq. 3).

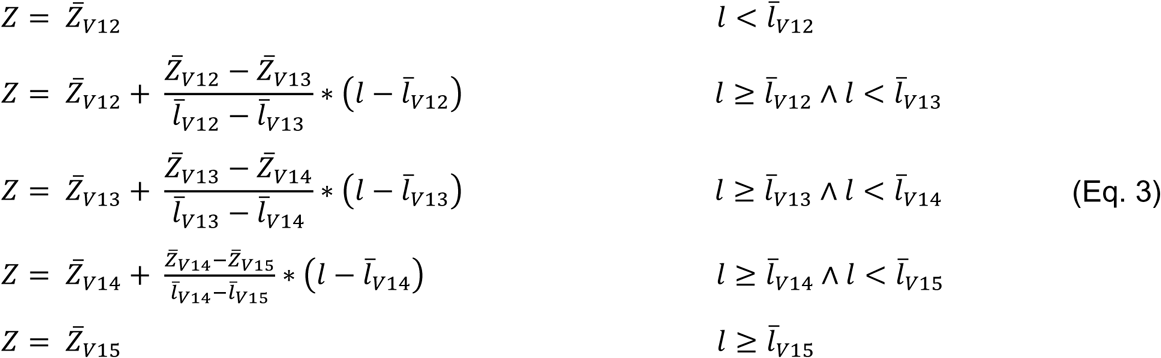

With *Z* the zone size (division zone or elongation zone), *Z̅*_*Vx*_ the mean zone size at V-stage *x*, *l* the internode length and *l̅*_*Vx*_ the mean internode length at V-stage *x*.

Second, to determine the relationship between internode length and time, the mean internode 12 growth curve under Control conditions was determined by taking the mean value of the beta-sigmoid growth function (Eq. 2) parameters determined for the six separate internode 12 length time series. By entering the mean parameter values into the beta-sigmoid growth function, a mean internode 12 growth curve could be determined, estimating internode 12 length as a function of time. By combining the internode length over time estimated from the beta-sigmoid growth curve, with the relationship between division zone and elongation zone size and internode length, the sizes of the division zone and elongation zone were estimated over time.

### Plant material collection for RNA sequencing

For all experiments, the inbred line B104 was used and grown in the Phenovision plant phenotyping platform. The plants were grown under well-watered conditions or subjected to drought treatment beginning at either the V5-stage or the V12-stage (Verbraeken et al., 2021). Different zones of leaf, internode and ear at distinct time points were sampled for RNA extraction and subsequent sequencing. A concise description of the materials collected to create the gene expression atlas is presented in Supplemental Table 6. For leaf sampling, the whole leaf was sampled before leaf emergence from the whorl. After the emergence, different leaf zones were sampled, as identified according to the physiological characters along the leaf. In brief, the division zone was sampled between 15-25 mm from the leaf base and contained mostly dividing cells. The elongation zone was sampled between 50-60 mm from the leaf base and contained mostly expanding cells. The differentiation zone was sampled between 100-140 mm from the leaf base and contained cells that stopped elongation but were not fully photosynthetically active. The mature zone was sampled as a piece of 40 mm in length at approximately the middle of the emerged leaf blade where cells were fully developed. During the vegetative stage (from V5 to 8 days after L18 appearance), L9, L12, L15 and L18 were sampled, while during the reproductive stage (from 4 days before ear appearance to silk emergence), the mature zone of L12, L18 and the ear leaf (usually L14 or L15) were sampled. Internodes were sampled in their entirety at the earliest growth stages. In the later stages, they were divided lengthwise in the middle, into a low (basal) and high (apical) zone or separated into three zones, with middle zone. Int9, Int12 and Int15 were sampled during the vegetative stage while the ear internode was sampled during the reproductive stage. Similar to the internodes, ears were sampled either in their entirety, divided lengthwise into two equally long parts or divided lengthwise into three equally long parts.

In total, 272 biological materials were collected for transcriptome analysis. For each of them, three biological replicates from independent greenhouse experiments were examined and each was a pool of three randomly chosen plants. Only for six materials, there were two biological replicates (Earx0_IntEarMid_V12, L18×0_IntEar_C, L18×0_IntEar_V5, L18×4_IntEarHigh_C, Silk_IntEarLow_V5, L9×0_L9Elong_V5).

### RNA extraction, sequencing, and analysis

Total RNA was extracted using TRIzol reagent (Thermo Fisher Scientific) according to the manufacturer’s instructions, followed by DNase treatment using the RQ1 RNase-free DNase kit (Promega). The extracted RNA was sent to GATC Biotech AG (Constance, Germany) for RNA sequencing. The libraries were constructed using NEBNext Kit for Illumina with poly-A selection. The mRNA-enriched libraries were sequenced in a paired-end mode with a read length of 125 bp on an Illumina HiSeq4000 with the TruSeq SBS Kit v3 (Illumina). The quality of the raw data was assessed using FastQC (http://www.bioinformatics.babraham.ac.uk/projects/fastqc/, version 0.11.5). Quality filtering was then performed with Trimmomatic (version 0.32) to remove adapters and low-quality reads. The filtered reads were subsequently mapped to the maize B73 reference genome sequence V3 (http://ftp.maizesequence.org/B73_RefGen_v3/) using GSNAP (v2013-06-27) (Wu et al., 2016). Concordantly paired reads that uniquely map to the genome were used for quantification on gene level with htseq-count from the HTSeq.py python package (version 0.6.1p1) (Anders et al., 2015). All analyses were done on the Galaxy instance of the VIB-UGent Center for Plant Systems Biology (Afgan et al., 2016). Raw count data (39,323 genes, 810 samples) were filtered to remove 3,291 genes with zero counts across all samples and 7,690 further genes with >0.5 counts per million (cpm) in <3 samples, resulting in a raw count matrix encompassing 28,342 genes across 810 samples, which was used for all further analyses. Differential expression analysis was performed with the edgeR version 3.27.13 (Robinson et al., 2010) and limma version 3.41.16 (Ritchie et al., 2015) packages in R version 3.6.0 (R Core Team, 2013) as described in Law et al. (2016), using the voom method (Law et al., 2014) to model the mean-variance relationship in the data. DEGs were defined by an absolute log_2_ fold change >1 and a Benjamini–Hochberg FDR-adjusted *P*-value <0.05 (Benjamini and Hochberg, 1995). Data normalization for visualization, principal component analysis (PCA) and clustering was done using DESeq2 version 1.25.10 (Love et al., 2014). The filtered raw count matrix was normalized and transformed using the vst() function of DESeq2, which performs DESeq2 size factor calculation, normalization for library size and fast estimation of the mean-dispersion trend in the data followed by a variance-stabilizing transformation (vst) (Love et al., 2014). vst() was run with default parameters. PCA was performed on the centered vst counts using the prcomp() function in the R stats package. The genes contributing most to the PCA separation were identified as the genes with the top 100 positive/negative loadings for PC1 and PC2. For identification of organ-specific genes, the tau metric was used as a measure to determine the gene specificity, with tau=0 meaning broad gene expression and tau=1 specific gene expression (Kryuchkova-Mostacci and Robinson-Rechavi, 2017). tau >0.9 was used as an empirical cutoff in classifying specifically expressed genes. GO enrichment analysis was performed using BINGO (Maere et al., 2005) and Benjamini-Hochberg adjustment of p-values to control the FDR (Benjamini and Hochberg, 1995). GO terms with an FDR-adjusted p-value <0.05 were considered significantly enriched. *K*-means clustering was performed on scaled & centered vst mean expression of genes that were differentially expressed in at least three comparisons among zones or time points from control samples (24,880 genes), leaf samples (22,480 genes), and internode samples (19,616 genes) using the *k*-means algorithm in R version 3.6.0 (R Core Team, 2013), defining 18, 15, and 9 clusters, respectively. Complete-linkage hierarchical clustering with Euclidean distance was conducted on the vst-normalized expression values using the Shiny app available at https://www.psb.ugent.be/shiny/grand-transcriptome/.

### Accession numbers

Unique identifiers for all genes mentioned in the text are provided in Supplemental Data Set 18. All data supporting this study are included in the article and its Supplementary material. Sequence data from this article can be found in ArrayExpress under accession number E-MTAB-14898.

## Supporting information

Supplemental Data Set 1

Supplemental Data Set 2

Supplemental Data Set 3

Supplemental Data Set 4

Supplemental Data Set 5

Supplemental Data Set 6

Supplemental Data Set 7

Supplemental Data Set 8

Supplemental Data Set 9

Supplemental Data Set 10

Supplemental Data Set 11

Supplemental Data Set 12

Supplemental Data Set 13

Supplemental Data Set 14

Supplemental Data Set 15

Supplemental Data Set 16

Supplemental Data Set 17

Supplemental Data Set 18

## Author contributions

N.W., S.C.B., J.V., W.B, D.I., S.Ma., and H.N. conceived the original screening and research plans; D.I., S.Ma., and H.N. supervised the project. L.V., N.W., and S.Me. performed the Phenovision experiments, sample preparation and RNA extraction; L.V. analyzed the organ growth data; J.Z and H.S performed transcriptomic data analysis and interpretation; B.C., J.D.B., K.D., A.N., K.M., J.M., provided technical assistance during experiments; J.Z., L.V., S.Ma., and H.N. wrote the article with input from the other authors.

## Supplemental Figures

**Supplemental Figure 1.**
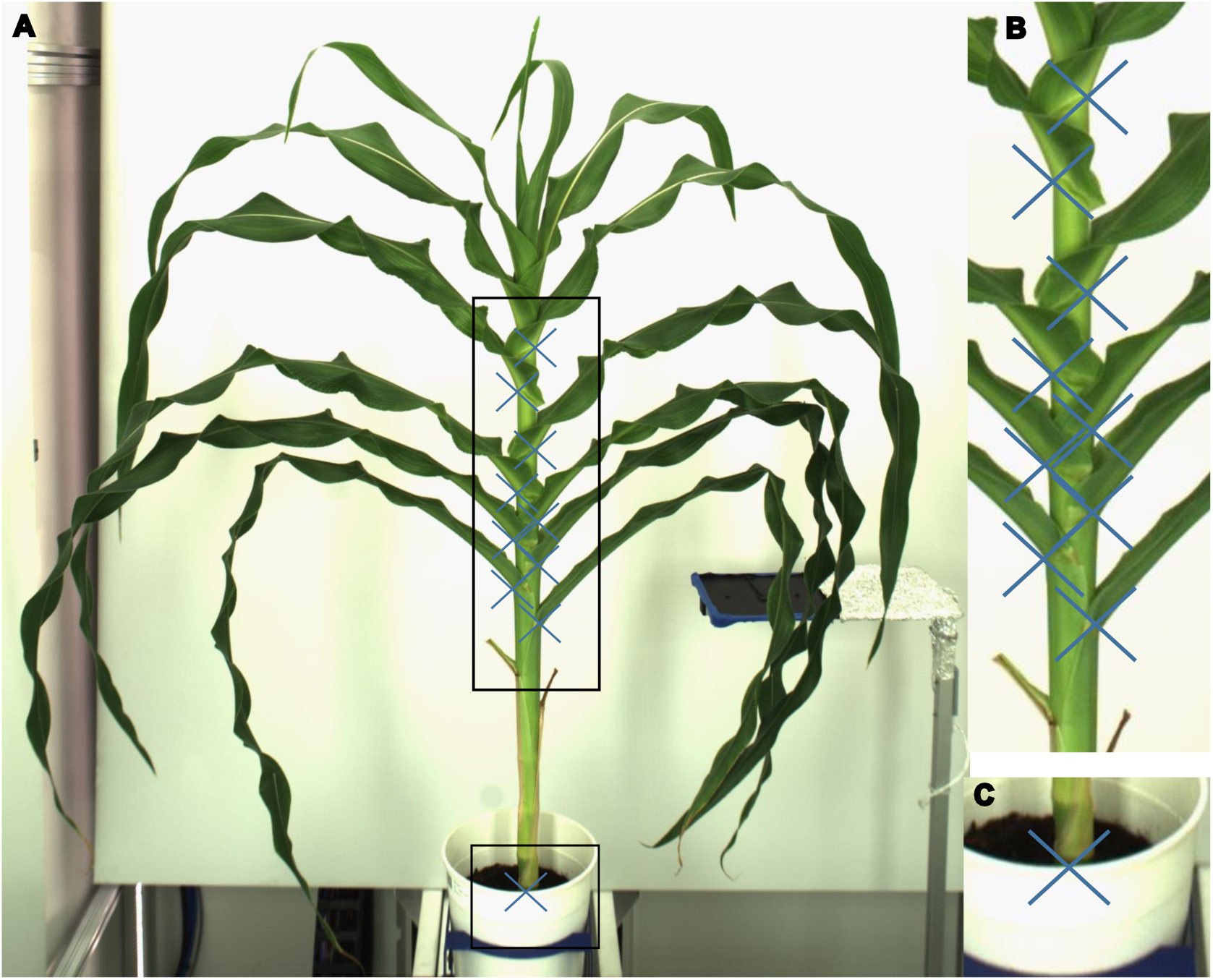
Leaf collar positions and soil level shown in a plant image captured in the Phenovision platform. **A)** Illustration of a plant image with an X marking the position of every leaf collar and the soil level. Leaf collar height was determined as the distance between the position of the leaf collar and the soil level. Areas marked with rectangles are shown in detail in B and C. **B)** Closeup detail of stem and leaf sheaths, showing exactly where a leaf collar the position was determined. **C)** Closeup detail of the stem at soil level.

**Supplemental Figure 2.**
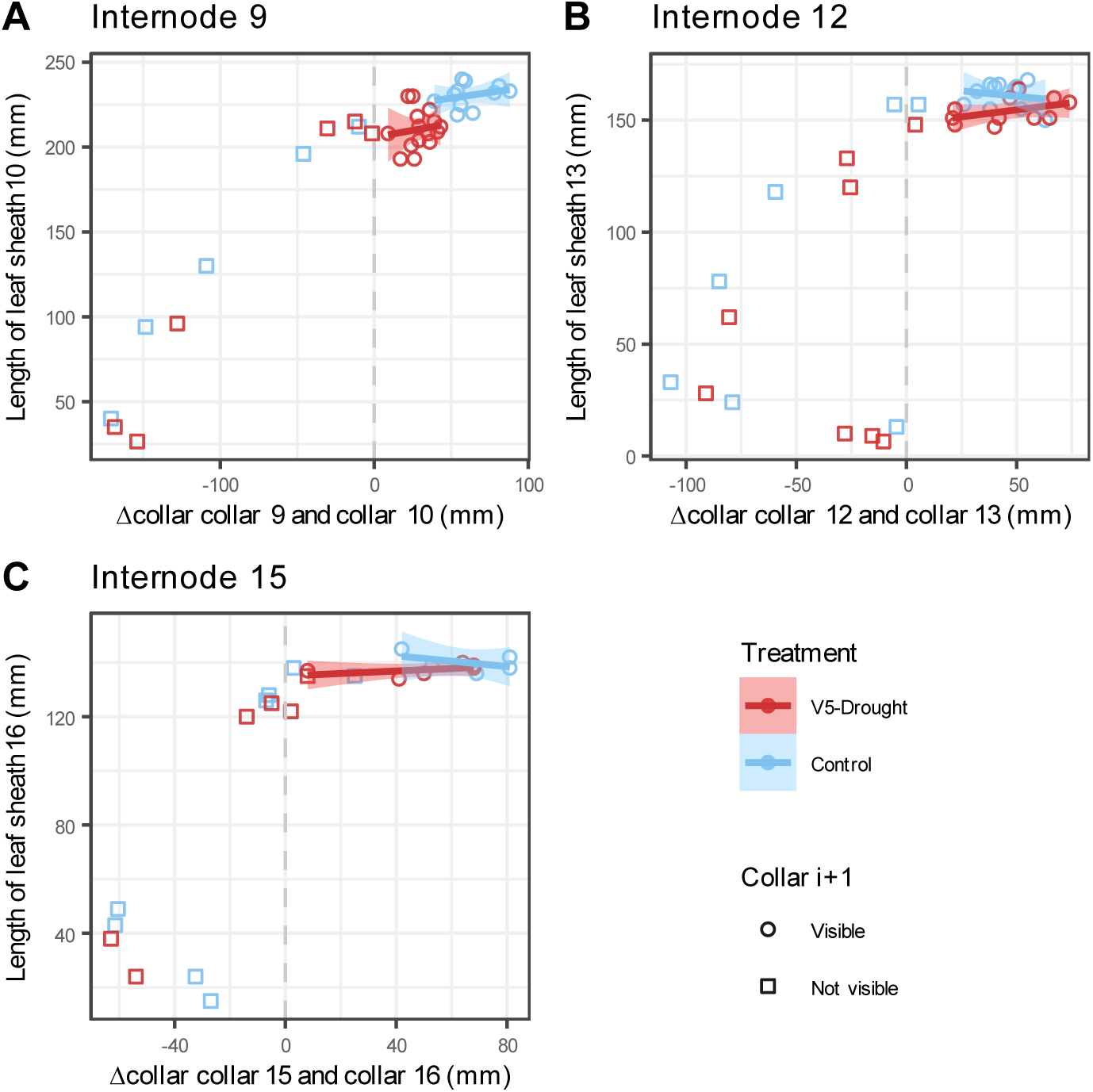
Illustration of the relation between leaf sheath length and Δcollar. The relation between length of leaf sheath i+1 and the difference in collar height between leaves i and i+1 is shown for i=9 (**A**), i=12 (**B**) and i=15 (**C**) in both Control (blue) and V5-Drought (red) conditions. Circles indicate data points after the emergence of the higher leaf collar i+1, where leaf collar i+1 is visible on the image. Squares indicate data points where leaf collar i+1 was not visible on the image, before the emergence of leaf collar i+1. Straight lines (linear models) were fitted to the data points with a visible higher leaf collar i+1 to examine the relationship between the length of the higher leaf sheath i+1 and the difference in collar height (Δcollar). There is no relationship between the length of the higher leaf sheath i+1 and the difference in collar height when the higher leaf collar i+1 is visible, indicating leaf sheath elongation is completed before the leaf collar emerges above the lower leaf collar and becomes visible.

**Supplemental Figure 3.**
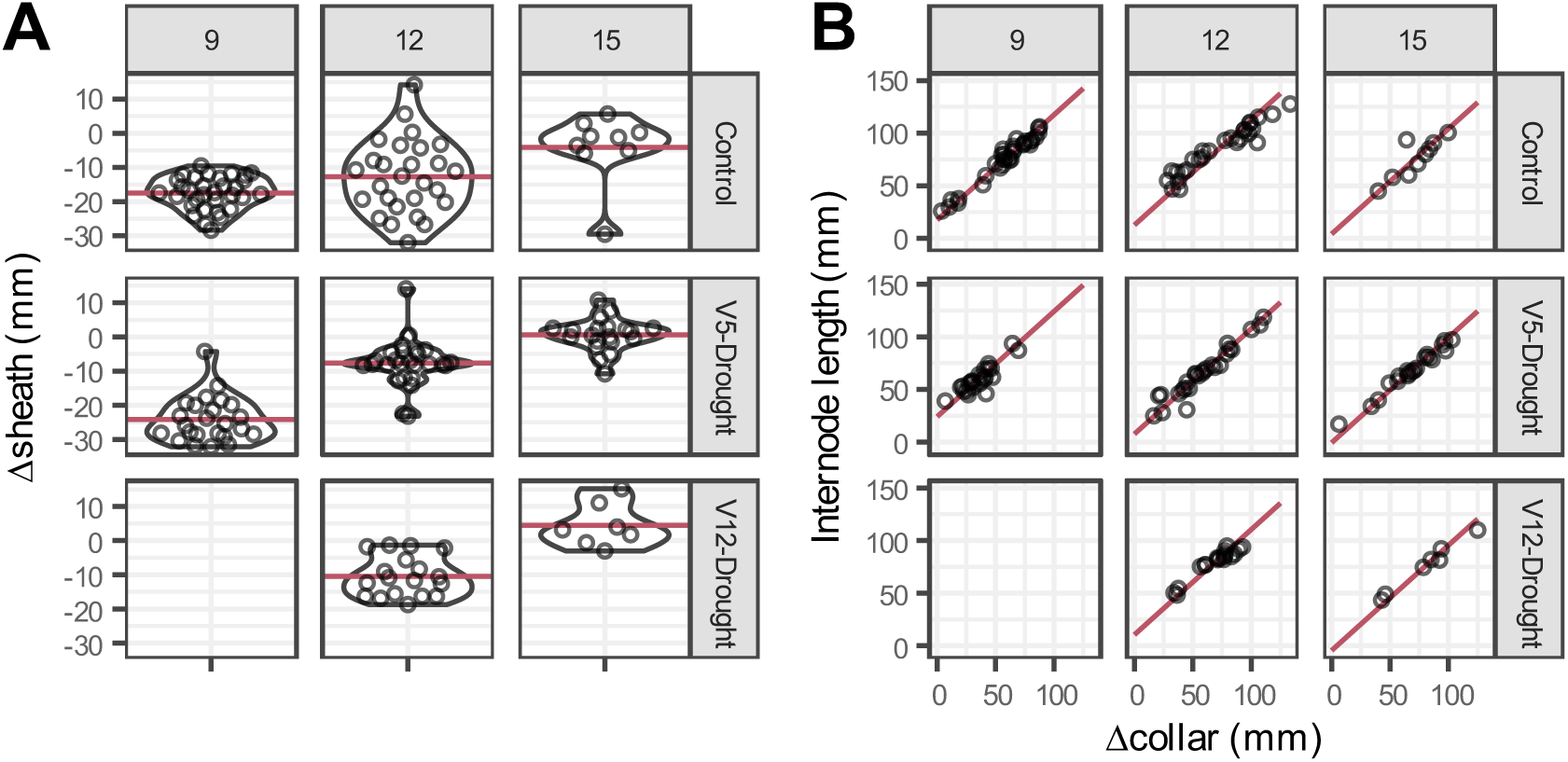
Distribution of Δsheath and relation between internode length and Δcollar. **A)** Violin plots of Δsheath values for the different internode ranks (9, 12 and 15) and watering treatments (Control, V5-Drought and V12-Drought) in the ground truth dataset. Red horizontal lines indicate mean Δsheath values used for the internode length estimation. **B)** Relation between internode length and Δcollar for the different internode ranks (9, 12 and 15) and watering treatments (Control, V5-Drought and V12-Drought). Red diagonal lines indicate the estimated internode lengths using the mean Δsheath values.

**Supplemental Figure 4.**
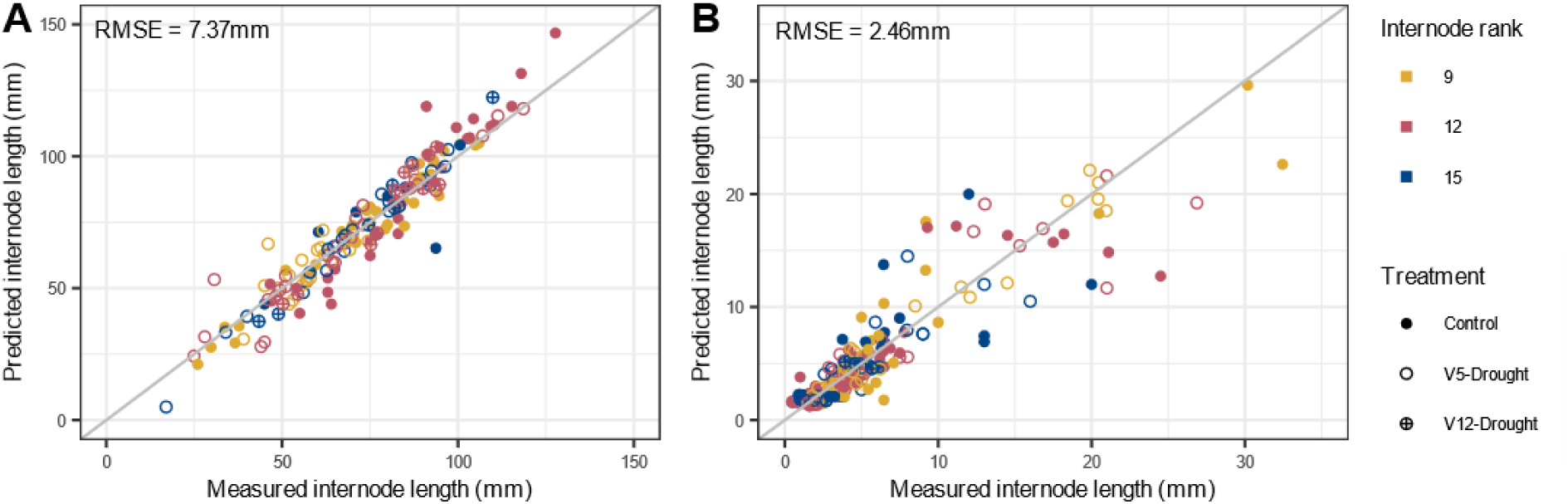
Performance of image-based internode length estimation methods for three different internodes under control and drought treatments. **A)** Direct internode length estimation, deriving the length of internode i from the difference in height between leaf collars i and i+1 and the mean Δsheath values for that internode rank and watering treatment. Predictions were made using leave-one-out-cross-validation: each estimate was determined using the mean difference in sheath length determined on the other ground truth measurements, not including the one being estimated. **B)** Indirect internode length estimation, for cases where the leaf collar at position i+1 was not visible on the image and the direct length estimation method could not be used. The length of internode i was predicted using a linear model on the difference in collar height between the two highest visible leaf collars at positions i and i-1, i-1 and i-2 or i-2 and i-3. Predictions were again made using leave-one-out-cross-validation.

**Supplemental Figure 5.**
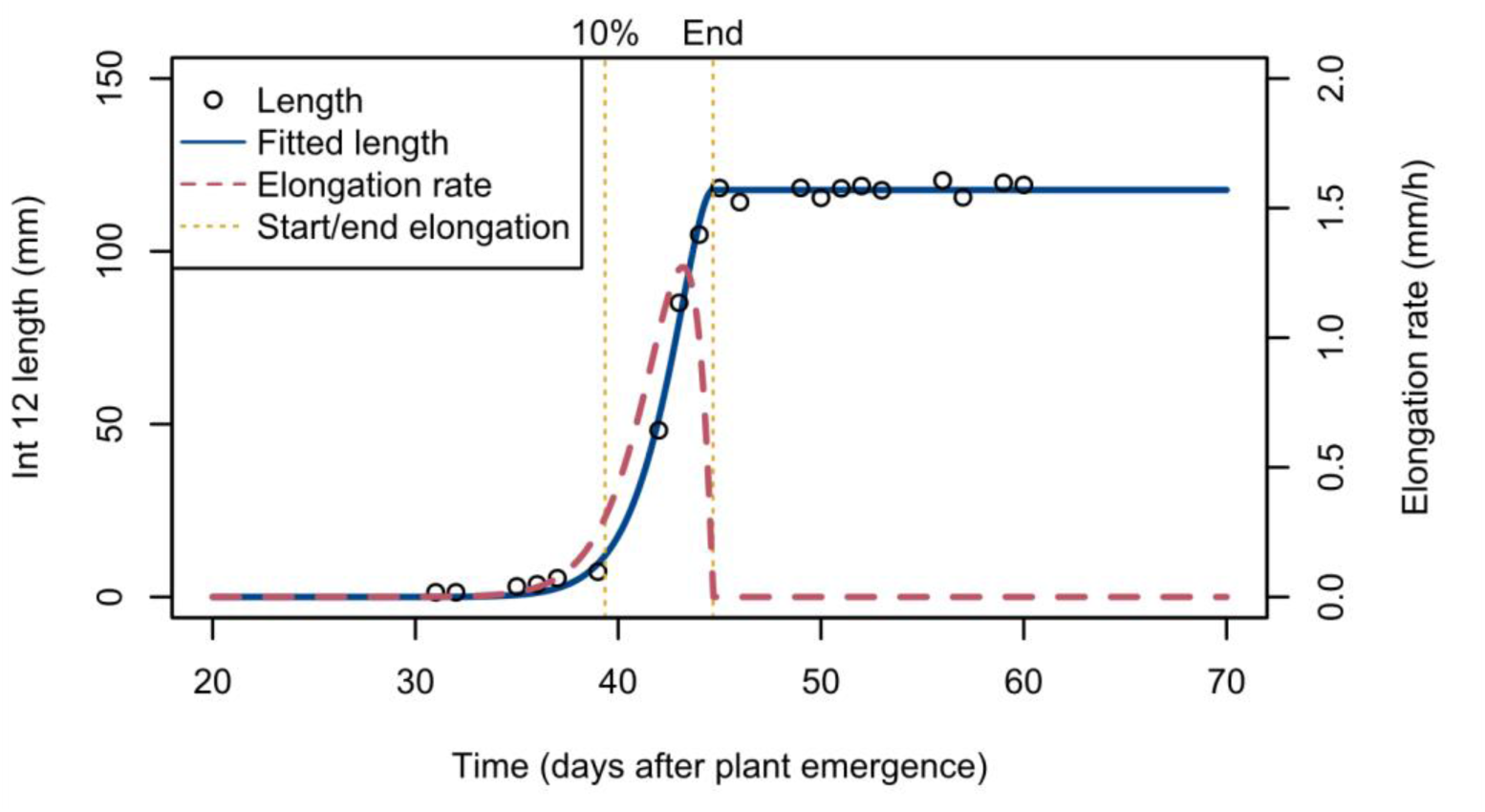
Estimates of internode 12 length and elongation rate over time. Circles (left y-axis) represent the internode 12 length estimates for a control-treated plant, with a beta-sigmoid growth function (blue solid line, left y-axis) fitted to the data. The red dashed line (right y-axis) shows the derived elongation rate. Yellow vertical dotted lines indicate the time when length reaches 10% of its maximum and reaches its maximum. These time points are considered as start and end of elongation for determining elongation duration and mean elongation rate.

**Supplemental Figure 6.**
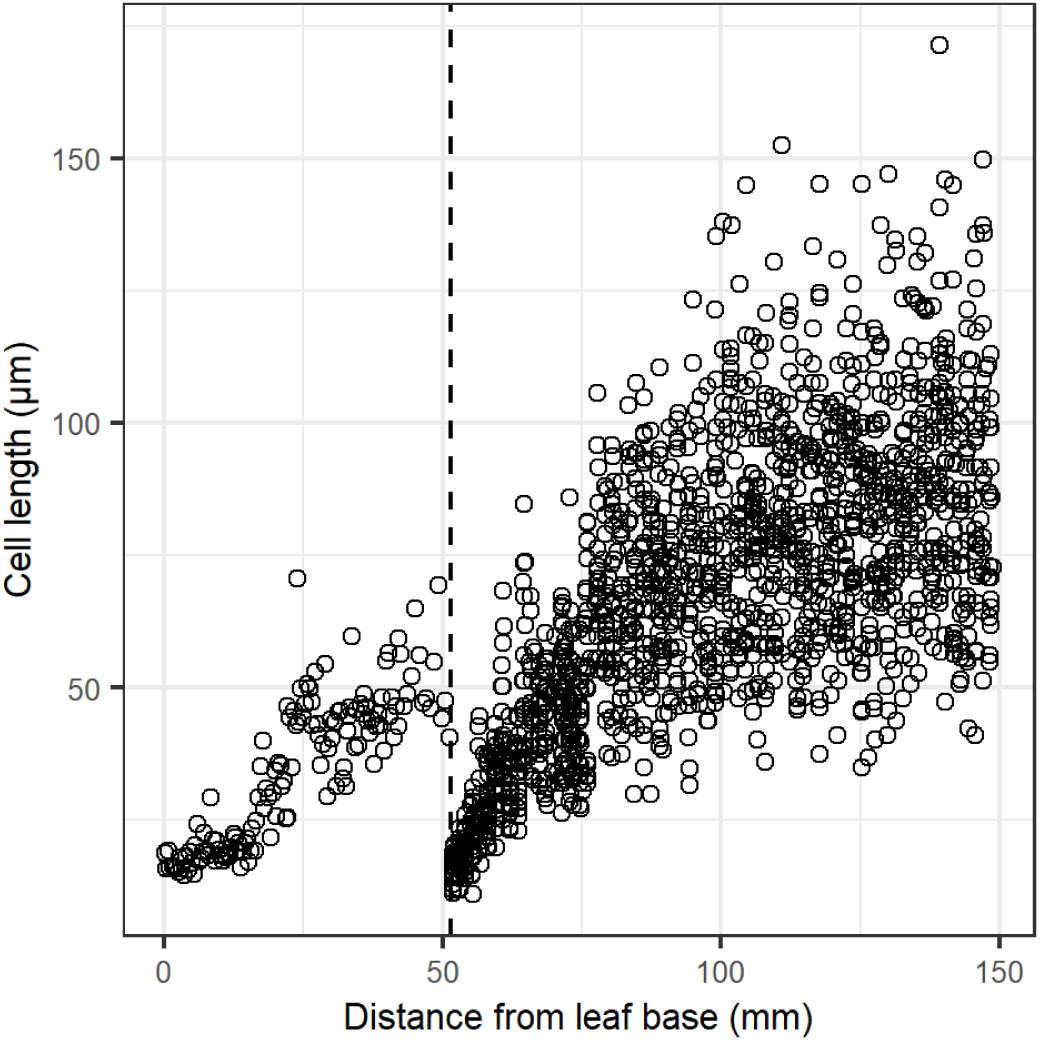
Cell length profile of leaf 9 at 8 days after leaf emergence under well-watered conditions. Points are individual cells, measured at random. The distinct growth zones for the leaf sheath (at the leaf base) and leaf blade (starting from 51.5 mm) are clearly visible in this case. The vertical dashed line at 51.5 mm indicates the ligule position.

**Supplemental Figure 7.**
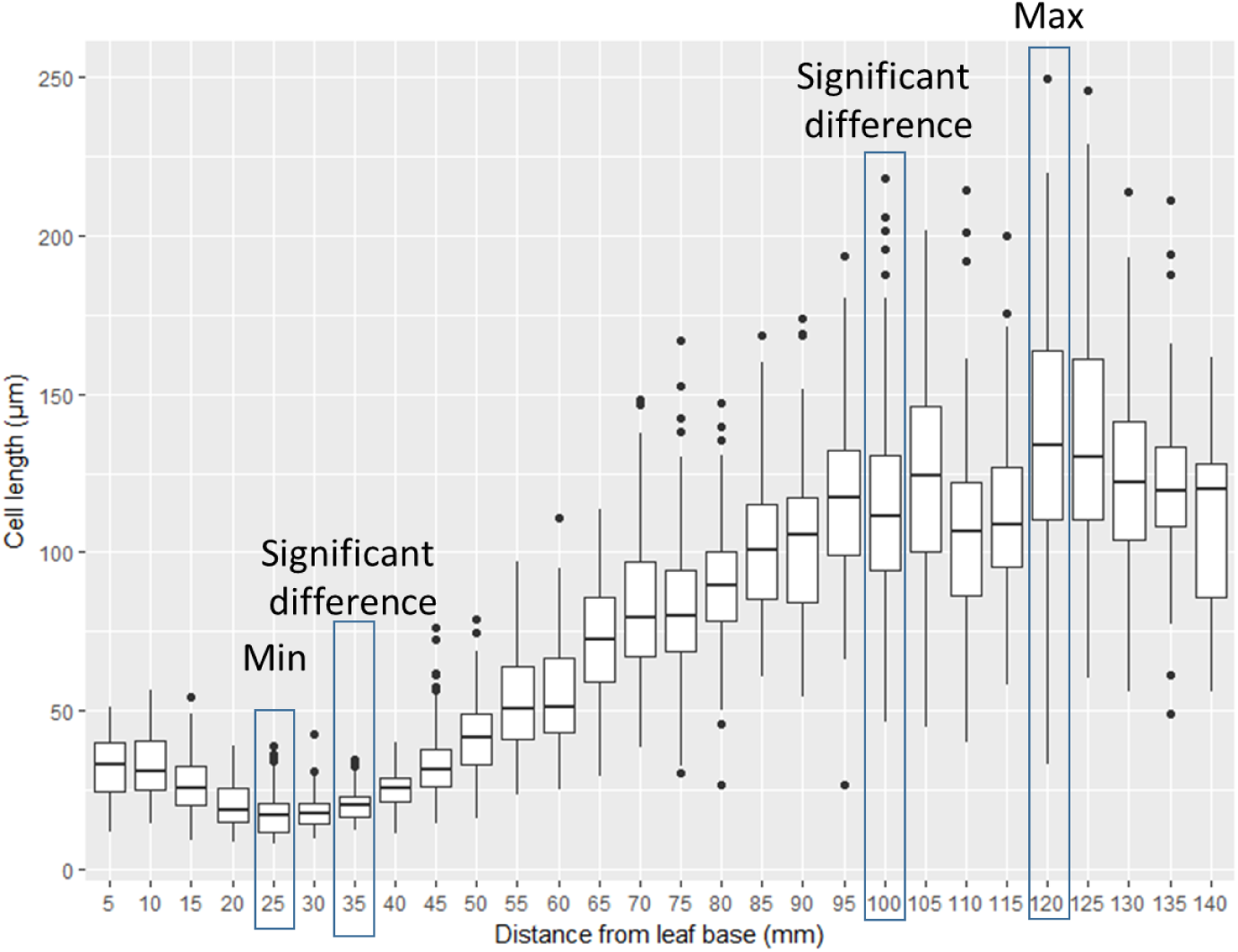
Epidermal cell length profile of leaf 12 under well-watered conditions. The epidermal cell length is shown over distance from leaf base for leaf 12 at leaf appearance, with cell lengths grouped in bins per 5mm. The bin at 25 mm has the lowest mean cell length. Cell length in bin 30 is not significantly different from the cell length in bin 25, but for bins 35, 40, 45, 50 and 55 there is a significant difference (t-test, p<0.05). The first of these five bins with a significant difference indicates the end of the division zone. The bin at 120 mm has the highest mean cell length. Cell length is significantly different in bins 115 and 110, but not in bin 105. In bins 100, 95, 90, 85 and 80 there is again a significant difference to bin 120 (t-test), which means that bin 100 indicates the end of the elongation zone.

**Supplemental Figure 8.**
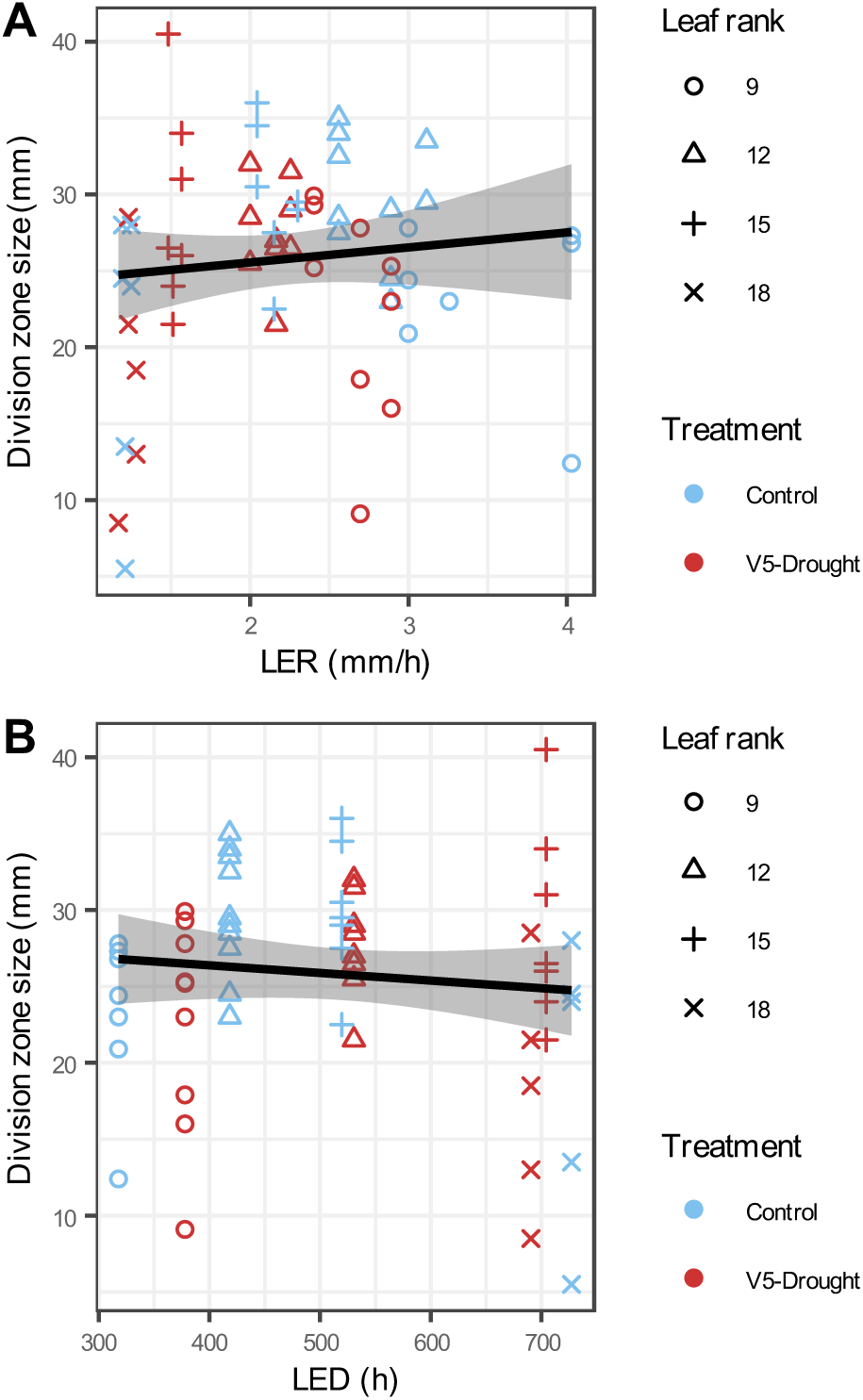
Relation between leaf division zone size and estimated leaf elongation rate (LER) as well as leaf elongation duration (LED). **A)** Relation between leaf division zone size and estimated LER at the time of sampling. **B)** Relation between leaf division zone size and LED. Each growth zone sample, for the two treatments (Control and V5-Drought), four leaf ranks (9, 12, 15 and 18), and the three sampling time points (day of leaf emergence, 4 days after, 8 days after), was matched to the LER and LED derived for the corresponding leaf rank, treatment, and sampling time point based on the fitted growth curves in Verbraeken et al. (2021). Black lines are linear regressions fitted to the data, whereas shaded areas indicate the 95% confidence interval for the fitted lines.

**Supplemental Figure 9.**
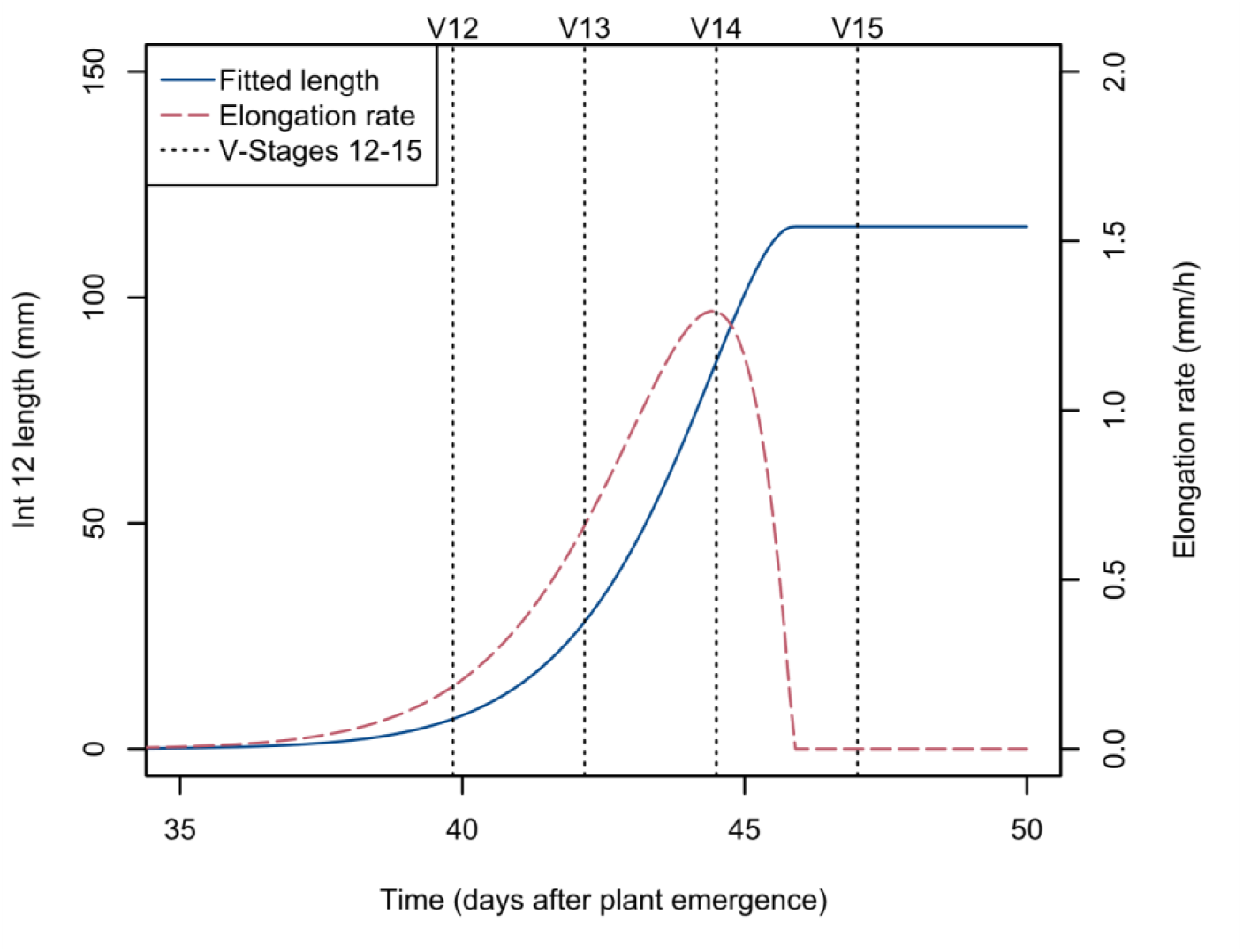
Timing of V-stages relative to internode 12 growth. The timing of V-stages (vertical black dotted lines) is shown relative to fitted internode 12 length (blue solid line, left y-axis) and elongation rate (red dashed line, right y-axis) for well-watered plants.

**Supplemental Figure 10.**
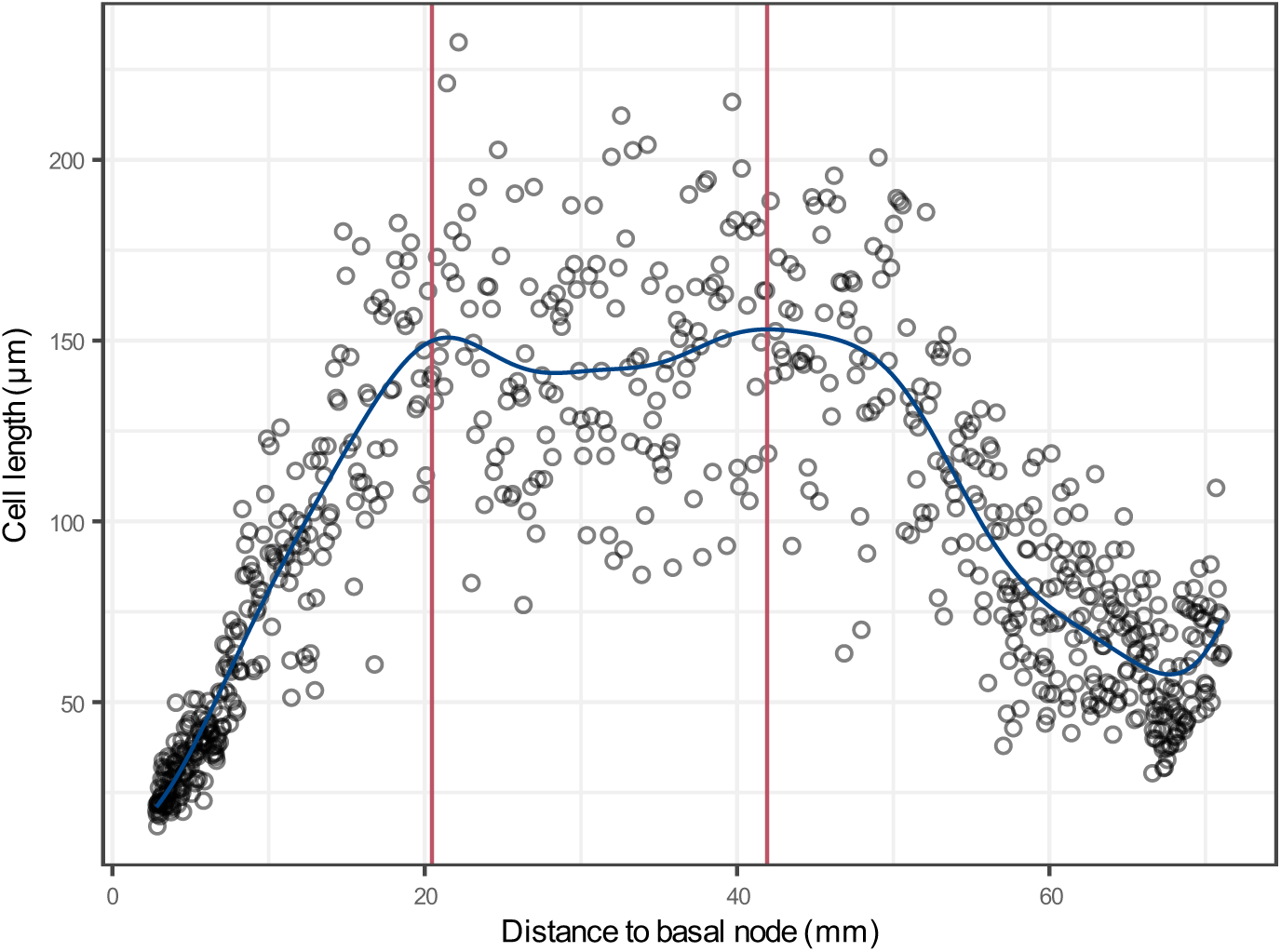
Cell length profile for a single internode 12. The cell length profile of one particular internode is shown, illustrating the reason behind imposing a limit on the end of the elongation zone as the maximal fitted cell length within the basal 40 mm. Black points indicate the lengths of individual cells along the internode length, and the solid blue line indicates the fitted cell length. The red vertical line on the left indicates the considered end of the elongation zone at 20.5 mm, whereas the red vertical line to the right indicates where the end of the elongation zone would have been set (41.9 mm) without the imposed limitation to locate it in the basal 40 mm. The cells between 20.5 mm and 41.9 mm are considered not to be elongating.

**Supplemental Figure 11.**
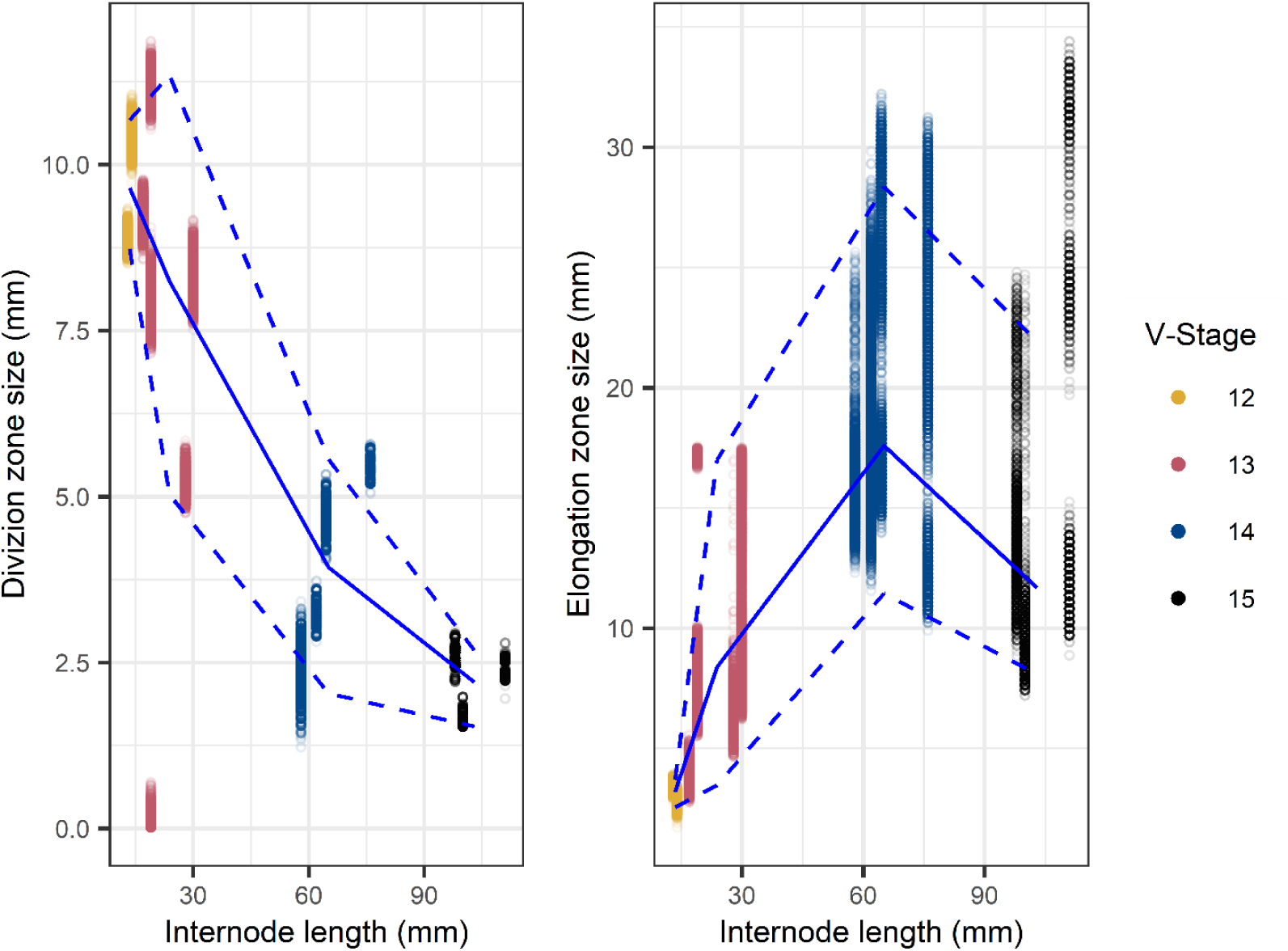
Relation between internode division and elongation zone size and total internode length for internode 12. **A)** Relation between the size of the division zone and the total internode length. **B)** Relation between the size of the elongation zone and the total internode length. Individual points are the results of the determination of zone size on bootstrap resampled distributions of cell lengths, the solid line connects the mean values per V-stage, and the dashed lines connect the upper and lower limits of the 95% confidence interval per V-stage.

**Supplemental Figure 12.**
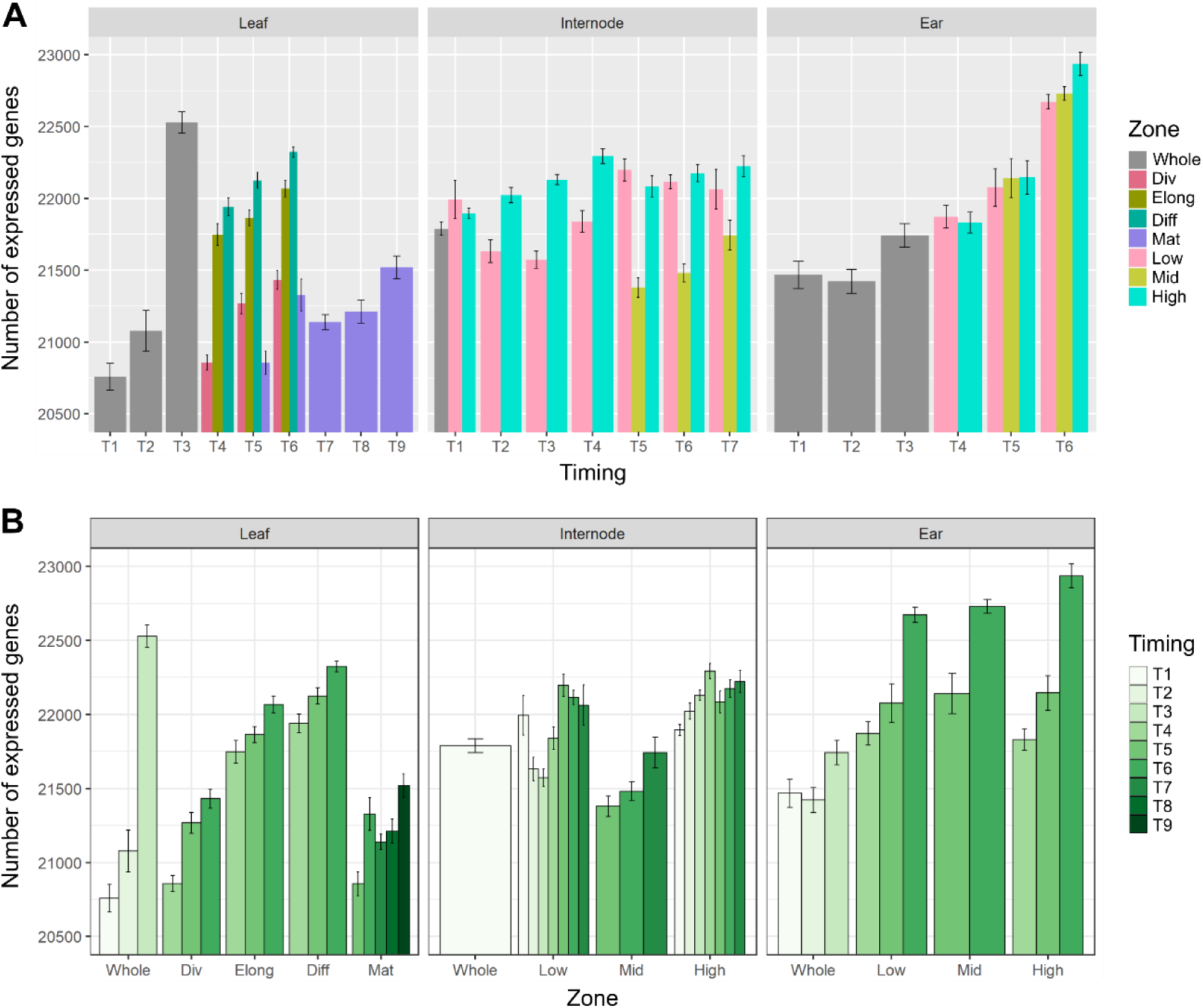
Number of expressed genes in all tissues. **A)** Total number of expressed genes in different zones of leaf, internode and ear tissue at different temporal stages (x-axis). **B)** Total number of expressed genes in different temporal stages of leaf, internode and ear tissue in different zones (x-axis). Only genes with more than 0.5 counts per million (cpm) in at least three samples were considered as expressed.

**Supplemental Figure 13.**
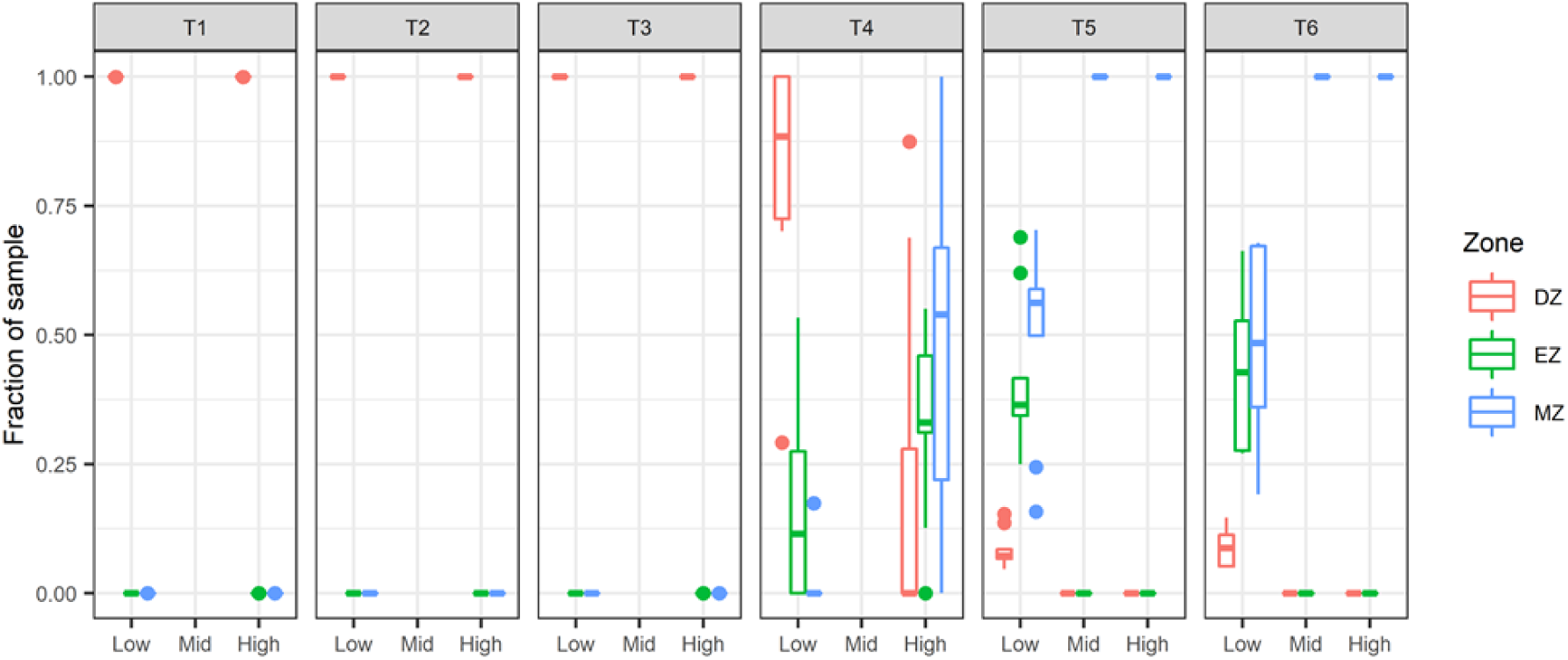
Fraction of the internode 12 well-watered transcriptome samples attributed to growth zones at different temporal stages. From T1 to T3, the entire internode 12 is considered division zone, at T4 88% of the low zone is division zone and 12% is elongation zone, while 0% of the high zone is division zone, 33% is elongation zone and 66% is mature zone. At T5, 7% of the low zone is division zone, 36% is elongation zone and 56% is mature zone. The mid and high zone are 100% mature zone. At T6, 9% of the low zone is division zone, 43% is elongation zone and 48% is mature zone. The mid and high zone are 100% mature zone.

**Supplemental Figure 14.**
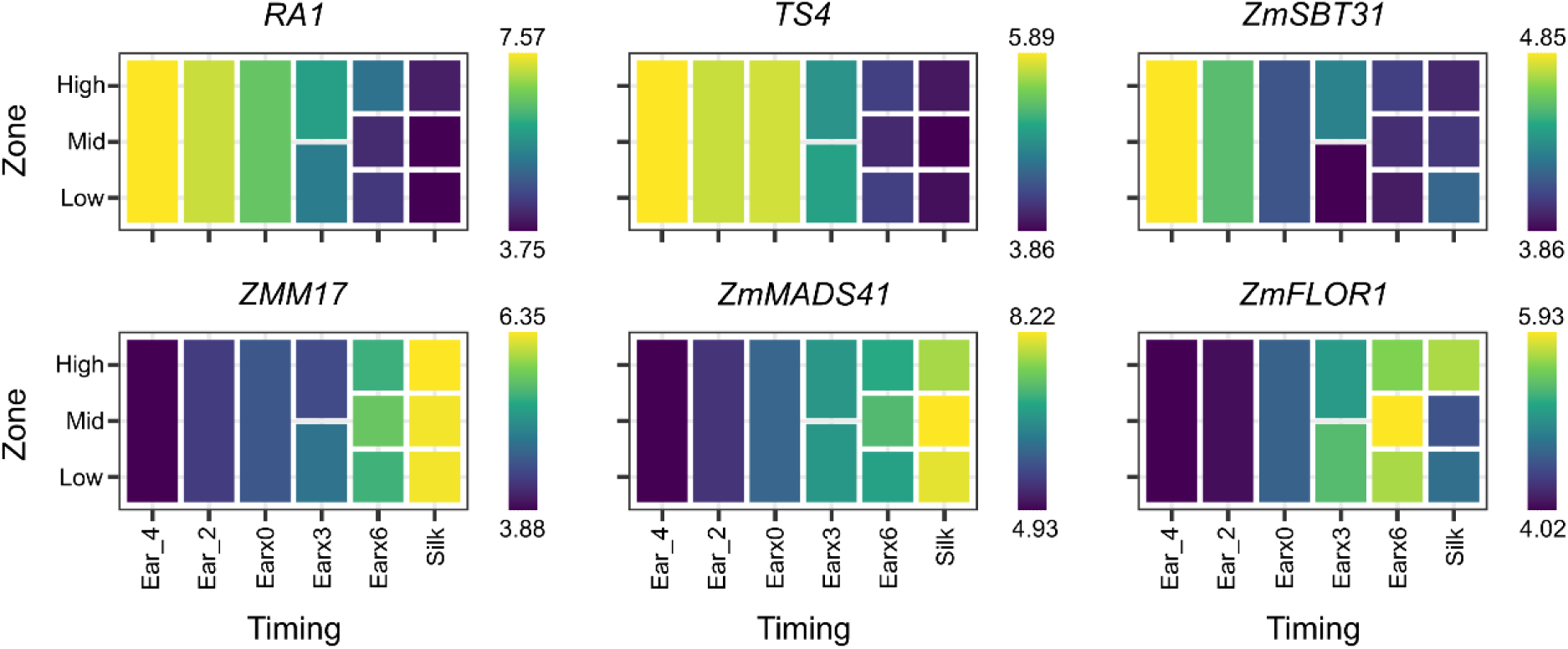
Heatmaps of six spatiotemporal DEGs during ear development. Gene expression values are variance-stabilized transformed counts.

**Supplemental Figure 15.**
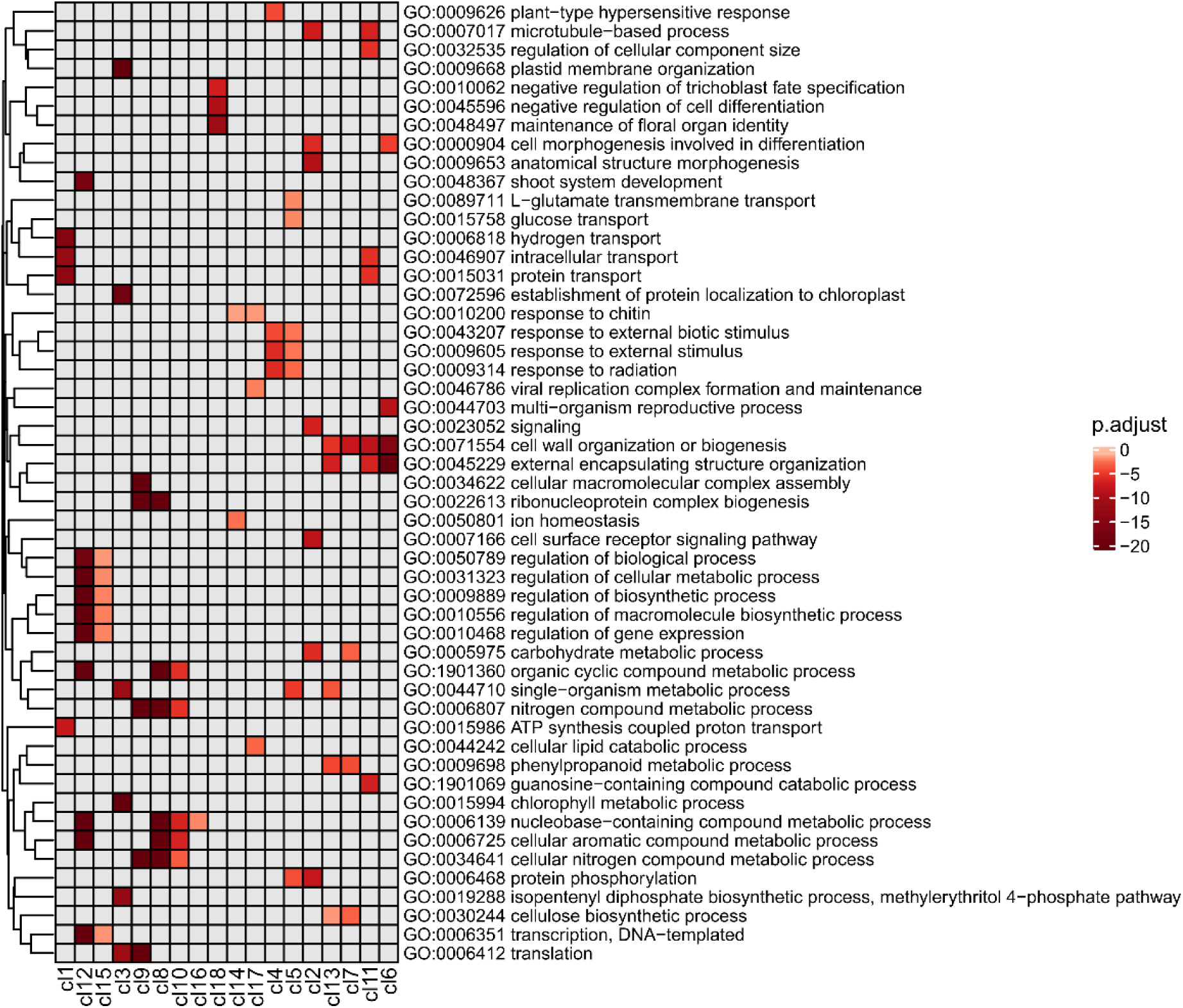
Clustered heatmap of GO-term enrichment analysis for genes in 18 clusters obtained by *k*-means clustering on expression profiles of control samples. Rows are clustered based on semantic similarity and the list of GO terms is filtered to reduce the complexity. The color intensities indicate the level of enrichment score (Benjamini-Hochberg FDR-adjusted p-values) of each GO term.

**Supplemental Figure 16.**
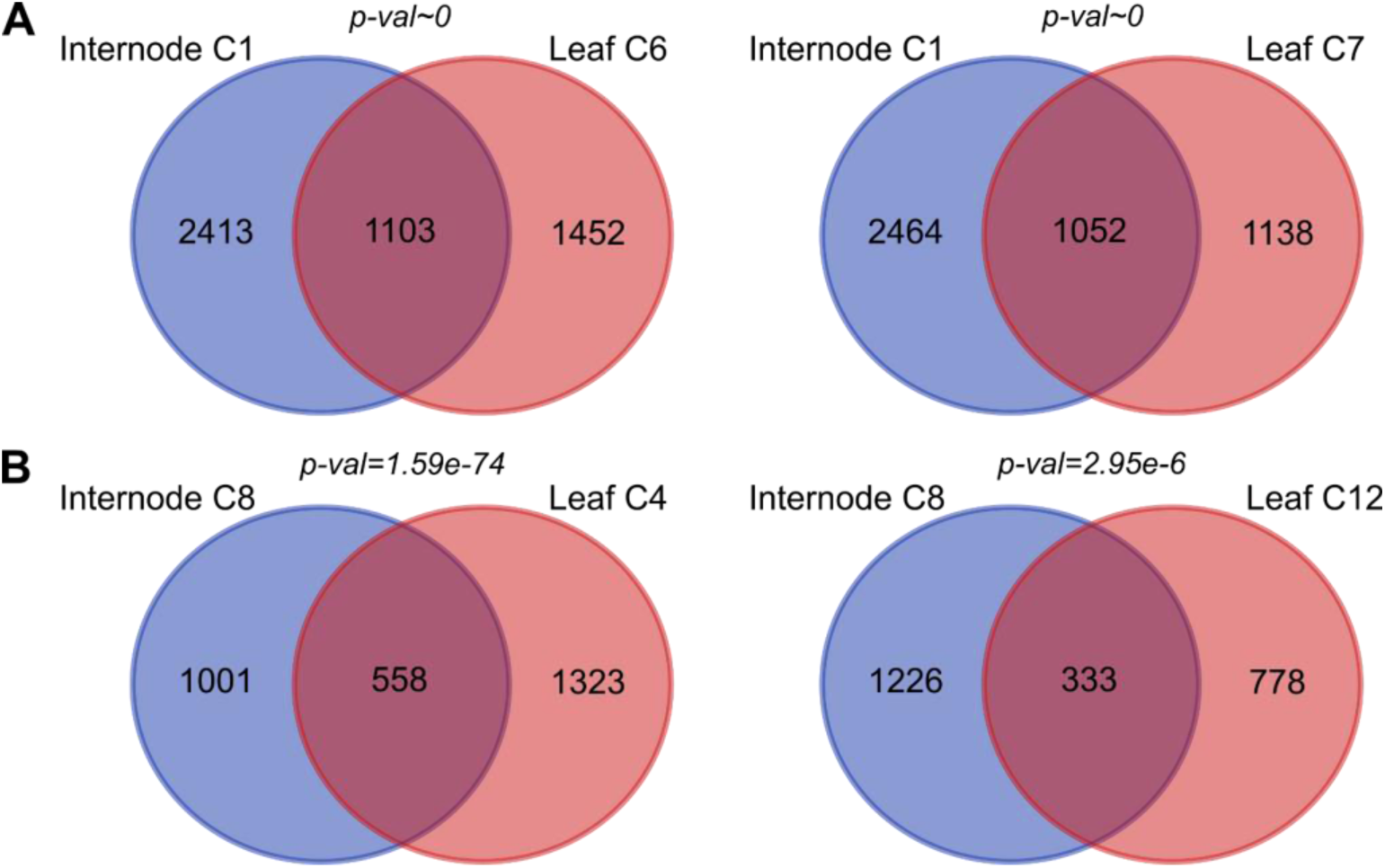
Overlap of DEGs between internode clusters and leaf clusters. **A)** Venn diagram showing the overlapping DEGs between internode cluster 1 (C1) and leaf clusters 6 (C6) and 7 (C7). **B)** Venn diagram showing the overlapping DEGs between internode cluster 8 (C8) and leaf clusters 4 (C4) and 12 (C12). The *p*-values are the result of a hypergeometric test.

**Supplemental Figure 17.**
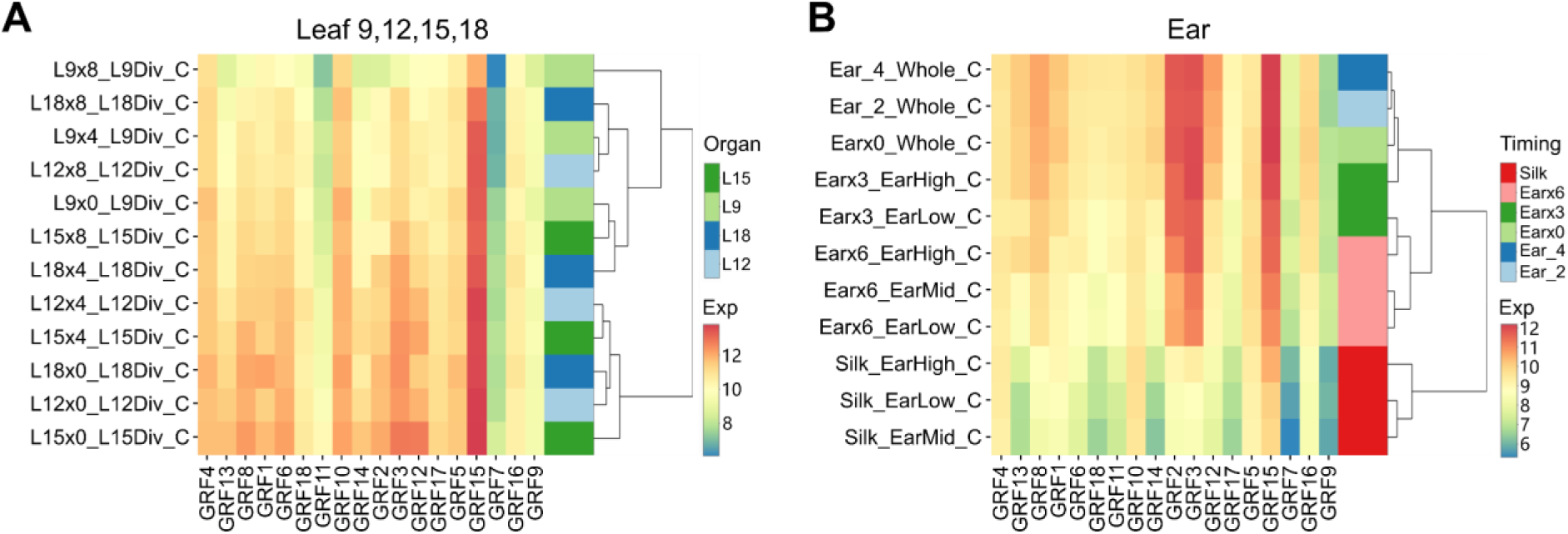
Expression patterns of *GRF* family genes in leaves, internodes, and ear. **A)** Hierarchical clustering heatmaps of 18 *GRF* genes in the leaf division zone of L9, L12, L15, and L18 over time. **B)** Hierarchical clustering heatmaps of 18 *GRF* genes across different zones of ear tissue over time. Gene expression values (Exp) are variance-stabilized transformed (vst) counts.

**Supplemental Figure 18.**
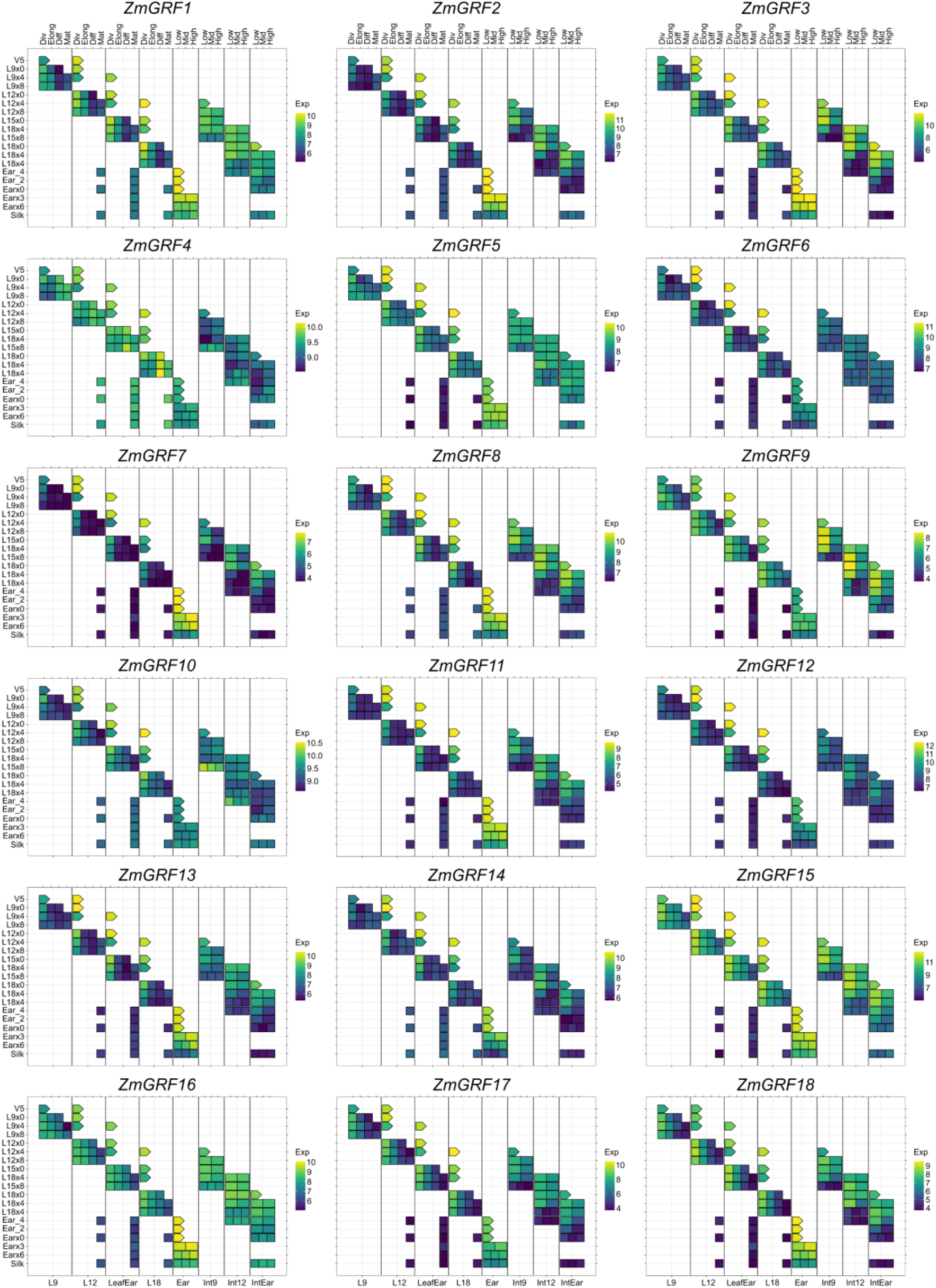
Expression heatmaps of individual *GRF* genes during development for leaves and internodes of different ranks and ear tissue. Gene expression values (Exp) are variance-stabilized transformed (vst) counts.

**Supplemental Figure 19.**
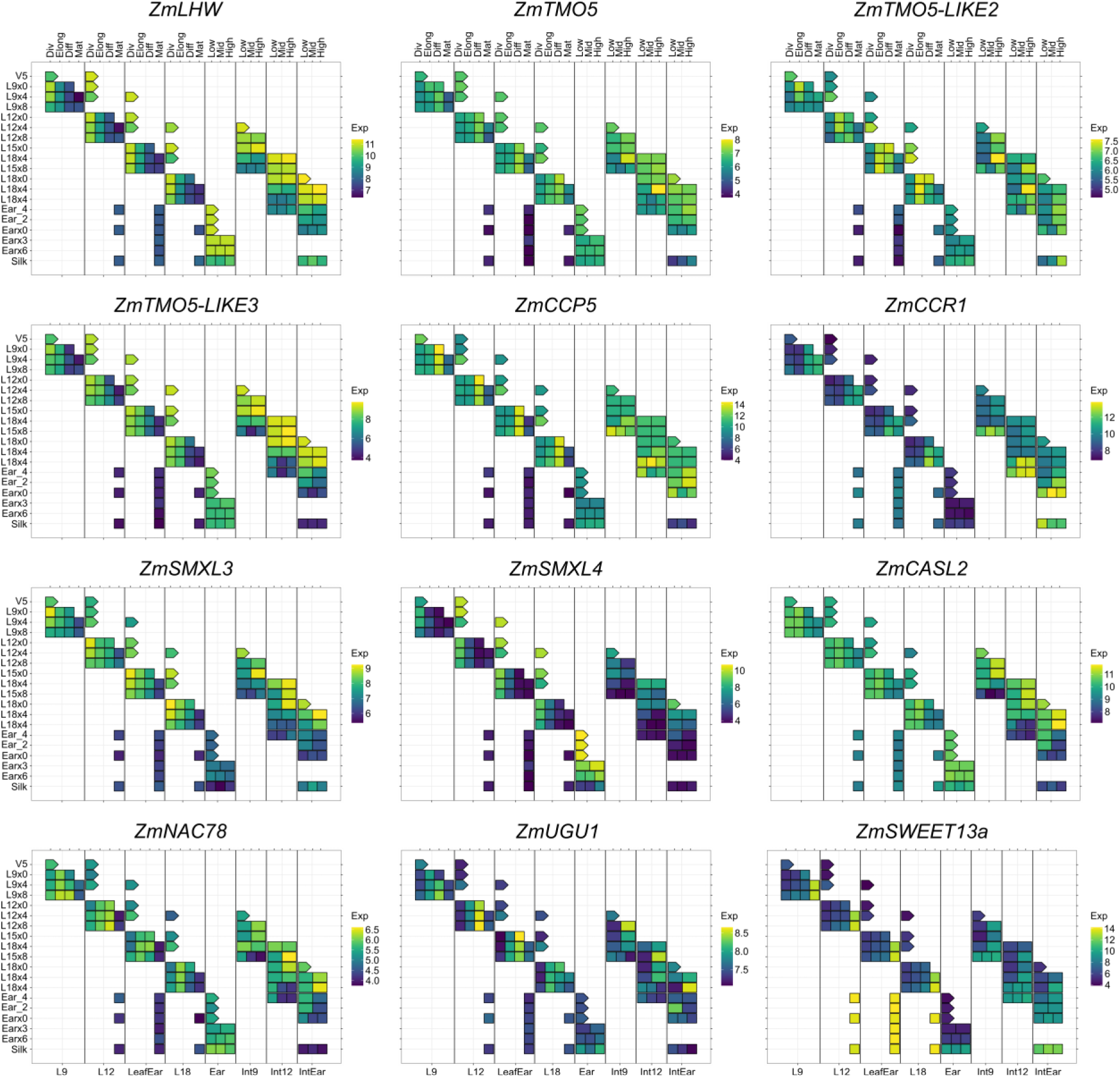
Expression heatmaps of vascular-related genes during development for leaves and internodes of different ranks and ear tissue. Gene expression values (Exp) are variance-stabilized transformed (vst) counts.

**Supplemental Figure 20.**
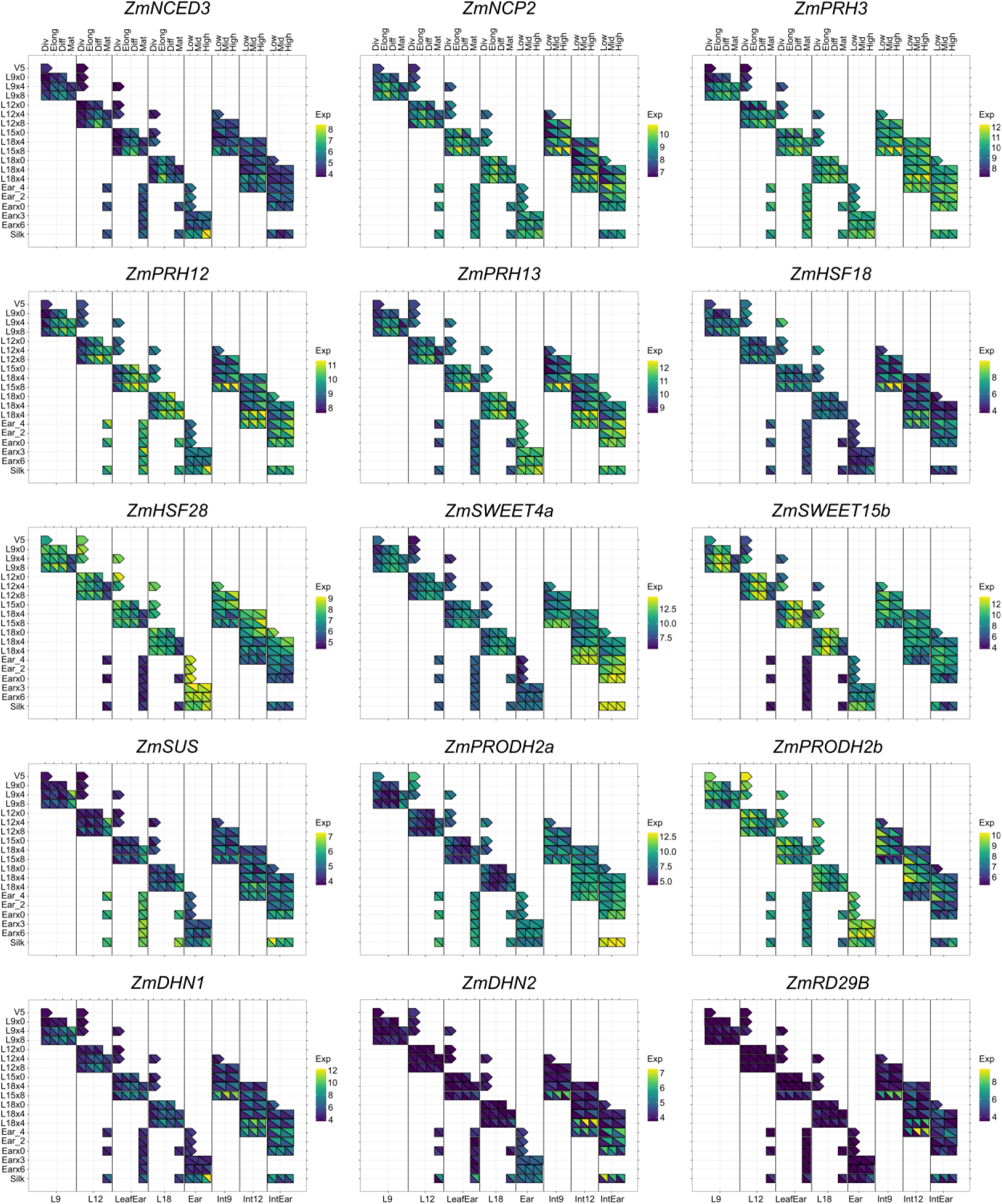
Expression heatmaps of several DEGs upon drought treatment during development for leaves and internodes of different ranks and ear tissue. The right upper triangle of each square represents gene expression under V5-Drought conditions, while the left lower triangle represents gene expression under Control conditions. Gene expression values (Exp) are variance-stabilized transformed (vst) counts.

**Supplemental Figure 21.**
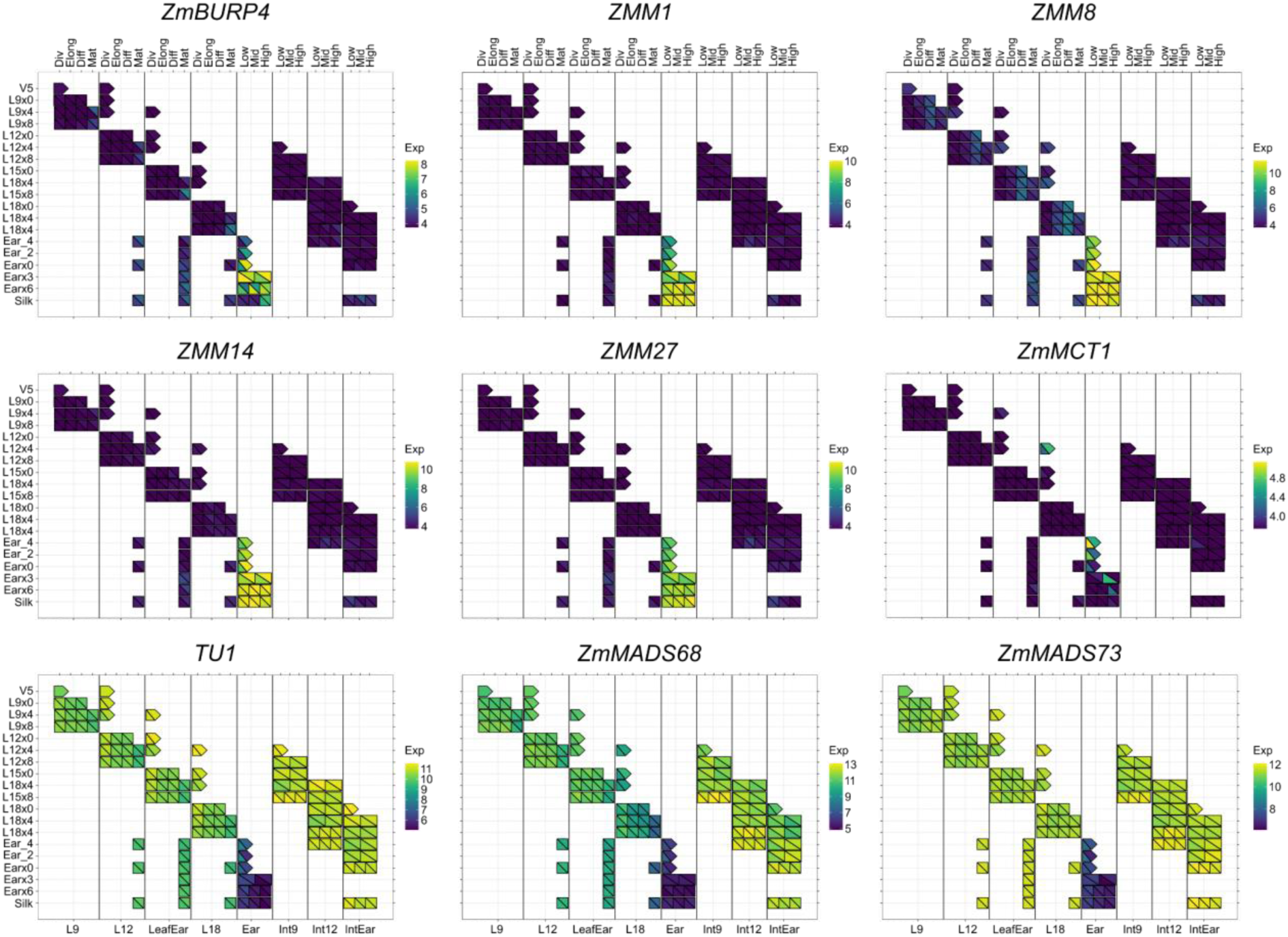
Expression heatmaps of several DEGs upon drought treatment during development for leaves and internode of different ranks and ear tissue. The right upper triangle of each square represents gene expression under V5-Drought conditions, while the left lower triangle represents gene expression under Control conditions. Gene expression values (Exp) are variance-stabilized transformed (vst) counts.

## Supplemental Tables

**Supplemental Table 1.** Sampling scheme of the destructive internode sampling. C, V5 and V12 indicate if a sample was taken for the Control, V5-Drought or V12-Drought treatment, respectively. Each sample was taken in three replicates in each experiment, with nine replicates in total. Starting from 4 days before ear emergence, the ear internode was sampled instead of internode 15. If the ear internode was internode 15, it was included in further analyses as internode 15.

| Timing plant development<br>(+/- x days) | Int9 | Int12 | Int15 | IntEar |
| --- | --- | --- | --- | --- |
| Leaf 12 appearance+4 (L12x4) | C, V5 |  |  |  |
| Leaf 12 appearance+8 (L12x8) | C, V5 |  |  |  |
| Leaf 15 appearance+0 (L15x0) | C, V5 |  |  |  |
| Leaf 15 appearance+4 (L15x4) | C, V5 | C, V5 |  |  |
| Leaf 15 appearance+8 (L15x8) | C, V5 | C, V5 |  |  |
| Leaf 18 appearance+0 (L18x0) |  | C, V5 | C, V5 |  |
| Leaf 18 appearance+4 (L18x4) |  | C, V5 | C, V5 |  |
| Leaf 18 appearance+8 (L18x8) |  | C, V5, V12 | C, V5, V12 |  |
| Ear emergence - 4 (Ear_4) |  | C, V5, V12 |  | C, V5, V12 |
| Ear emergence - 2 (Ear_2) |  |  |  | C, V5, V12 |
| Ear emergence (Earx0) |  |  |  | C, V5, V12 |
| Silk emergence (Silk) |  |  |  | C, V5, V12 |

**Supplemental Table 2.** Ground truth internode data with the number of observations for each combination of conditions. Column ‘Collar i+1 visible’ indicates if the collar of the leaf attached to the apical node of the internode was visible in its associated RGB image (TRUE), and thus whether the internode was relevant for the direct internode length estimation method (TRUE) or the indirect internode length estimation method (FALSE). Column ‘Number’ indicates the number of observations for the specific combination of conditions specified by the other columns. Column ‘Mean Δsheath (mm)’ indicates mean Δsheath values for different treatments and internodes. Column ‘SD’ indicates standard deviation for Mean Δsheath values.

| Internode | Treatment | Collar i+1 visible | Number | Mean Δsheath (mm) | SD |
| --- | --- | --- | --- | --- | --- |
| Int9 | Control | TRUE | 32 | -17.49 | 4.76 |
| Int12 | Control | TRUE | 27 | -12.69 | 10.73 |
| Int15 | Control | TRUE | 9 | -4.17 | 10.2 |
| Int9 | V12-Drought | TRUE | 0 |  |  |
| Int12 | V12-Drought | TRUE | 18 | -10.45 | 5.87 |
| Int15 | V12-Drought | TRUE | 7 | 4.49 | 6.44 |
| Int9 | V5-Drought | TRUE | 24 | -24.17 | 6.51 |
| Int12 | V5-Drought | TRUE | 26 | -7.66 | 6.9 |
| Int15 | V5-Drought | TRUE | 20 | 0.61 | 4.8 |
| Int9 | Control | FALSE | 32 |  |  |
| Int12 | Control | FALSE | 36 |  |  |
| Int15 | Control | FALSE | 30 |  |  |
| Int9 | V12-Drought | FALSE | 0 |  |  |
| Int12 | V12-Drought | FALSE | 0 |  |  |
| Int15 | V12-Drought | FALSE | 9 |  |  |
| Int9 | V5-Drought | FALSE | 32 |  |  |
| Int12 | V5-Drought | FALSE | 38 |  |  |
| Int15 | V5-Drought | FALSE | 30 |  |  |

**Supplemental Table 3.** Effects of V5-Drought on the growth traits of internodes 9, 12 and 15. Listed are the mean values per treatment are listed, the relative effect of the V5-Drought treatment on the trait compared to the Control treatment, the absolute effect and the Holm family-wise error rate (FWER)-adjusted *p*-value for statistical significance of the treatment effect.

| Mean elongation rate (mm/h) |  |  |  |  |  |
| --- | --- | --- | --- | --- | --- |
| Internode | Control | V5-Drought | Relative effect | Absolute effect | Adj. p |
| Int9 | 0.80 | 0.46 | -41.73% | -0.33 | 0.01953 |
| Int12 | 0.84 | 0.41 | -51.04% | -0.43 | 8.909*10<sup>-4</sup> |
| Int15 | 0.98 | 0.52 | -46.31% | -0.45 | 5.023*10<sup>-4</sup> |
| Elongation duration (days) |  |  |  |  |  |
| Internode | Control | V5-Drought | Relative effect | Absolute effect | Adj. p |
| Int9 | 4.5 | 6.3 | 40.94% | 1.8 | 0.02942 |
| Int12 | 5.2 | 7.5 | 44.14% | 2.3 | 0.002160 |
| Int15 | 5.1 | 7.2 | 41.21% | 2.1 | 0.006044 |
| Final internode length (mm) |  |  |  |  |  |
| Internode | Control | V5-Drought | Relative effect | Absolute effect | Adj. p |
| Int9 | 94 | 76 | -19.51% | -18 | 1 |
| Int12 | 116 | 76 | -34.35% | -40 | 0.03280 |
| Int15 | 132 | 95 | -28.16% | -37 | 0.05063 |

**Supplemental Table 4.** Leaf growth zones and elongation rates across leaf rank, treatment, and time points. Mean size of the leaf lamina division zone, lamina elongation zone, and their standard deviations (SD) per combination of leaf rank, treatment and time point are shown. Leaf elongation rate (LER) is derived from the fitted growth curves in Verbraeken et al. (2021) for each leaf rank, treatment and time point.

| Leaf rank | Treatment | Timing | sample number | Mean division zone size (mm) | SD division zone size (mm) | Mean elongation zone size (mm) | SD elongation zone size (mm) | LER (mm/h) |
| --- | --- | --- | --- | --- | --- | --- | --- | --- |
| L9 | Control | L9x0 | 3 | 24.37 | 3.45 | 58.83 | 7.77 | 3.00 |
| L9 | Control | L9x4 | 3 | 22.17 | 8.46 | 65.17 | 15.57 | 4.03 |
| L9 | Control | L9x8 | 1 | 23.00 | NA | 36.50 | NA | 3.26 |
| L9 | V5-Drought | L9x0 | 3 | 28.13 | 2.56 | 55.00 | 5.27 | 2.40 |
| L9 | V5-Drought | L9x4 | 3 | 21.43 | 4.84 | 57.50 | 3.61 | 2.89 |
| L9 | V5-Drought | L9x8 | 3 | 18.27 | 9.36 | 62.83 | 2.08 | 2.69 |
| L12 | Control | L12x0 | 5 | 31.50 | 3.34 | 52.30 | 5.89 | 2.56 |
| L12 | Control | L12x4 | 3 | 32.17 | 2.31 | 58.17 | 9.02 | 3.11 |
| L12 | Control | L12x8 | 3 | 25.50 | 3.12 | 50.17 | 9.87 | 2.89 |
| L12 | V5-Drought | L12x0 | 3 | 28.67 | 3.25 | 54.50 | 5.20 | 2.00 |
| L12 | V5-Drought | L12x4 | 3 | 29.00 | 2.50 | 51.50 | 2.65 | 2.25 |
| L12 | V5-Drought | L12x8 | 3 | 25.00 | 3.04 | 45.83 | 5.51 | 2.16 |
| L15 | Control | L15x0 | 3 | 33.67 | 2.84 | 56.17 | 2.52 | 2.04 |
| L15 | Control | L15x4 | 3 | 29.17 | 0.29 | 53.50 | 2.00 | 2.30 |
| L15 | Control | L15x8 | 3 | 25.83 | 2.89 | 41.17 | 7.51 | 2.15 |
| L15 | V5-Drought | L15x0 | 3 | 31.17 | 8.08 | 55.83 | 6.66 | 1.48 |
| L15 | V5-Drought | L15x4 | 3 | 30.33 | 4.04 | 49.17 | 1.53 | 1.57 |
| L15 | V5-Drought | L15x8 | 3 | 22.33 | 1.44 | 41.50 | 1.73 | 1.51 |
| L18 | Control | L18x0 | 2 | 26.25 | 2.47 | 49.00 | 0.71 | 1.19 |
| L18 | Control | L18x4 | 2 | 26.00 | 2.83 | 37.00 | 13.44 | 1.25 |
| L18 | Control | L18x8 | 2 | 9.50 | 5.66 | 50.00 | 10.61 | 1.21 |
| L18 | V5-Drought | L18x0 | 2 | 25.00 | 4.95 | 46.50 | 1.41 | 0.99 |
| L18 | V5-Drought | L18x4 | 2 | 15.75 | 3.89 | 42.50 | 2.83 | 1.17 |
| L18 | V5-Drought | L18x8 | 1 | 8.50 | NA | 52.50 | NA | 1.27 |

**Supplemental Table 5.** Internode growth zone size at different V-stages. Mean distance to the internode base for the top of the division zone and the top of the elongation zone, the size of the elongation zone and the lower (LowerCI) and upper (UpperCI) limits of the 95% confidence intervals for these traits are shown.

| V-Stage | internode<br>Length<br>(mm) | Division<br>zone<br>LowerCI<br>(mm) | Division<br>zone<br>(mm) | Division<br>zone<br>UpperCI<br>(mm) | Elongation<br>zone<br>LowerCI<br>(mm) | Elongation<br>zone (mm) | Elongation<br>zone<br>UpperCI<br>(mm) | Elongation<br>zone<br>length<br>LowerCI<br>(mm) | Elongation<br>zone<br>length<br>(mm) | Elongation<br>zone<br>length<br>UpperCI<br>(mm) |
| --- | --- | --- | --- | --- | --- | --- | --- | --- | --- | --- |
| 12 | 13.5 | 8.73 | 9.65 | 10.67 | 12.23 | 12.85 | 13.40 | 2.55 | 3.20 | 3.70 |
| 13 | 23.8 | 5.03 | 8.24 | 11.35 | 10.85 | 16.62 | 24.94 | 3.47 | 8.37 | 16.99 |
| 14 | 65.1 | 2.03 | 3.93 | 5.56 | 16.63 | 21.52 | 33.37 | 11.44 | 17.59 | 28.36 |
| 15 | 103.0 | 1.54 | 2.21 | 2.68 | 9.84 | 13.88 | 24.39 | 8.09 | 11.68 | 21.91 |

**Supplemental Table 6.**
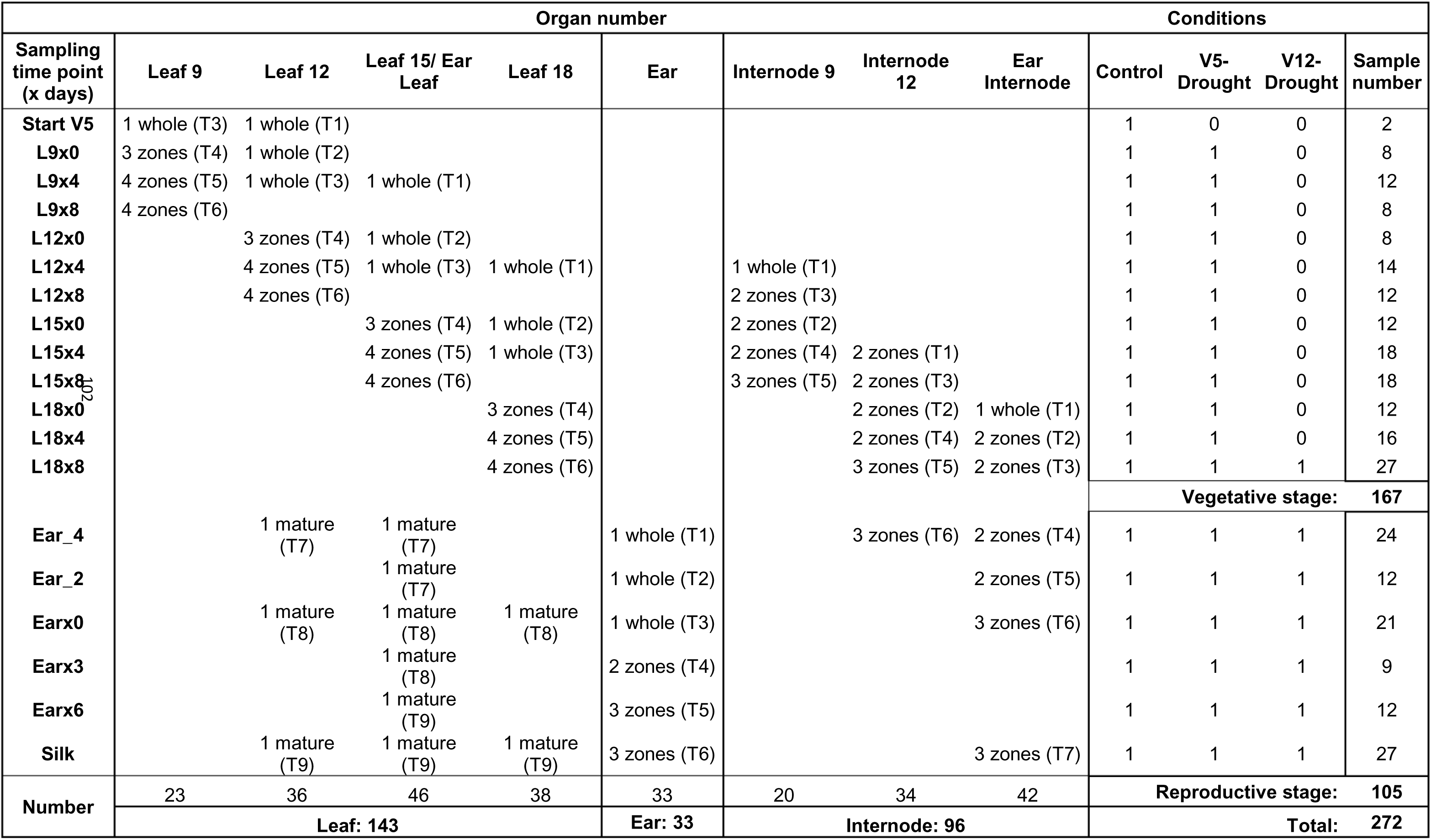
Overview of samples for transcriptome analysis.

**Supplemental Table 7.** Number of DEGs between different leaf zones in leaf 15 (L15).

| Zone<br>comparison | Time point |  |  |
| --- | --- | --- | --- |
|  | L15x0 | L15x4 | L15x8 |
| Elong vs Div | 6924 | 6316 | 6874 |
| Diff vs Div | 13272 | 12830 | 12596 |
| Diff vs Elong | 8487 | 7156 | 6774 |
| Mat vs Div |  | 15912 | 15627 |
| Mat vs Elong |  | 14481 | 14438 |
| Mat vs Diff |  | 12775 | 12483 |

**Supplemental Table 8.**
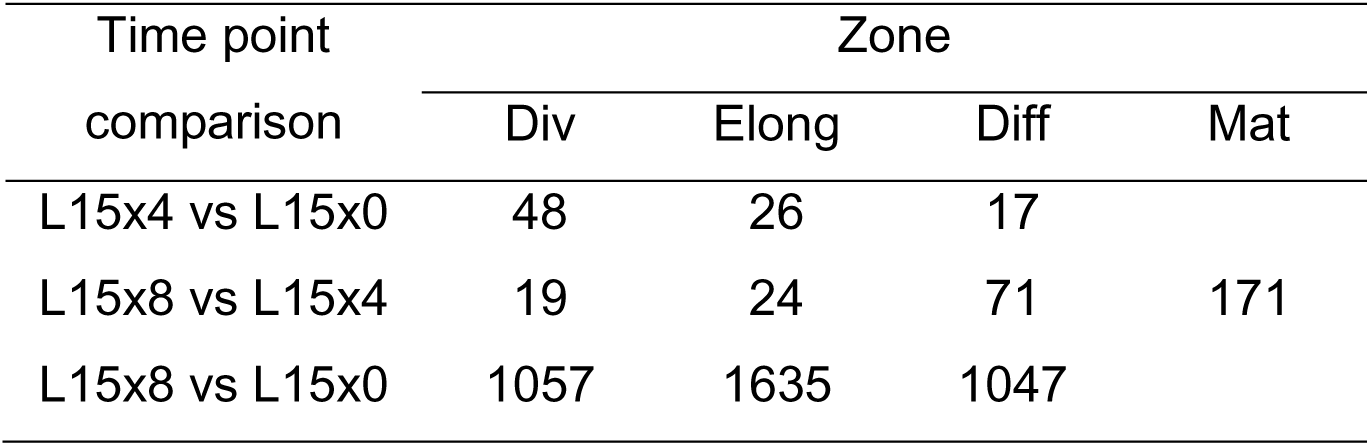
Number of DEGs between different sampling time points in leaf 15 (L15).

**Supplemental Table 9.** Number of DEGs between different sampling time points in internode 12 (Int12).

| Time point<br>comparison | Zone |  |  |
| --- | --- | --- | --- |
|  | Low | Mid | High |
| L18x0 vs L15x4 | 165 |  | 200 |
| L15x8 vs L15x4 | 487 |  | 2141 |
| L18x4 vs L15x4 | 3061 |  | 7493 |
| L18x8 vs L15x4 | 10526 |  | 12365 |
| Ear_4 vs L15x4 | 11689 |  | 12688 |
| L15x8 vs L18x0 | 3 |  | 1 |
| L18x4 vs L18x0 | 1692 |  | 5525 |
| L18x8 vs L18x0 | 10555 |  | 11470 |
| Ear_4 vs L18x0 | 11794 |  | 11653 |
| L18x4 vs L15x8 | 975 |  | 3675 |
| L18x8 vs L15x8 | 10325 |  | 10496 |
| Ear_4 vs L15x8 | 11847 |  | 10753 |
| L18x8 vs L18x4 | 8202 |  | 7064 |
| Ear_4 vs L18x4 | 10590 |  | 7935 |
| Ear_4 vs L18x8 | 910 | 220 | 159 |

**Supplemental Table 10.**
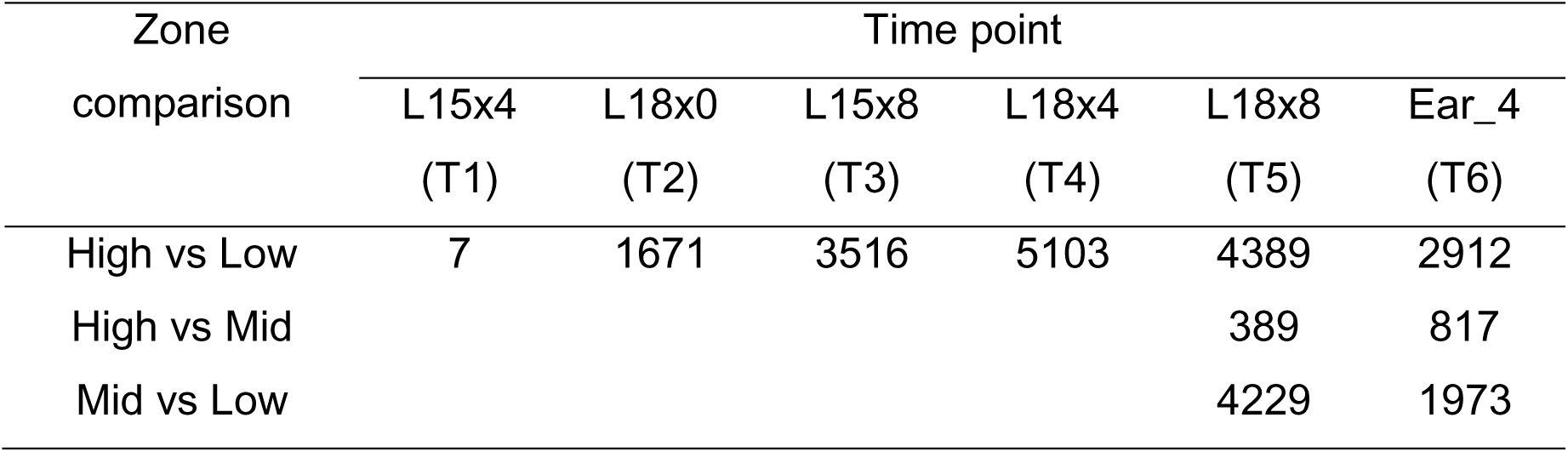
Number of DEGs between different internode zones in internode 12 (Int12).

| Zone<br>comparison | Time point |  |  |  |  |  |
| --- | --- | --- | --- | --- | --- | --- |
|  | L15x4<br>(T1) | L18x0<br>(T2) | L15x8<br>(T3) | L18x4<br>(T4) | L18x8<br>(T5) | Ear_4<br>(T6) |
| High vs Low | 7 | 1671 | 3516 | 5103 | 4389 | 2912 |
| High vs Mid |  |  |  |  | 389 | 817 |
| Mid vs Low |  |  |  |  | 4229 | 1973 |

**Supplemental Table 11.** Number of DEGs between different sampling time points in ear tissue.

| Time point<br>comparison | Zone |  |  |  |
| --- | --- | --- | --- | --- |
|  | whole | Low | Mid | High |
| Ear_2 vs Ear_4 | 5 |  |  |  |
| Ear_0 vs Ear_4 | 42 |  |  |  |
| Ear_0 vs Ear_2 | 6 |  |  |  |
| Earx6 vs Earx3 |  | 16 |  | 27 |
| Silk vs Earx3 |  | 6569 |  | 6224 |
| Silk vs Earx6 |  | 4829 | 4941 | 5053 |

**Supplemental Table 12.** Number of DEGs between different ear zones during ear development.

| Zone<br>comparison | Time point |  |  |
| --- | --- | --- | --- |
|  | Earx3 | Earx6 | Silk |
| High vs Low | 18 | 62 | 856 |
| High vs Mid |  | 19 | 975 |
| Mid vs Low |  | 2 | 3 |

## Supplemental Data Set

**Supplemental Data Set 1.** Ground truth internode lengths and Δcollar values.

**Supplemental Data Set 2.** Estimated internode length and analysis of internode growth traits.

**Supplemental Data Set 3.** Individual leaf epidermal cell length measurements and analysis of leaf growth zones.

**Supplemental Data Set 4.** Individual internode epidermal cell length measurements.

**Supplemental Data Set 5.** Summary of RNA sequencing libraries and mapping statistics.

**Supplemental Data Set 6.** DESeq2-normalized and vst-transformed expression levels of genes in different samples.

**Supplemental Data Set 7.** Spatiotemporal DEGs during leaf 15 development.

**Supplemental Data Set 8.** Spatiotemporal DEGs during internode 12 development.

**Supplemental Data Set 9.** Spatiotemporal DEGs during ear development.

**Supplemental Data Set 10.** *K*-means clusters of all expressed genes in control samples.

**Supplemental Data Set 11.** *K*-means clusters of all expressed genes in leaf samples.

**Supplemental Data Set 12.** *K*-means clusters of all expressed genes in internode samples.

**Supplemental Data Set 13.** Overlap genes between leaf clusters and internode clusters.

**Supplemental Data Set 14.** Organ-specific genes.

**Supplemental Data Set 15.** V5-Drought inducible DEGs during leaf development.

**Supplemental Data Set 16.** V5-Drought inducible DEGs during internode development.

**Supplemental Data Set 17.** V5-Drought inducible DEGs during ear development.

**Supplemental Data Set 18.** List of genes described in this study.

